# MALT1 is an E3 ligase catalyzing its own activation by monoubiquitination

**DOI:** 10.64898/2026.11.23.768581

**Authors:** Marcus J C Long, Manhui Luo, Carla Chiavi, Richard Cowen, Chantal Décaillet, Gareth Hall, Margot Thome

**Author notes:** Department of Chemistry, University of Oxford, Mansfield Road, Oxford, OX1 3TA, United Kingdom, and Department of Pharmacology, University of Oxford, Mansfield Road, Oxford, OX1 3TA, United Kingdom (Marcus J C Long). Correspondence should be addressed to M.T. or M.J.C.L. : Margot Thome, Department of Immunobiology, Faculty of Biology and Medicine, University of Lausanne, CH-1066 Epalinges, Switzerland, Marcus JC Long, Department of Chemistry, University of Oxford, Mansfield Road, Oxford, OX1 3TA, United Kingdom.

## Abstract

The protease mucosa-associated lymphoid tissue lymphoma translocation protein-1 (MALT1) is an important signaling component of the antigen receptor-induced activation of lymphocytes. Abnormal MALT1 activity is linked to emergence of several lymphoma subtypes. Humans and mice with inactivating MALT1 mutations, on the other hand, show defects in specific classes of immune cells and develop autoimmunity. MALT1 protease activity depends on its inducible K644-monoubiquitination. We have previously identified a negatively charged ubiquitin binding site in MALT1, located ∼30Å from K644, that is necessary for MALT1 monoubiquitination. However, the machinery delivering activated ubiquitin to MALT1 and transferring it to K644 remains unknown. Here we provide evidence that MALT1 is an E3-ligase catalyzing autoubiquitination via a cascade involving MALT1 residue C687, accepting ubiquitin from a ubiquitinated E2-enzyme, and transferring that active ubiquitin to the MALT1 monoubiquitination site, K644. We further identified MALT1 residue Y631, which lies between MALT1(C687) and MALT1(K644), as a gatekeeper of this process. Lymphocyte lines expressing MALT1(C687A) or MALT1(Y631A) mutants showed impaired or increased activation, respectively. Finally, we identified Ube2L3 as a candidate cysteine-specific E2-enzyme servicing MALT1. Overexpression of inactive Ube2L3 compromised MALT1 monoubiquitination and protease activity, indicating Ube2L3 inhibition is a potential means to dampen MALT1 ubiquitination. Collectively, the findings identify MALT1 as an E3 ligase capable of catalyzing its own activation through auto-ubiquitination, which may open new avenues for therapeutic regulation of MALT1 activity.

**Brief abstract:** Understanding key enzymatic steps in immune cell activation is crucial for our basic comprehension of the immune response and for developing new therapeutics for immunomodulation and treatment of haematological cancers. Here we uncover new insights into the monoubiquitination-dependent activation of MALT1, a key immune regulatory protease. These findings disclose new MALT1 functions and chemically actionable players in adaptive immunity and lymphomagenesis.

- MALT1 uses a gated thioester relay system involving MALT1(C687) (transthioesterase) and MALT1(Y631) (gate) to ubiquitinate its own residue K644
- This process involves the E2 enzyme Ube2L3 and likely other, similar E2 enzymes
- Impairment of this relay system suppresses MALT1 protease activity and cytokine production

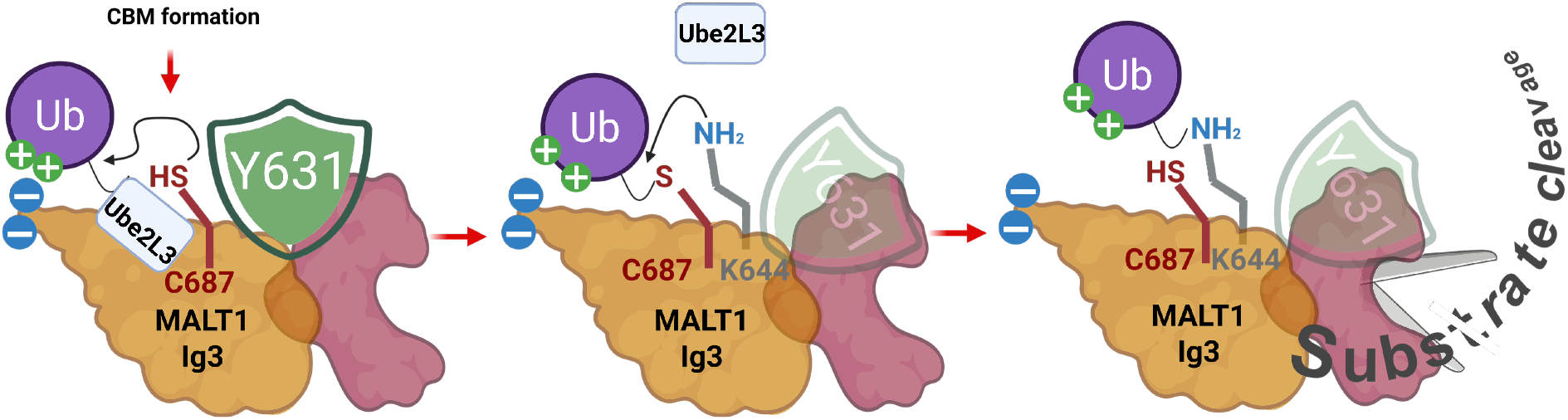

## Introduction

The activation of lymphocytes through antigen-induced T and B-cell receptor (T/BCR) signaling is crucial for correct functioning of the adaptive immune response. Defects in antigen-receptor signaling are linked to immunodeficiencies, whilst constitutive activation of these processes favors autoimmunity and development of lymphoid malignancies. Many of the intricacies of this pathway have been uncovered. In healthy T-and B-cells, triggering of the T/BCR induces recruitment of downstream kinases activating a core signalosome, consisting of the scaffold protein CARMA1 (also known as CARD11), the adaptor protein B-cell lymphoma-10, BCL10, and the protease mucosa-associated lymphoid tissue translocation protein-1, MALT1 (*1*). This ternary complex, known as “CBM”, scaffolds polyubiquitin chains, both K63-linked (formed predominantly by the ubiquitin ligase TNF receptor-associated factor 6, TRAF6) (*2*) and linear (formed exclusively by the linear ubiquitin chain assembly complex, LUBAC) (*3*). Each individual CBM component is essential for CBM’s scaffolding function, which controls activation of the transcription factors NF-κB1 and AP-1. Ubiquitin chains on CBM recruit the IκB kinase (IKK) complex inducing phosphorylation of IκB, the inhibitor of the transcription factor NF-κB1. IκBα is thus degraded, releasing NF-κB1 for nuclear entry and stimulation of gene expression. CBM also favors recruitment and activation of kinases promoting activation of AP-1, a heterodimer composed of proteins of the jun, fos and ATF families (*1*). Together, the NF-κB1 and AP-1 transcription factors induce expression of a battery of genes that promote lymphocyte activation and proliferation, spurring inflammatory immune processes.

Another consequence of CBM formation is the activation of the protease activity of MALT1, which requires MALT1 monoubiquitination on K644 (*4*). MALT1 protease activity greatly strengthens NF-κB1 and AP-1 signaling by cleaving negative regulators of these pathways. Key substrates include: the deubiquitinating enzymes (DUBs) A20 and CYLD, which remove polyubiquitin chains (*5, 6*); the NF-κB2 family member, RelB, which forms transcriptionally inactive complexes with NF-κB1 family members (*7*); and Regnase-1 and Roquin-1/2 that negatively regulate short-lived proinflammatory transcripts (*8, 9*). The crucial role for MALT1 protease activity in T/BCR signaling is clear from cultured lymphocyte models, such as the human T-cell line, Jurkat, or the B-cell line, BJAB, wherein MALT1 protease activity can be genetically or pharmacologically downregulated. MALT1 protease activity is also essential for the *in vitro* and *in vivo* growth of B-cell lymphoma models in which MALT1 is constitutively active due to oncogenic mutations in its upstream regulators, such as CARMA1 (*10*). Mice bearing inactivating MALT1 mutations show severe immune defects, which manifest in several specific subsets of B-and T-lymphocytes, NK and myeloid cells. The observed defective immune responses are combined with autoimmunity, resulting from a concurrent defect in the development of natural regulatory T (nTreg) cells (*11*) (*12–14*), which serve to keep the immune system in check.

In resting T-and B-cells, MALT1 forms an inactive heterodimer with BCL10 (*15, 16*). *In vitro*, MALT1 dimerization/oligomerization stimulates MALT1 protease activity (*5, 17, 18*). Naturally, MALT1-oligomerization can occur in activated lymphocytes through recruitment into the CBM complex, but in simple cell culture models, such as 293T cells, MALT1 can be artificially oligomerized and activated through co-transfection with BCL10 or oncogenic CARMA1 mutants (*5, 18, 19*). Contemporaneously with CBM formation, MALT1 is monoubiquitinated at a specific residue, K644, located at the surface of its Ig3 domain (*4*). This step is crucial for upregulation of MALT1 protease activity, most likely by inducing a conformational change that is transmitted from the Ig3 to the adjacent protease domain (*20*). Constitutive MALT1 monoubiquitination is also a hallmark of CBM-driven lymphomas (*4*). MALT1(K644R) that cannot undergo monoubiquitination, is protease deactivated but competent for scaffolding. Its expression in CBM-driven lymphoma models reduces cell viability, and cultured immune cells expressing this mutant show reduced cytokine production upon stimulation. Conversely, appendage of ubiquitin to the C-terminus of MALT1 (MALT1-Ub), mimicking the CBM-induced K644-monoubiquitinated state, stimulates protease activity *in vitro* and in cells in the absence of other activation stimuli (*4*).

Therefore, MALT1 monoubiquitination is crucial for MALT1 protease activity; nevertheless, it remains unclear how MALT1 monoubiquitination occurs and which enzymes are involved in ubiquitin conjugation. We have previously defined a negatively charged patch (comprising E696 and D697) in the MALT1 Ig3 domain that binds free ubiquitin (*20*). This site is referred to here as “UBS1”. Mutations in this region, notably MALT1(E696,D697), suppress MALT1 monubiquitination in cells, dampening MALT1 protease activity. However, in MALT1-Ub fusion constructs, UBS1 domain inactivation no longer suppresses MALT1 protease activity (*20*). Thus, this region is only important for initial CBM-mediated ubiquitin recruitment and conjugation of ubiquitin to MALT1(K644), but not for the subsequent, ubiquitin-dependent MALT1 activation step.

Classically, ubiquitination requires activated ubiquitin (formed by a ubiquitin E1-enzyme; two exist in humans) (*21*), and a ubiquitin conjugating enzyme (E2; there are ∼40 in humans) (*22*), that accepts activated ubiquitin from the E1. The final step in the cascade involves a ligase (E3; there are ∼700 in humans). The majority of ligases are really interesting new gene (RING)-ligases that bridge the activated E2 and the target, facilitating ubiquitin transfer to an acceptor lysine directly from the E2 (*23*). A small subset, comprising Homologous to the E6AP Carboxyl Terminus (HECT) and RING in between RING (RBR) ligases, ∼30 E3s (∼5 %), have intrinsic capacity to form ubiquitin thioesters and transfer ubiquitin to substrates themselves (*24*). Identifying specific players in ubiquitin cascades and their precise biochemical functions are both crucial for biological interrogations. Such information provides a means to intervene pharmaceutically in ubiquitin-dependent pathways, distinct from targeting players directly involved in a specific pathway.

There have been many efforts to test MALT1 protease inhibitors to combat inflammatory diseases and cancers. A major drawback of these efforts has been that prolonged treatment of animals with MALT1 inhibitors impairs the development of Tregs (*25, 26*). Animals treated with MALT1 inhibitors develop autoimmunity, reminiscent of observations in MALT1 catalytically deficient mutants in mice (*11–13, 26*). However, MALT1 protease inhibition indicates differently in diseased versus healthy animals (*27*), and there is evidence that induced MALT1 protease inactivation is better tolerated than germline inactivation (*28*). Recent studies have additionally suggested that acute MALT1 inhibition can convert immunosuppressive Tregs into pro-inflammatory effector cells in the context of the tumor microenvironment of solid cancers; this can strengthen anti-tumor immune responses (*29, 30*). Thus, gaining a better understanding of how MALT1 is activated and identifying other targets to modulate MALT1 protease activity is of fundamental importance to human health.

One of the outstanding unknowns surrounding MALT1 protease activation is how the key regulatory lysine residue MALT1(K644) becomes ubiquitinated. This must start from ubiquitin, or more likely an E2-ubiquitin complex, binding UBS1, for subsequent conjugation onto K644. Although UBS1 and K644 are located on the surface of the MALT1 Ig3 domain, due to the 30 Å distance between these sites (*20*), direct transfer of ubiquitin is unlikely. Here we propose a ubiquitin relay system within MALT1 that transfers ubiquitin from its binding site at UBS1 to K644. This relay system depends on transthioesterification, using a specific surface residue of MALT1, C687, followed by transamidation to K644. Mutants affecting this pathway either positively or negatively impact MALT1-ubiquitination and protease activity. Gene silencing of a putative E2 involved in this pathway, and novel MALT1 interactor, Ube2L3, diminishes MALT1-ubiquitination. These findings identify MALT1 as an active E3 ligase inducing its own activation through Ube2L3-mediated monoubiquitination.

## Results

### C687 is a crucial residue involved in MALT1 monoubiquitination in HEK293T cells

Examination of X-ray structures and AlphaFold models of MALT1 showed that the ubiquitin binding site UBS1 (comprising residues E696 and D697; blue) and the ubiquitin conjugation site K644 (grey) lie on the same face of the MALT1 Ig3 domain, but are separated by approximately 30 Å (**Figure 1A**). This arrangement is not ideal for direct K644 ubiquitination, as UBS1 points away from K644. We however noticed a surface-accessible cysteine residue, C687 (red), positioned between UBS1 and K644 and substantially closer to K644 (**Figure 1A**). C687 is also highly conserved across mammals and fish despite sequence variation in the surrounding region (**Figure 1B**). We therefore hypothesized that C687 acts as an intermediary by accepting ubiquitin from a UBS1-bound ubiquitin-E2 thioester, forming a transient C687-Ub thioester. This putative transthioesterification step is common in biochemical processes and is overall energy neutral, rapid, and reversible (*32*). This intermediate could then facilitate ubiquitin transfer to K644 (**Figure 1C**), a pathway expected to be kinetically favoured over direct transamidation (*33*)(*34*). Structural modelling showed that conformational flexibility could bring the C687 thioester within ∼9 Å of K644 (**Appendix Figure S1A,B**), whereas K644 remained ∼15 Å from UBS1 (**Appendix Figure S1C**). Consistent with this, the region around C687 exhibited high flexibility in structural models and the greatest conformational variation between apo and ligand-bound MALT1 structures (*35*)(*36*) (**Appendix Figure S1C-E**). Moreover, the region proximal to C687 showed the strongest position variation when comparing ligand-docked and apo structures of MALT1 (**Appendix Figure S1E**). Together, these findings suggest that activated Ubiquitin is more likely transferred from UBS1 to K644 via a C687-ubiquitin intermediate than by direct proximity.

**Figure 1.**
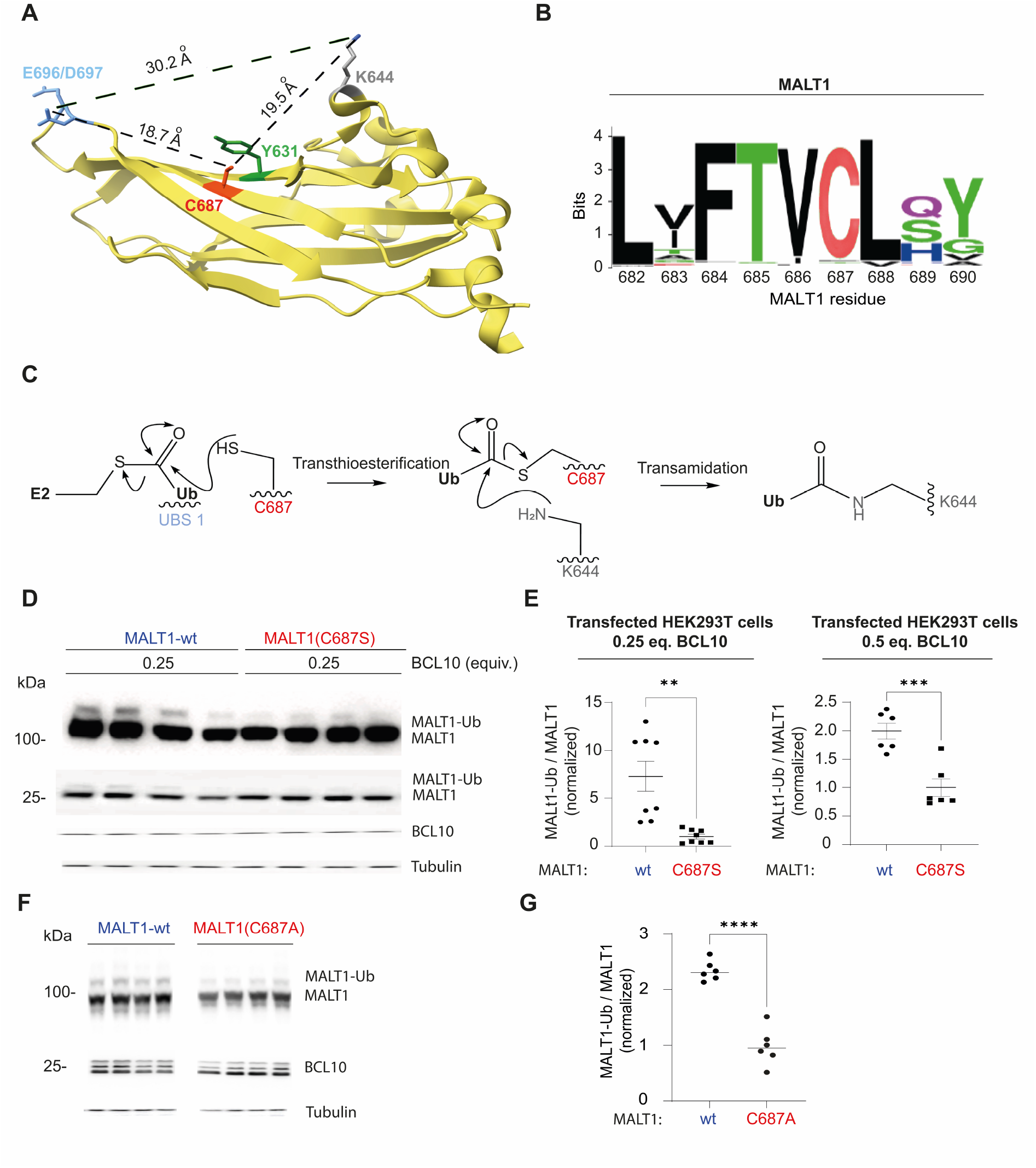
C687 plays a crucial role in MALT1 monoubiquitination. (**A**) X-ray structure of MALT1 6F7I, showing the Ig3 domain with UBS1 in periwinkle, and residues described in this work, C687 (red), Y631 (green), K644 (grey with amine function in dark blue), as well as important distances; the conformation of Y631 overhanding C687 represents the “gate closed” conformation. (**B**) Sequence logo for C687, illustrating its conservation across mammals and fish. (**C**) Proposed mechanism for ubiquitination of K644 via a C687 ubiquitin-thioester intermediate. Wavy line indicates specific sites: UBS1 (periwinkle); C687 (red); K644 (grey) located at the surface of the Ig3 domain. (**D**) HEK293T cells were transfected as shown with either wt-MALT1 or MALT1(C687S) and treated with MLT-748 for 24 h. After this time samples were analyzed by western blot as indicated. (**E**) Quantification of data in **D** (n=8,6). (**F**) HEK293T cells were transfected as shown with either wt-MALT1 or MALT1(C687A) and treated with MLT-748 for 24 h. After this time samples were analyzed by western blot as indicated. (**G**) Quantification of data in **F** (n=6). Data are shown as four independent biological replicates. Data were analyzed by two-tailed, unpaired *t*-test; *0.05>p≥0.01; **0.01>p≥0.001; ***0.001>p≥0.0001; ****p<0.0001.

To test this hypothesis, we investigated how MALT1 ubiquitination was changed upon mutating C687 into Serine (C687S). For this we examined ubiquitination of MALT1-wt and MALT1(C687S) in HEK293T cells cotransfected with differing amounts of BCL10, an experimental system that promotes MALT1 activation and MALT1 ubiquitination. Cells were also treated with the MALT1 inhibitor, MLT-748, as MALT1 inhibition stabilizes MALT1 monoubiquitination (*4*). We found that in presence of low amounts of BCL10, cells transfected with MALT1(C687S) showed decreased MALT1-monoubiquitination relative to cells transfected with MALT1-wt (**Figure 1D and E**). This effect was less pronounced when the amount of BCL10 was increased (**Figure 1D and E**). Similar observations were made for MALT1(C687A) (**Figure 1F and G**). Overall, these findings suggest that C687 aids the transfer of ubiquitin from a UBS1-bound ubiquitin-thioester to K644, most likely by facilitating transthioesterification.

### Y631 acts as a gatekeeper for ubiquitin transfer during MALT1-monoubiquitination in HEK293T cells

When considering potential mechanisms of regulation of ubiquitin transfer from C687 to K644, we noticed a tyrosine residue, Y631, positioned less than 4Å above C687, along the trajectory between C687 and K644. We reasoned that Y631 may regulate ubiquitin transfer from C687 to K644 (**Figure 2A**). Indeed, in two PDB structures (6H4A and 4L1R) (*37*), the phenol sidechain within Y631 undergoes a rotation to a conformation making C687 much more accessible to K644, confirming the feasibility of our proposed mechanism (**Figure S1A**, white versus green phenol rings). Y631 is not particularly well conserved between fish and human and lies in a region of low overall conservation (**Figure 2B**). Nonetheless, Y631 is well conserved in mammals, and we thus investigated the importance of Y631 in MALT1 monoubiquitination further. Proceeding with the hypothesis that Y631 is a gatekeeper of UBS1-K644 ubiquitin transfer, we examined the effect of a size-reducing Y631A mutation on MALT1 monoubiquitination in HEK293T cells. We therefore compared MALT1 monoubiquitination for the protease-inactive form of MALT1 (C464A) and the double mutant MALT1(C464A,Y631A) (*4*), and found that MALT1 monoubiquitination was dramatically increased in the latter (**Figure 2C and D**). Similar results were found in HEK293T cells transfected with MALT1-wt or MALT1(Y631A) and treated with the MALT1 inhibitor MLT-748 (**Figure S1B and C**). In this system we also tested the effect of mutation of the critical tyrosine residue Y631 into phenylalanine, MALT1(Y631F), that did not show elevation of ubiquitination (**Figure S1B and C**). Furthermore, introduction of the C687S mutation into the Y631A mutant suppressed the increased monoubiquitination of the Y631A mutant (**Figure 2E and F**). This result implies that the Y631A mutation increases MALT1 monoubiquitination by augmenting a pathway involving C687.

**Figure 2.**
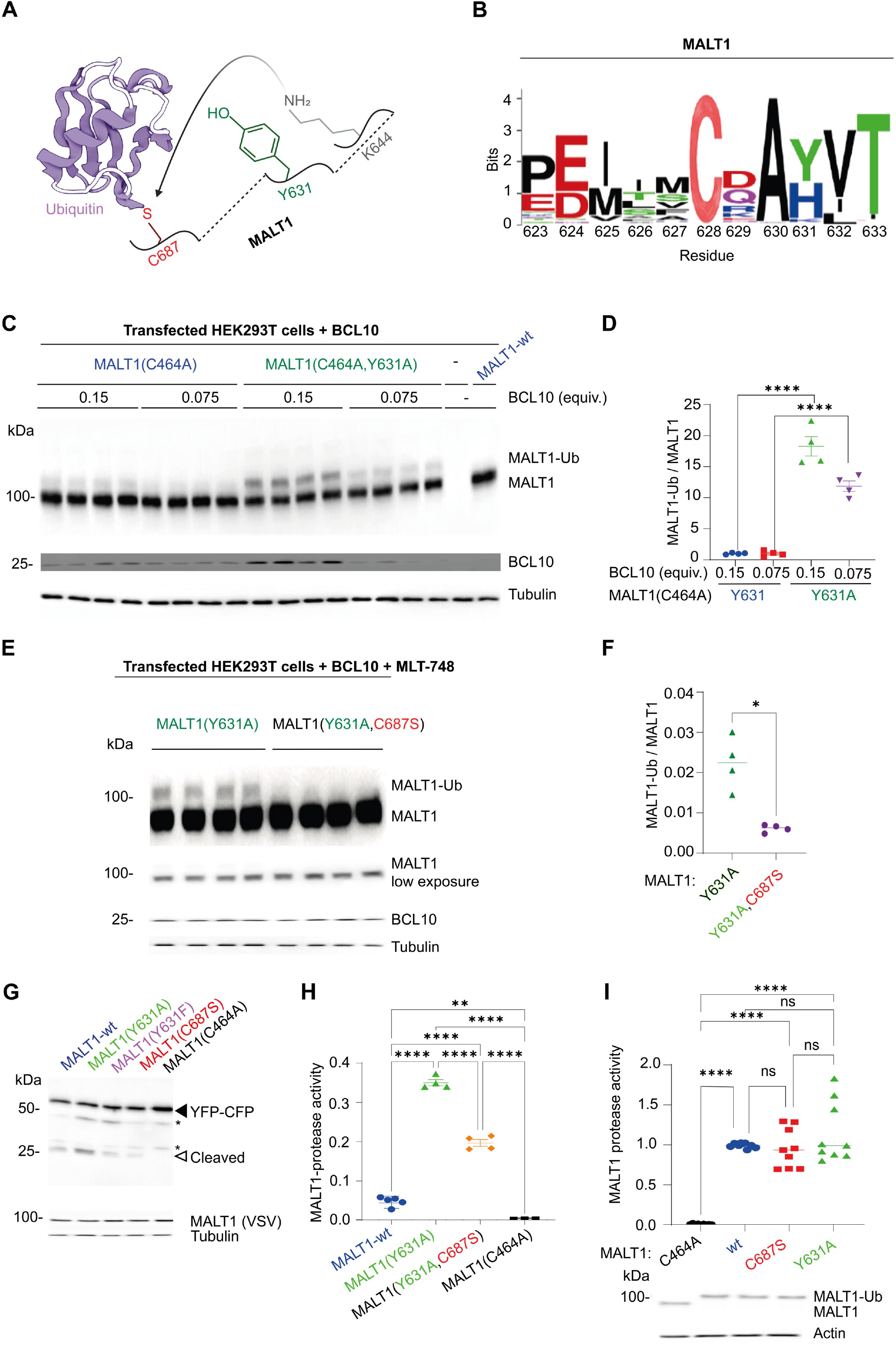
Y631 acts as a gatekeeper residue during MALT1 monoubiquitination via C687. (**A**) Schematic for Y631 blocking access of K644 to Ub-conjugated C687. (**B**) Sequence logo around Y631, showing moderate to strong conservation of Y or H at this position. (**C**) HEK293T cells were transfected as shown with either MALT1(C464A) or MALT1(C464A,Y631A) (quadruplicate samples from independently transfected, and processed wells are shown for each condition) and analyzed by western blot as indicated. (**D**) Quantification of data in **C** (n=4). (**E**) HEK293T cells were transfected as described and treated with MLT-748 for 24 h (quadruplicate samples from independently transfected, and processed wells are shown for each condition). These samples were independently analyzed by western blot as indicated. (**F**) Quantification of data in **E** (n=4). All data are shown as four independent biological replicates. (**G**) HEK293T cells were transfected with the stated MALT1 constructs and YFP-LVSR-CFP and analyzed by western blot for GFP, VSV, and Tubulin. (**H**) HEK293T cells were transfected with the stated MALT1 constructs and YFP-LVSR-CFP and MALT1 protease activity was measured by FRET. (**I**) HEK293T cells were transfected with the stated MALT1 constructs (mostly bearing C-terminal ubiquitin) and YFP-LVSR-CFP and MALT1 protease activity was measured by FRET. Closed arrows indicate full length protein; open arrows indicate cleaved proteins. □ denotes non-specific band. Data were analyzed by one-way ANOVA followed by Tukey’s test or t-test; *0.05>p≥0.01; **0.01>p≥0.001; ***0.001>p≥0.0001; ****p<0.0001.

We previously identified an Ig3 domain mutant MALT1(Y657A) that increases the intrinsic protease activity of MALT1 and causes pronounced changes (>0.1 ppm) in the 2D TROSY-HSQC ^1^H/^15^N NMR spectrum of homogeneously ^15^N labelled MALT1 (*20*), particularly at the Ig3-protease interface and near the active site. In contrast, MALT1(Y631A) caused only minimal structural perturbations, limited to residues surrounding Y631. These observations are consistent with altered shielding from replacement of the tyrosine aromatic ring by alanine (**Appendix Figure S2A-C**). Importantly, these perturbations mapped to residues near C687, supporting their spatial proximity. This result indicates that the MALT1(Y631A) phenotype is unlikely to result from gross structural changes, but rather reflects a functional impairment of the Y631 gatekeeper residue.

### C687 and Y631 are important for MALT1 protease activity in HEK293T cells

MALT1-monoubiquitination is required for MALT1 protease activity (*20*). We thus hypothesized that MALT1(C687S) and MALT1(C687A) would display reduced protease activity whereas MALT1(Y631A) would have augmented activity. To test this hypothesis, HEK293T cells were transfected with a previously described FRET-based MALT1 protease activity reporter (*20*). This construct expresses a YFP-CFP fusion bearing LVSR, an ideal MALT1 substrate sequence, in the linker region between the two fluorescent proteins. Assessing cleavage of the reporter by western blot or measuring the ratio of FRET negative : FRET positive cells within a population of transfected cells thus gives a readout for MALT1 protease activity. Overexpression of MALT1-wt induced cleavage of the YFP-CFP reporter, in contrast to a previously described catalytically inactive MALT1(C464A)-mutant (**Figure 2G**) (*5*). Consistent with our previous data for MALT1-monoubiquitination, Y631A transfected cells (but not a Y631F mutant) showed elevated MALT1 protease activity, whereas MALT1(C687S)-transfected cells showed impaired MALT1 protease activity (**Figure 2G, Figure S1D and E**). MALT1(C687A) showed similarly reduced activity (**Figure S1F**).

Revisiting our alignment above (**Figure 2B**), we noticed that in some species, the prevailing residue in the 631 position is histidine. In human MALT1, mutating tyrosine to histidine (Y631H) had minimal effect on MALT1 protease activity in HEK293T cells (**Figure S1G**). Thus, tyrosine and histidine in position 631 are functionally analogous, explaining variability at this residue. Finally, we assessed the effect of combined mutation of Y631 and C687. Consistent with the MALT1-monoubiquitination data above (**Figure 1D**, **2G**), cells transfected with MALT1(Y631A,C687S) showed significantly less MALT1 protease activity than those transfected with MALT1(Y631A) (**Figure 2H**). Thus, Y631 and C687 regulate MALT1 ubiquitination and protease activity in opposing senses in transfected HEK293T cells.

Altogether, these data fit with a model in which C687 aids the transfer of ubiquitin from UBS1 to K644, and Y631 (or another polar aromatic residue such as histidine in the same position) gates transfer. Consistent with this notion, MALT1-wt and MALT1(Y631H) were more active than MALT1(Y631F). These findings may be linked to an interaction between the carboxylate side chain within D629 and the polar proton within Y/H631 in the open conformation, although this is relatively long range (5 Å) in the PDB structures (**Figgure S1A**). Regardless, mutating Y631 to a smaller residue, such as alanine, forms a permanently open state that augments ubiquitination by improving the rate of transfer of ubiquitin to C687 and/or the rate of transamidation. Crucially, the Y631A mutation did not significantly affect the structure of the Ig3 domain (**Appendix Figure S2A-C**). Consistent with these structural data, C-terminal ubiquitin fusion constructs of MALT1-wt, MALT1(Y631A), MALT1(C687A), which mimic the constitutively monoubiquitinated state and no longer require K644 ubiquitination to be active (*4*), had comparable MALT1 protease activity in transfected HEK293T cells (**Figure 2I**). Thus, C687 and Y631 are important residues controlling MALT1 protease activation through monoubiquitination.

### C687 positively regulates MALT1 protease activity in Jurkat T cells

We progressed to examine how mutation of C687 affects MALT1 function in Jurkat T cells. These are a commonly used model of naïve T-cells that have no basal NF-κB1 or MALT1 protease activity. In this cell line, MALT1 protease activity can be stimulated by combined treatment with phorbol myristate acetate and the calcium ionophore, ionomycin, (PI) that mimics strong TCR engagement, or by a combination of agonistic anti-CD3 and anti-CD28 antibodies. In two different MALT1-silenced cell lines derived from different MALT1-targeting gRNAs, we re-expressed MALT1-wt and MALT1(C687A). Upon MALT1 protease inhibition and PI-stimulation, MALT1(C687A)-expressing cells showed reduced MALT1-monoubiquitination (**Figure 3A and B**) and reduced cleavage of kinetically slow MALT1 substrates, such as HOIL (**Figure 3C, Figure S2A and B**) and CYLD (**Figure S2A and C**). Roquin-1 and Regnase-1, intrinsically better MALT1 substrates due to them having more efficient cleavage sites, were not significantly affected, consistent with these substrates being less dependent on optimal activation of MALT1 (**Figure 3C, Figure S2A,D and E**)(*17*). MALT1 scaffolding function, that is not linked to MALT1 monoubiquitination and is monitored by assessing IκB phosphorylation, was also unaffected (**Figure 3C, Figure S2A and F**). Nonetheless, there was significantly reduced IL-2 secretion in MALT1(C687A) expressing Jurkat lines relative to MALT1-wt expressing lines stimulated with PI or agonistic cross-linked anti-CD3/CD28 antibodies, especially when suboptimal stimulation with low antibody concentrations was applied (**Figure 3D and E**). Similar results were observed in Raji B cells, in which the C687 mutation also reduced MALT1 monoubiquitination without affecting MALT1 scaffold function (**Figure 3F and G**). Ubiquitination impairment also conferred reduced IL-8 production in PI-stimulated Raji B cells (**Figure 3H**). We thus conclude that the main manifestation of MALT1(C687A) mutation in lymphocytes is a reduced MALT1-monoubiquitination, with consequently reduced substrate cleavage and cytokine responses upon cellular activation.

**Figure 3.**
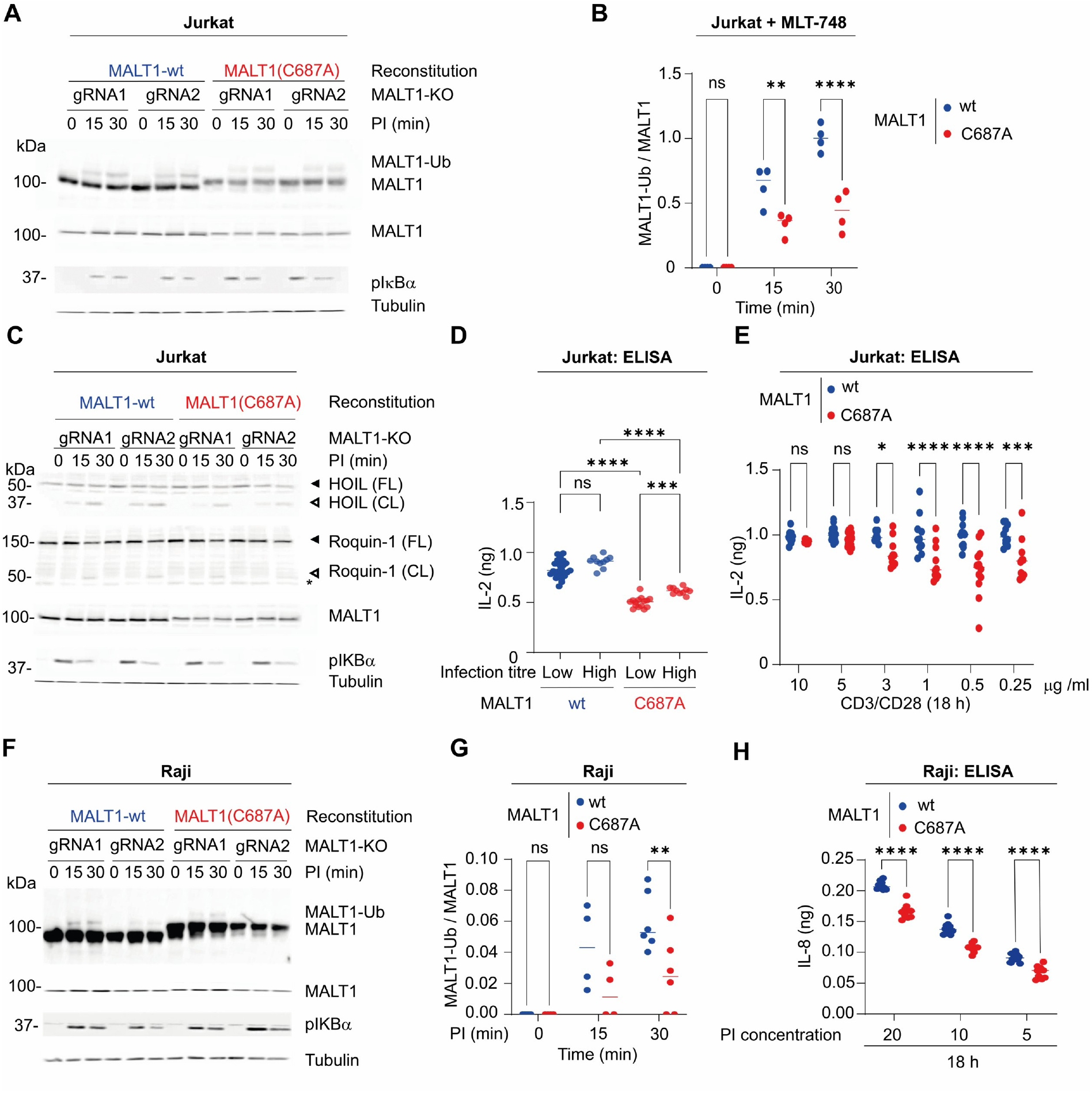
C687 regulates MALT1 protease activity in Jurkat and Raji cells. (**A**) MALT1-silenced Jurkat cells were infected with the stated MALT1 constructs and PI-stimulated (20 ng/ml PMA, and 1.00 μM ionomycin) as described. Lysates were analyzed by western blotting as indicated. (**B**) Quantitation of data in **A** (n=4). (**C**) The indicated Jurkat cell lines were PI-stimulated (20 ng/ml PMA, and 1.00 μM ionomycin) as described and analyzed by western blotting with the stated antibodies. (**D**) The stated Jurkat cell lines, transduced with low or high amounts of the indicated MALT1 expression constructs, were PI-stimulated (20 ng/ml PMA, and 1.00 μM ionomycin) for 18 h then IL-2 secretion was measured by IL-2 ELISA. (**E**) The stated Jurkat cell lines were stimulated with various concentrations of CD3/CD28 for 18 h then IL-2 secretion was measured by IL-2 ELISA. (**F**) MALT1-silenced Raji cells were infected with the stated MALT1 constructs and PI-stimulated (20 ng/ml PMA, and 1.00 μM ionomycin) for the indicated times. Lysates were analyzed by western blotting with the stated antibodies. (**G**) Quantitation of data in **F** (n=4,n=6). (**H**) Lines from **F** were stimulated with varying concentrations of PI (**20**: 20 ng/ml PMA, and 1.00 μM ionomycin; **10**: 10 ng/ml PMA, and 0.50 μM ionomycin; **5**: 5 ng/ml PMA, and 0.25 μM ionomycin) for 16 h then supernatants were analyzed by IL-8 ELISA (n=8). Tubulin was used as a loading control. CL – cleaved; FL – full length. Closed arrows indicate full length protein; open arrows indicate cleaved proteins. □ denotes non-specific band. Data were analyzed by either two-way ANOVA followed by Dunnett’s test or by one-way ANOVA followed by Tukey’s test; *0.05>p≥0.01; **0.01>p≥0.001; ***0.001>p≥0.0001; ****p<0.0001.

### Y631 negatively regulates MALT1 protease activity in Jurkat T-cells, via C687

Next, we examined the importance of Y631 and C687 in jointly regulating MALT1-dependent substrate cleavage in Jurkat T-cells. To assess the effect of Y631 and C687 mutations, MALT1-deficient Jurkat cells were reconstituted with either MALT1-wt, MALT1(Y631A), or MALT1(Y631A,C687S) constructs. MALT1(Y631A)-expressing cells showed elevated monoubiquitination relative to MALT1-wt expressing cells, as expected, whilst MALT1(Y631A,C687S) showed reduced MALT1-monoubiquitination relative to MALT1(Y631A)-expressing cells (**Figure 4A,B**). We next compared MALT1 protease activity in MALT1-wt and MALT1(Y631A) expressing cells by assessing cleavage of HOIL and CYLD, two suboptimal and kinetically slow substrates. MALT1(Y631A) showed significantly faster and stronger cleavage of HOIL and CylD than MALT1-wt (**Figure 4C,D,E**). Stimulation-induced IκBα phosphorylation, which relies on MALT1 scaffolding function and is independent of MALT1 monoubiquitination, was unaffected by the Y631 mutation (**Figure 4C**). Thus, the gatekeeper residue Y631 negatively regulates MALT1 monoubiquitination and protease activity, via C687, in activated T cells.

**Figure 4.**
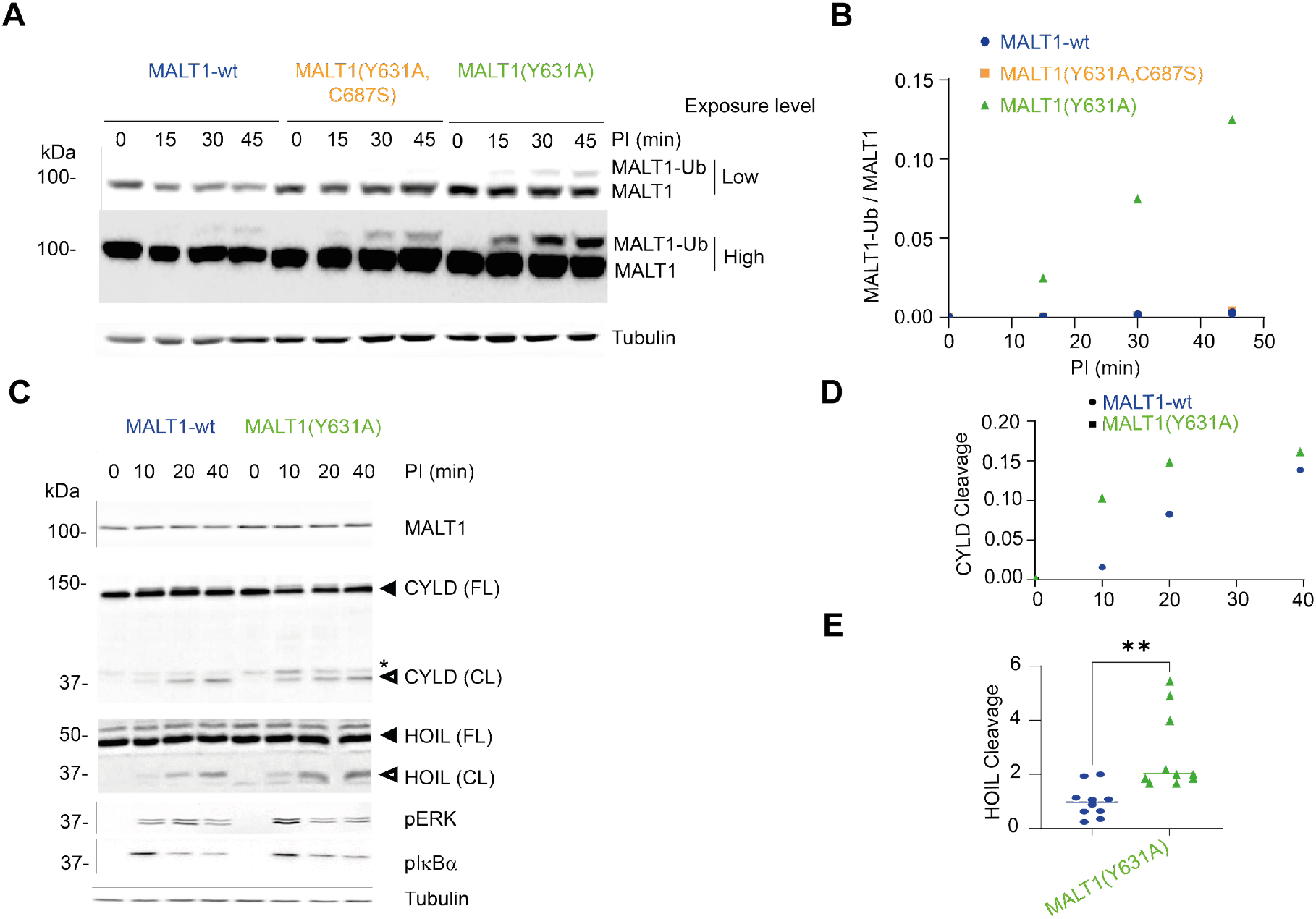
MALT1(Y631A) shows elevated monoubiquitination and protease activity in Jurkat T cells. (**A**) MALT1-deficient Jurkat T cells were infected with viruses delivering the stated constructs. These cells were sorted based on GFP that was co-expressed with MALT1 under an IRES system. Cells were pretreated with the MALT1 inhibitor MLT-748 for 3 h, then PI-stimulated (20 ng/ml PMA, and 1.00 μM ionomycin) for the stated times, and analyzed using western blotting as indicated. (**B**) Quantitation of data in **A**. (**C**) MALT1-deficient Jurkat cells expressing the stated constructs were PI-stimulated (20 ng/ml PMA, and 1.00 μM ionomycin) for the stated times, and then analyzed by western blotting as indicated. Two ladder lanes were removed from the HOIL cleavage panel for clarity (last and 2^nd^ to last lanes). (**D**) Quantification of CYLD cleavage data in **C**. (**E**) HOIL cleavage (for which cleavage in MALT1-wt expressing cells was set to 1 for each timepoint) was quantified in the stated lines (n=10). Tubulin was used as a loading control. CL – cleaved; FL – full length. Closed arrows indicate full length protein; open arrows indicate cleaved proteins. □ denotes non-specific band. Data were analyzed by a two-tailed, unpaired *t*-test; **0.01>p≥0.001.

### MALT1 ubiquitination is unaffected by various E3 ligase inhibitors and MALT1 can be ubiquitinated *in vitro* in the presence of activated E2 conjugating enzymes

If MALT1 can accept activated ubiquitin via C687 and transfer it to K644, the most likely candidates for this process are E2 conjugating enzymes. To test this hypothesis, we first examined if inhibition of different E3 ligases affected MALT1 protease activity. We used HOIPIN-8, an inactivator of the LUBAC E3 ligase HOIP that suppresses linear ubiquitination, and the NEDDylation inhibitor, pevonedistat, that inhibits the largest body of E3 ligases, Cullin-RINGs (of which ∼250 exist in humans) (*38*). These E3 inhibitors did not affect MALT1 protease activity and by extension MALT1-monoubiquitination (**Figure S3A**). Treatment with TAK-243, that inhibits E1-mediated activation and shuts down all ubiquitination in cells, was used as a positive control, testifying to the importance of MALT1-monoubiquitination for protease activity (**Figure S3A**). The IKK inhibitor MRT67307, on the other hand, did not impact MALT1-monoubiquitination. We interpreted these data to indicate that MALT1 is likely ubiquitinated by (an) activated E2-ubiquitin conjugate(s), whose activities are only affected by TAK-243, but not by a panel of inhibitors of E3 ligases.

To investigate if MALT1 can be recognized by E2s in the absence of a defined E3, we expressed MALT1, the E1 activating enzyme Ube1, and several E2 conjugating enzymes (UBC13, Ube2D3, and Ube2L3) recombinantly in *E coli* for subsequent purification (illustrated in **Figure S3B** for Ube2L3). We note that under these conditions it is very unlikely that MALT1 monoubiquitination can be trapped because the complex regulation of the CBM complex cannot be recapitulated easily in a tube. However, this assay can show sufficiency of E2s for MALT1 ubiquitination. When recombinant MALT1 was mixed individually with the activated E1, ubiquitin, and the different E2 enzymes in the presence of ATP, there was time-dependent ubiquitination of MALT1, including formation of a band likely corresponding to monoubiquitinated MALT1 and a large amount of polyubiquitinated MALT1 *in vitro* (**Figure S3C**). We conclude that MALT1 has indeed the capacity to use E2-ubiquitin conjugates for its own ubiquitination, to form diverse ubiquitinated species, consistent with its ability to bind free ubiquitin and accept ubiquitin directly from E2 enzymes.

### Ube2L3 is an E2-conjugating enzyme important in MALT1 K644 monoubiquitination in Jurkat T-cells and Raji B-cells

Our attention then turned to identifying the putative E2(s) responsible for creating MALT1-Ubiquitin by proximity profiling, using an ultraID-MALT1 construct expressed in MALT1-silenced Jurkat T cells. As a comparison, we used an ultraID-P2A-MALT1 construct that expresses the same ultraID and MALT1 proteins, but separated by virtue of ribosomal skipping (**Figure 5A**). Data obtained with this ultraID-P2A-MALT1 construct account for non-specific biotin labelling by ultraID (*39*), in a background that is identical to ultraID-MALT1. Transduced cells expressed the respective MALT1 constructs at close to endogenous levels (**Appendix Figure S3A**). Functional enrichment analysis of the ∼250 hits specific to ultraID-MALT1 showed several terms related to NF-κB (FDR 2.6×10^-4^) and downstream TCR signaling (FDR 3.4×10^-5^). In addition to known MALT1 interactors, BCL10 and TRAF6, and 17 known or predicted MALT1 substrates, we uncovered a single E2 that was selectively enriched in ultraID-MALT1, Ube2L3 (**Figure 5A**, for full data sets, please see: **PXD069479** and **Table S1**).

**Figure 5.**
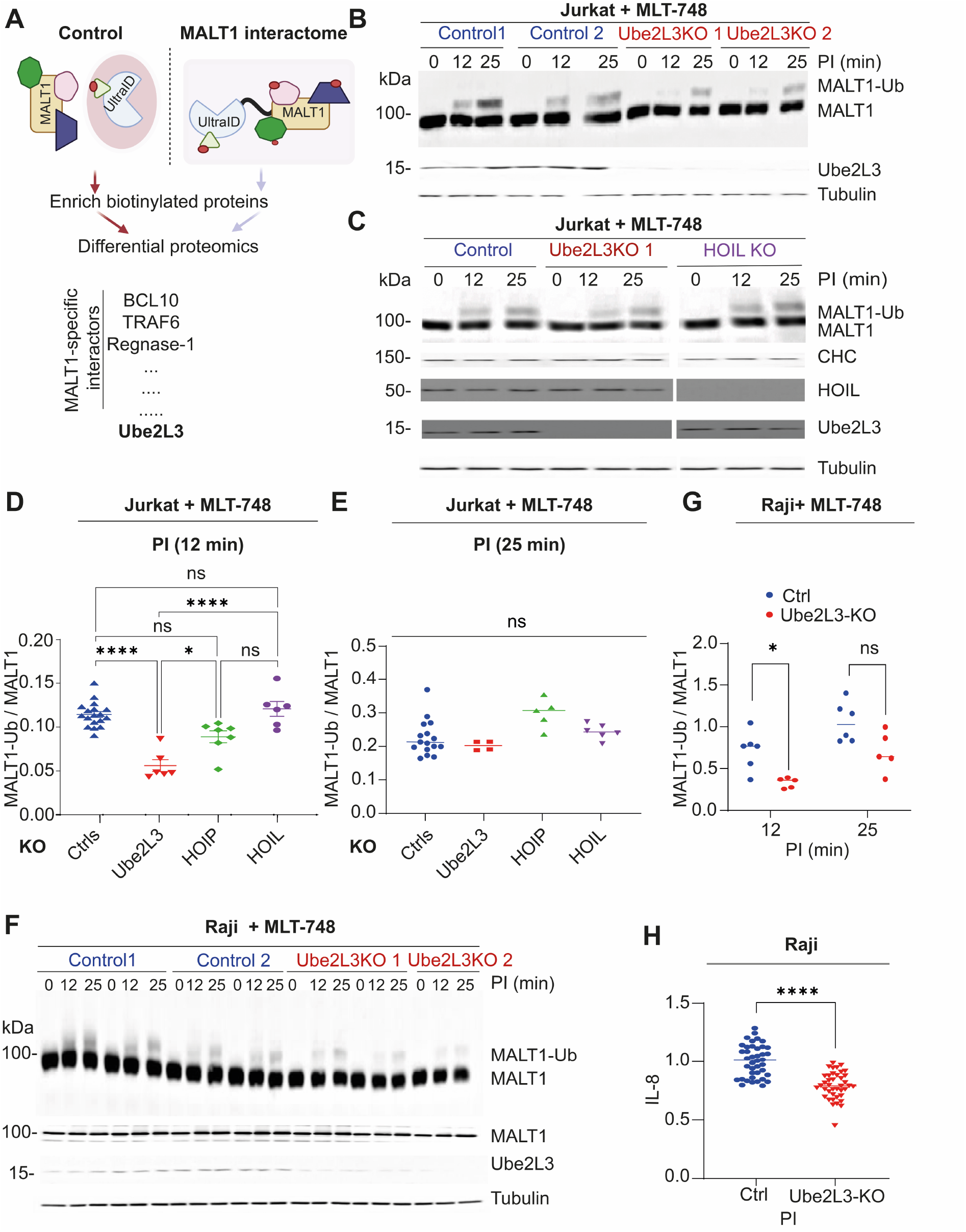
The E2 conjugating enzyme Ube2L3 contributes to MALT1 K644 monoubiquitination in Jurkat and Raji. (**A**) UltraID-MALT1 (**Interactome**) based strategy for identifying new MALT1 interactors and selected hits using ultraID-P2A-MALT1 (**Control**) as a comparison. (**B**, **C**) Jurkat cells transduced with viruses delivering Cas9 and the stated gRNA were treated with MLT-748, PI-stimulated (20 ng/ml PMA, and 1.00 μM ionomycin) for the stated times, and analyzed by western blot as indicated. (**D**, **E**) Quantification of data in **B** (n=18,6,7,6) and **C** (n=16,4,5,6) for the 12 min and 25 min time points of PI-stimulation (3 ng/ml PMA, and 0.15 μM ionomycin), respectively. (**F**) Raji cells transduced with control or Ube2L3-specific CRISPR constructs were treated with MLT-748, PI-stimulated (20 ng/ml PMA, and 1.00 μM ionomycin) for the stated times, and analyzed by western blot as indicated. (**G**) Quantification of data in **F** (n=6,5). (**H**) The stated Raji cells were PI-stimulated (3 ng/ml PMA, and 0.15 μM ionomycin) for 14 h then analyzed for IL-8 secretion by ELISA (n=40). Tubulin was used as a loading control. Data were analyzed by one-or two-way ANOVA, followed by Dunnett’s (one control group) or Tukey test (comparison across whole data set), or in cases where there were only two groups, a two-tailed, unpaired *t*-test; *0.05>p≥0.01; **0.01>p≥0.001; ***0.001>p≥0.0001; ****p<0.0001.

The identification of Ube2L3 was particularly interesting, since we showed above that Ube2L3 can support MALT1 ubiquitination *in vitro*. Relatively few E2-conjugating enzymes have significant transthioesterase activity; most preferentially perform the classical function, conjugation of ubiquitin directly onto lysine (transamidase activity). Ube2L3 is perhaps the best characterized transthioesterase E2, and has the highest known propensity of human E2s to transfer ubiquitin to active sulfur nucleophiles (*40*). Thus, as our data already implicated a cysteine within MALT1, C687, as an acceptor of activated ubiquitin, Ube2L3 seemed an ideal candidate for this process. We thus compared ubiquitination of MALT1 in control and Ube2L3-deficient cells (in two separate polyclonal lines derived from two different Ube2L3-targeting gRNAs). In both lines, there was a reduction of MALT1-monoubiquitination in early timepoints post PI stimulation (**Figure 5B-D**). At later timepoints, suppression of MALT1-monoubiquitination was no longer apparent (**Figure 5E**). As Ube2L3 is believed to be one of the E2s that service the LUBAC, we also investigated how LUBAC silencing affected MALT1-monoubiquitination. HOIL-and HOIP-silencing had no or minimal effect on MALT1-monoubiquitination, indicating that LUBAC activity was not important for MALT1 monoubiquitination (**Fig.5D, Figure S4A**), consistent with data with HOIPIN-8 above (**Figure S3A**). Similar suppression of MALT1-monoubiquitination in the early timepoints post stimulation was found in Raji B cells (**Figure 5F,G**). In Raji cells, Ube2L3 silencing also caused a decrease in CYLD cleavage (**Figure S4B**) and in PI-induced cytokine production (**Figure 5H**). Overall, these findings provide a consistent picture that Ube2L3 is a novel MALT1 interactor and that MALT1 can thus act as a catalytically active E3 ligase for its own monoubiquitination. Given that many E2 enzymes, including Ube2L3, and closely related E2s, have been proposed to service HECT-like E3 ligases, it is likely that several E2s can service MALT1, which would explain why the phenotypes observed were partial (*41*).

### Inactive Ube2L3 inhibits MALT1 monoubiquitination and activity dependent on C687

With these data in hand, we investigated the mechanistic links between Ube2L3 and MALT1 protease activity in more detail. In the absence of known chemical inhibitors of Ube2L3, we mimicked chemically induced loss of Ube2L3 activity by expressing inactive Ube2L3. We examined how this situation affected MALT1 monoubiquitination in transfected HEK293T cells. Co-transfection of inactive Ube2L3, Ube2L3(C86S), reduced MALT1 monoubiquitination by 50 % (**Figure 6A,B**). Consistent with these data, co-transfection of Ube2L3(C86S) or another inactive mutant Ube2L3(H119Q)(*42*), also reduced MALT1-wt protease activity (**Figure 6C**). We then investigated how Ube2L3(C86S) affected MALT1-dependent substrate cleavage in Jurkat T cells. We found that MALT1-monoubiquitination and MALT1 protease activity on substrates with relatively poor cleavage sites (CYLD and HOIL) was reduced in these Jurkat lines(*17*) (**Figure 6D,E, Figure S5A,B**), consistent with our model. Cleavage of substrates with an optimal MALT1 protease cleavage site (Regnase-1 and Roquin-1) were unaffected. Altogether, these results indicate that Ube2L3 inhibition reduces MALT1 activity and thereby modulates MALT1 protease spectrum.

**Figure 6.**
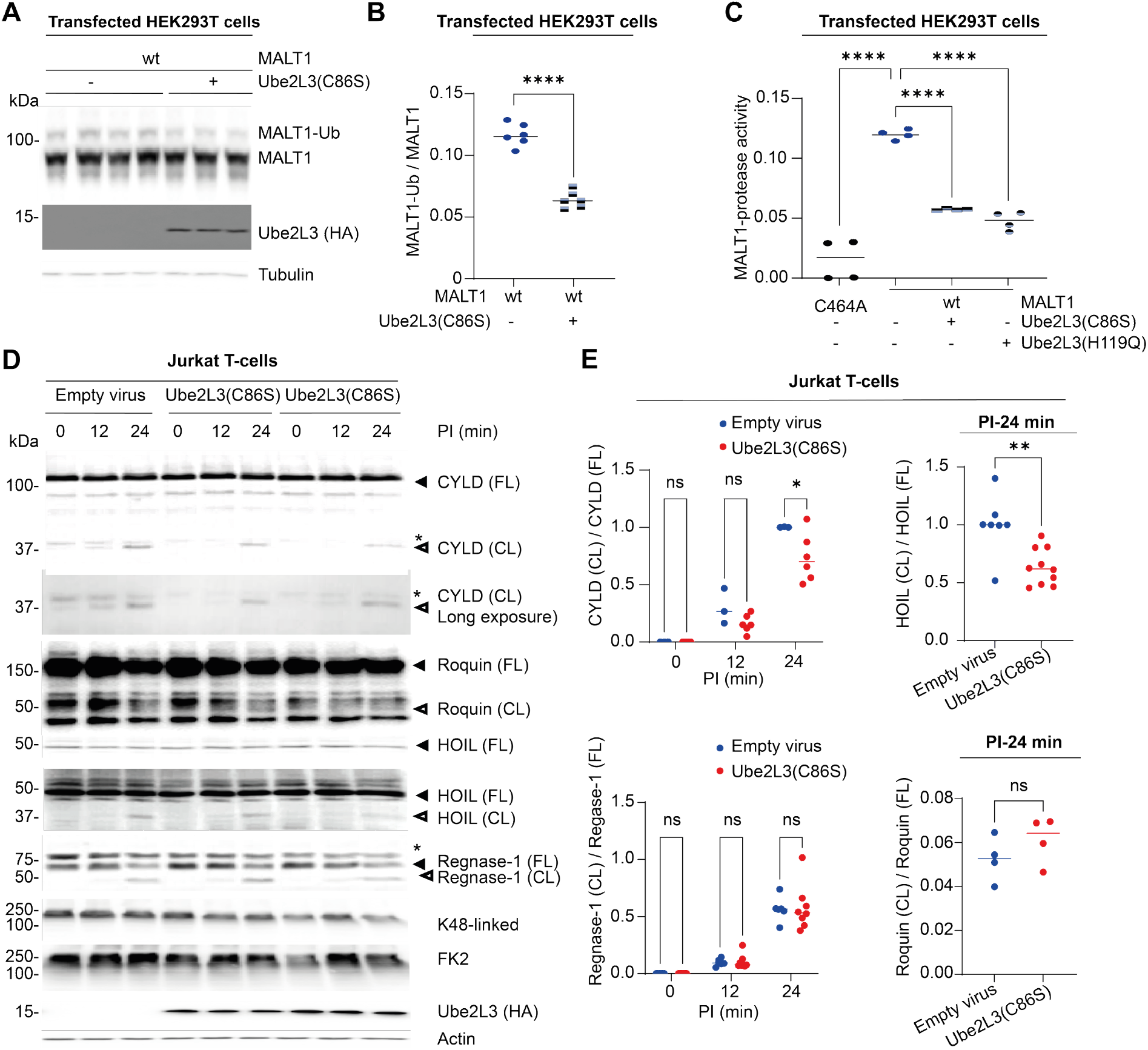
Inactive Ube2L3 suppresses MALT1 monoubiquitination and protease activity. (**A**) HEK293T cells were transfected as described (wt-four replicates; wt + Ube2L3-three replicates) and then analyzed by western blot. (**B**) Quantitation of data in **A** (n=6). (**C**) HEK293T cells were transfected as described and then MALT1 protease activity was measured by flow cytometry using the FRET activity reporter (n=4). (**D**) Jurkat cells were infected with viruses delivering HA-Ube2L3(C86S) (two lines) or an empty virus (one line) and then PI-stimulated (20 ng/ml PMA, and 1 μM ionomycin) as described and analyzed by western blot. (**E**) Quantitation of data in **D** (CYLD: n=3,6; HOIL: n=7,10; Regnase-1 n=4,8; Roquin-1 n=4,4). Tubulin and Actin were used as loading controls. CL – cleaved; FL – full length. Closed arrows indicate full length protein; open arrows indicate cleaved proteins. □ denotes non-specific band. Data were analyzed by one or two-way ANOVA, followed by Dunnett’s test, or for cases where there were only two variables, two-tailed, unpaired *t*-test; *0.05>p≥0.01; **0.01>p≥0.001; ***0.001>p≥0.0001; ****p<0.0001.

Finally, our data indicated that Ube2L3 might be able to bind MALT1, most likely adjacent to the ubiquitin-binding site, UBS1. An analysis of the best docking pose predicted by AlphaFold3 put Ube2L3 close to the UBS1 site, which is necessary if MALT1 binds the ubiquitin-loaded Ube2L3 protein. We identified one residue, E624, that is situated on a loop adjacent to UBS1, but appeared particularly important for the Ube2L3 MALT1 interaction through proximity to a positively charged residue, Ube2L3(K96) (**Appendix Figure S3B**). E624 was mutated to Arginine to create a charge repulsion between Ube2L3 and MALT1. Consistent with this model, in transfected HEK293T cells, MALT1(E624R) was much less ubiquitinated than MALT1-wt in 293T cells (**Figure 7A,B**). The ubiquitination levels of this mutant were not further affected by Ube2L3(C86S) expression, in contrast to MALT1-wt (**Figure 7A,B**). MALT1(E624R) also showed reduced MALT1 protease activity that was overall similar to MALT1-wt cells transfected with inactive Ube2L3(C86S) and to cells expressing the thioester-deficient MALT1(C687A) mutant (**Figure 7C**). Moreover, MALT1(C687A) and the Ube2L3 binding-deficient MALT1(E624R) were less sensitive to protease inhibition by Ube2L3(C86S) expression (**Figure 7C**). MALT1(C687A) was particularly resistant to protease inhibition by Ube2L3(C86S) expression, as was MALT1(C687A)-ubiquitination to Ube2L3 expression (**Figure S5C,D**). Altogether, these results are consistent with a model where Ube2L3-mediated stimulation of MALT1 protease activity is dependent on the active site C687 of the MALT1 E3 ligase activity and on a putative Ube2L3-binding site on the surface of the MALT1 Ig3 domain surrounding E624.

**Figure 7.**
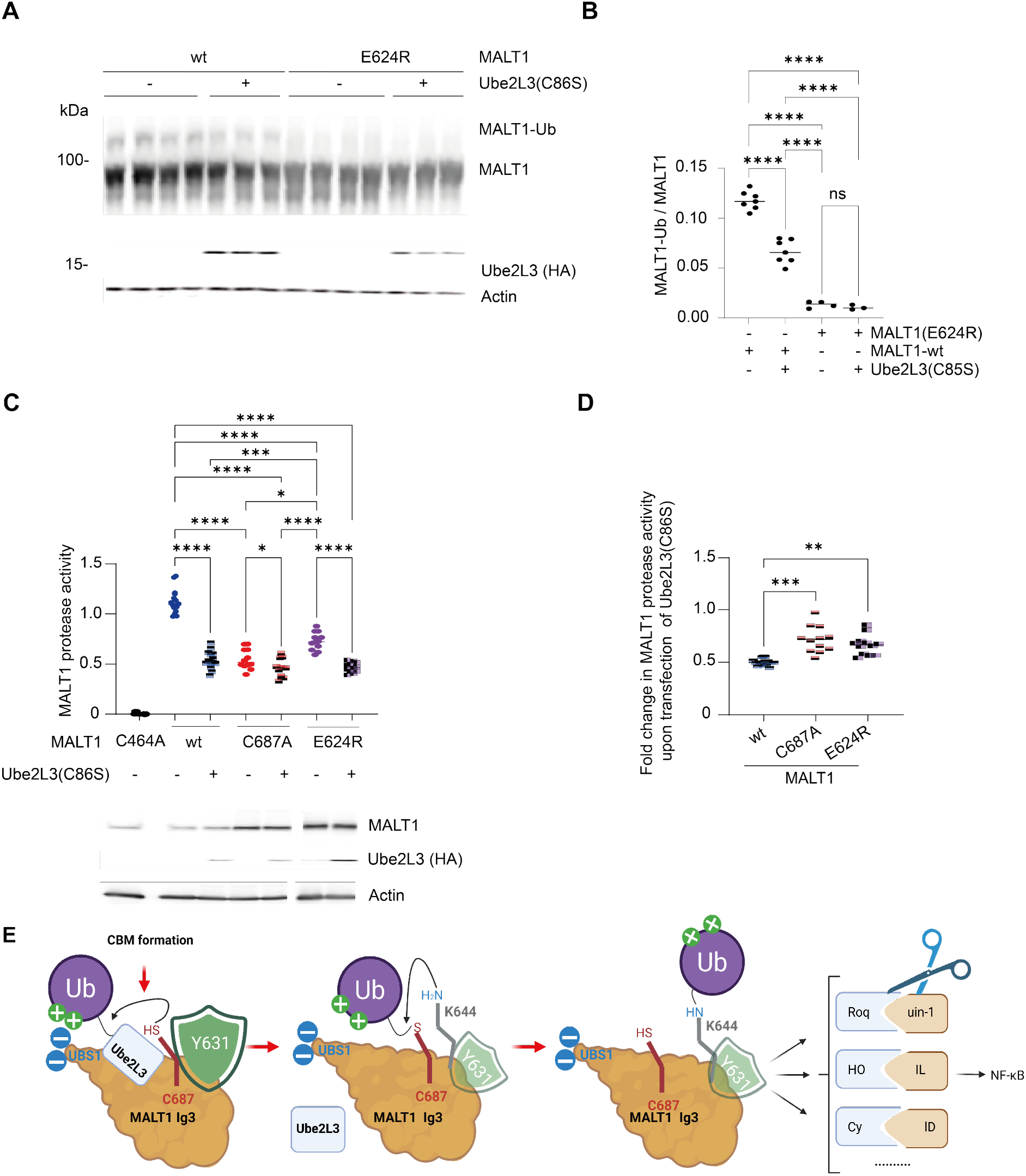
The E2 enzyme Ube2L3 contributes to MALT1 K644 monoubiquitination in Jurkat T-cells. (**A**) HEK293T cells were transfected as described (0.2 equivalents of BCL10), treated with MLT-748, and then MALT1 ubiquitination was assessed by western blot; right hand side quantitation. (**B**) HEK293T cells were transfected as described (0.2 equivalents of BCL10) and then MALT1 ubiquitination was assessed by western blot; below quantitation (n=7,7,4,3). (**C**) HEK293T cells were transfected as described and then MALT1 protease activity was measured using by flow cytometry using the FRET activity reporter (n=12). (**D**) Fold change in MALT1 protease activity upon co-transfection with Ube2L3 (n=12). (**E**) Model for Ube2L3-mediated monoubiquitination of MALT1(K644), mediated by C687 trans-thioesterification and gated by Y631. This process stimulates MALT1 protease activity and promotes NF-κB signaling. Actin was used as loading control. CL – cleaved; FL – full length. Closed arrows indicate full length protein; open arrows indicate cleaved proteins. Data were analyzed by one-way ANOVA, followed by Dunnett’s (one control group) or Tukey test (comparison across whole data set); *0.05>p≥0.01; **0.01>p≥0.001; ***0.001>p≥0.0001; ****p<0.0001.

## Discussion

Here, we provide several lines of evidence supporting a model in which MALT1 acts as an auto-E3 ligase, promoting its own monoubiquitination via the recruitment of a Ub-loaded E2. We identify one E2 servicing MALT1, Ube2L3 (**Figure 7E**). Our mutational analysis identified a cysteine residue (C687) on the surface of the MALT1 Ig3 domain, between the ubiquitin binding site UBS1 and the Ub acceptor site K644, perfectly placed to act as a ubiquitin thioester shuttle from UBS1 to K644. Additionally, we show that a polar aromatic residue, Y631 in human MALT1 (or a histidine residue found in the corresponding position in some other species), acts as a gatekeeper, regulating the initial transfer of Ub from the E2 onto C687. Finally, we show that the proposed mechanism is relevant for the activation of MALT1 protease by monoubiquitination and the responsive capacity of lymphocyte cell lines.

Amongst the different ubiquitinated protein pools, monoubiquitination may be the most prevalent ubiquitin modification in cells (*43*). Protein monoubiquitination is a crucial signaling regulatory event, both as a seed for polyubiquitination and as a signaling function. Protein monoubiquitination can, for example, affect the target protein’s conformation, catalytic activity, subcellular localization, and physical interaction with cellular targets (*44, 45*). Thereby, protein monoubiquitination is linked to a set of conserved signaling pathways involved in fundamental regulatory processes, including transcriptional regulation (*46*). In many instances, specific E3 ligases have been assigned to these processes (*47*). However, in a few instances, protein monoubiquitination occurs in an E3-independent manner (*48*). E3-independent monoubiquitination requires the presence of a ubiquitin binding domain (UBD) in the substrate protein, which typically allows recruitment of a Ub-loaded E2 enzyme for subsequent, direct transfer to the acceptor residue, typically lysine, within the UBD. This type of monoubiquitination often shifts the substrate protein into an inactive conformation, through an intramolecular interaction of the UBD with the newly attached monoubiquitin moiety (*48*). Here we delineated a different autocatalytic mechanism of monoubiquitination that activates the protease MALT1, likely through intermolecular MALT1-ubiquitin interactions (*20*). MALT1 monoubiquitination also requires a UBD but this interaction leads to ubiquitin transfer from an E2 to a conserved cysteine sensor, C687, followed by intramolecular transfer to a specific lysine residue, K644 (**Figure 7E**). It is likely that the large distance between UBS1 and K644 that makes direct transfer between UBS1 and K644 kinetically slow, or not possible, also explains why MALT1 monoubiquitination does not follow the autoinhibition paradigm and why such a transfer mechanism is needed.

Generally, the mechanism proposed here is ascribed to ubiquitin ligases or E2s; it is to our knowledge not known outside of these enzymes. Unsurprisingly, the mechanism of Ubiquitin transfer is reminiscent of HECT E3 ligases and RBRs like the LUBAC, that undergo transesterification with a Ub-E2 conjugate to a catalytically active cysteine. The main difference to the model proposed here for MALT1 is that HECT E3/LUBAC ligases typically deliver ubiquitin intermolecularly to specific targets. However, self-ubiquitination by HECT/LUBAC ligases is also possible (*49*). The ubiquitin transfer mechanism we propose is similar to the E3 ligase MYCBP2 that undergoes internal trans-thioesterification prior to performing ubiquitination (*50*), as well as UBE2O that has a hybrid E2/E3 function involving trans-thioesterification (*51*). Moreover, the E2 Ubc7 can undergo catalytic site cysteine polyubiquitination (which can be transferred to neighboring lysines), regulating its stability (*52*). Cysteine polyubiquitination is also linked to immune responses (*53*). More generally S-to-N acyl transfers are established means of conjugation and primary structure modulation, often leveraged in chemical biology (*54*), but also present in biological systems, such as inteins (*55*). Unsurprisingly, several authors have hinted that ubiquitination processes could well be more complex and diverse than the E1, E2, E3 cascade outlined in traditional biology textbooks (*56*). Many of these unusual examples are found in pathogens (*57*); we provide a notable mammalian example here.

What makes the MALT1 E3 active site C687 special -is it simply location (i.e. proximity to a UBS), or are other properties such as reactivity important? The former is almost certainly a crucial element: C687 is among the most surface-accessible cysteines within MALT1 and well placed to intercept activated ubiquitin bound to UBS1 (*58*) (**Figure 1A**). As for the latter, C687 rarely appears in cysteine activity profiling datasets (*59, 60*) that seek to identify inherently reactive cysteines. However, these methods can only cover a limited range of the >200’000 cysteines in the proteome (*61*). The pKa of C687 was also not flagged as particularly low by PypKa (*62*), but pKa modelling is complex and not particularly well correlated with reactivity (*63*). One interesting observation, that had experimental backing through mutagenesis, was the role of a sulfur-arene interaction in gating transfer from C687 to K644. Cysteine-arene interactions, which are not uncommon in crystal structures (*64*), have previously been proposed to promote cysteine reactivity. However, Y631A was more prone to ubiquitin transfer, suggesting it instead sterically blocked/gated transfer across surfaces. Indeed, aromatic residues are the most common gating residues in enyzmes (*65*) and cysteine-arene interactions are also known to weakly stabilize protein structure (*66*). This suggests the possibility of regulating MALT1 activity via an alteration in the orientation of the gatekeeper residue Y631, which might be inducible e.g. by MALT1 conformational changes occurring in the context of CBM complex formation or upon ubiquitin binding. Indeed, reexamination of 2D HSQC spectra of MALT1 with and without ubiquitin bound to the C-terminus shows that the region around Y631, including D629 that may act as part of the “gate” for ubiquitin transfer, is significantly structurally perturbed upon ubiquitin binding (*20*). Perhaps more simply, MALT1(Y631A) expression provides the means to activate MALT1 protease without changing basal conformational properties through modulating monoubiquitination. This is distinct from previous mutants reported, such as MALT1(Y657A) that alter conformational dynamics of MALT1 protease in the ground state and are not linked to ubiquitination. We further note that when considered against 300 proteins spanning mammalian, reptile, bird and nematode MALT1, Y631, K644, C687, and UBS1 are similarly highly conserved, testifying to the importance of the relay circuit we outline for mammalian MALT1 protease activity.

Of course, another aspect that facilitates thioester exchange followed by transamidation is binding to an E2, most of which prefer to perform transamidation. Interestingly, we identified Ube2L3 as a new MALT1 interactor, an E2 that is arguably the best transthioesterase in the proteome. Our mutagenesis implies that Ube2L3 interacts with MALT1 close to UBS1, as would be expected based on the need for an E2-Ub complex to ubiquitinate MALT1. It thus seems likely that the positioning of UBS1, next to an E2 binding site, and close to an acceptor residue (C687) is crucially important for the sort of autoubiquitination we report here to occur.

Although the data show that C687-mediated monoubiquitination is important in initiating MALT1 ubiquitination on K644, it is clearly not the only pathway by which MALT1 ubiquitination can occur.

Given the known promiscuity and overlapping functions of E2s (these are ∼15-fold less prevalent than E3s), and our own data showing that several E2s can ubiquitinate MALT1 *in vitro*, this is not entirely unsurprising. Indeed, at least 12 % of E2s can perform transthioesterification to the LUBAC (*67*). Such E2s can likely also assume the role of Ube2L3 in MALT1 monoubiquitination, as demonstrated by the fact that MALT1-monoubiquitination still occurred in a reduced manner when Ube2L3 was silenced. Docking of transamidating E2s could also allow direct transfer to K644, which is slower, but would build up overtime, consistent with our observations. Moreover, CBM formation necessitates MALT1 fiber formation that could allow intermolecular transfer of ubiquitin. This process would likely still depend on UBS1, but less on C687. Consistent with such a model, our published phenotypes from UBS1 mutants are stronger than what we observe here for MALT1(C687A). Transfer of Ubiquitin from MALT1-UBS1 to MALT1-K644 *in trans* across a fiber/aggregate, is analogous to a classical E3 ligase function.

MALT1’s protease function is essential for appropriately controlled immune responses, but also for the growth of a variety of cancer cells that are characterized by constitutive MALT1 activity (*1*). MALT1 inhibitors are presently undergoing evaluation in clinical trials for human B-cell lymphomas (*68*), but may not be ideally suited for immunomodulation because MALT1 inhibition also affects the development of regulatory T cells (*25, 26*). Our data open new avenues for therapeutic targeting of MALT1-dependent pathologies, by identifying an additional, E3-like autoubiquitination activity of MALT1 that might be amenable to targeting by small molecule inhibitors. A further, alternative option for a more subtle therapeutic regulation of MALT1 activity could be to target the E2 enzyme(s) servicing MALT1. This avenue is indeed often considered more pharmaceutically feasible than targeting E3-ligases that tend to have no chemical activity (*69*). Inhibition of the E2 enzyme Ubc13-Uev1A, which is crucial for canonical NF-κB activation through K63-linked polyubiquitination of NEMO, for example, inhibits the proliferation and survival of DLBCL cell lines and primary DLBCL cells (*70*). Similarly, redundance in E2s servicing MALT1 could be an advantage, because it may allow selective dampening of MALT1 ubiquitination in specific cell types that may depend on different E2s for activating MALT1. Otherwise, polypharmacologic inhibitors may be able to target a broader subset of cysteine-preferring E2s (*71*). Nonetheless, expression of inactive Ube2L3 was able to suppress MALT1 activity in Jurkat T cells, indicating that development of Ube2L3 inhibitors may be able to exert a dampening effect on MALT1 protease and hence adaptive immune cell activation.

Interestingly, a role for MALT1 as an E3 ligase had been proposed earlier, based on the observation that a cIAP2-MALT1 fusion protein, derived from a chromosomal translocation present in MALT lymphoma, promotes NEMO ubiquitination (*72*). This ubiquitination activity was subsequently attributed to MALT1’s capacity to recruit the E3 ligase TRAF6 (*2*) (an interaction we confirmed by ultraID-MALT1). In our work we did not observe altered, NEMO/IKK-dependent IκB phosphorylation in cells expressing the HECT-E3-inactive MALT1(C687A). It therefore seems unlikely that MALT1 has a HECT-E3 ligase-like function that promotes polyubiquitination of NEMO. Based on our data, MALT1 would likely *not* catalyze polyubiquitin chain formation itself. Instead it would rather catalyze monoubiquitination, either to seed polyubiquitin chains, or to form states analogous to MALT1-Ubiquitin. It is nonetheless interesting to consider whether MALT1 could have two different ubiquitin transfer activities, essentially a scaffold-mediated transfer (traditional or RING-like) and a transfer of chemically activated ubiquitin (referred to here as HECT-like). The different substrates and biological contributions that would result from these two functions are particularly interesting questions that are beyond the scope of this initial paper. However, the literature provides several examples of E3s that conjugate ubiquitin on both their target proteins and themselves (*73*).

In conclusion, our findings identify MALT1 as an E3-ligase catalyzing its own activation by monoubiquitination in a process involving Ube2L3, opening diverse options for therapeutically targeting MALT1 activity in the context of immunomodulation and cancer therapy.

## Materials and Methods

| ***Antibodies*** | **Source** | **Identifier** |
|---|---|---|
| *Roquin-1* | Abcam | Ab70195 |
| *MALT1* | Homemade |  |
| *Regnase-1* | R&D | MAB7875 |
| *BCL10 (FL)* | SantaCruz | SC-5273 |
| *Biotin-HRP conjugate* | Cell Signaling | D5A7 |
| *HA* | BioLegend | 16B12 |
| *HOIL-1* | SantaCruz | SC-365523 |
| *CYLD* | Cell Signaling | D1A10 |
| *Myc-tag* | Cell Signaling | 71D10 |
| *HOIP* | Cell Signaling | E6M5B |
| *IkBα* | Cell Signaling | L35A5 |
| *GFP* | ENZO | CAC-FRL-GFP-RB-AF2020-20-EX |
| *pERK* | Sigma | M8159 |
| *pIκBα Ser32/36* | Cell Signaling | 5A5 |
| *Tubulin* | Sigma | T5168 |
| *CD3* | Invitrogen | clone OKT3 |
| *CD28* | Invitrogen | CD28.2 |
| *Cross-linking antibody* | Jackson Immunoresearch | 115-006-062 |
| *CHC* | BD | 610500 |
| *Actin* | SantaCruz | SC-47778 |
| *Linear ubiquitin* | Sigma | ZRB2114 |
| *FK2 (mono-and polyubiquitinated proteins)* | ENZO | LSI-AB-0120 |
| *K48-linked ubiquitin* | Abcam | EP8589 |
| *Ube2L3* | Cell Signaling | 3848S |
| **Materials & chemicals** | ***Source*** | ***Identifier*** |
| *RPMI medium with glutamax* | Gibco | 61-870-127 |
| *Biotin-free RPMI medium plus glutamine* | MyBioSource | MBS653376 |
| *DMEM* | GIBCO | D5671 |
| *phorbol 12-myristate 13-acetate* | Sigma | 79346 |
| *Ionomycin* | Sigma | I0634 |
| *DMSO* | SIGMA | D2438 |
| *Streptavidin magnetic beads* | Pierce | 88817 |
| *IL-2 ELISA* | BD OptEIA | 555190 |
| *IL-8 ELISA* | Immunotools | 31670089 |
| *Puromycin* | Sigma | P7255 |
| *Blasticidin S hydrochloride* | Sigma | 15205 |
| *G418* | Gibco | 10131027 |
| *SuperSignal West Dura* | ThermoFisher | 34076 |
| *BD LSR-II* | BD biosciences |  |
| *Spectramax ID5* | Molecular devices |  |
| *MLT-748* | Medchemexpress | HY-115466 |
| *Nickel NTA agarose* | Qiagen | 30210 |
| *HOIPIN-8* | MedChemExpress | HY-122882 |
| *Pevonedistat* | MedChemExpress | HY-10484 |
| MRT67307 | MedChemExpress | HY-13018A |
| TAK-243 | MedChemExpress | HY-100487 |
| 3,3',5,5'-tetramethylbenzidine (TMB) | Thermoscientific | 34028 |
| *IPTG* | GoldBio | I2481C5 |
| **Base Plasmids** | ***Source*** | ***Identifier*** |
| pWPI | Addgene | 12254; Trono Laboratory |
| *pWPI-MCS-IRES-GFP-Blasticidin(r)* | This paper |  |
| *pWPI-ultraID-MCS-IRES-GFP-Blasticidin(r)* | This paper |  |
| *pLentiCRISPRV2 NEO* | Addgene | 127644; Manel Laboratory |
| *pLentiCRISPRV2 PURO* | Addgene | 52961; Zhang Laboratory |
| *pLentiCRISPRV2 BLAST* | Addgene | 52962; Zhang Laboratory |
| *PSPAX2* | Addgene | 12260; Trono Laboratory |
| *pCMV VSV-G* | Addgene | 8454; Weinberg Laboratory |
| *pET21d Ube1* | Addgene | 34965; Wolberger Laboratory |
| *pET28aUbe2V2* | This paper |  |
| *pET21(+)Ubc13* | Addgene | 212715; Ulrich Laboratory |
| *pET15bUbcH5C* | Addgene | 11473; Weissman Laboratory |
| **Software** | ***Source*** | ***Use*** |
| *Graphpad Prism* | Graphpad | Data preparation, curve fitting and analysis |
| *IMAGE J* | National Institutes of Health | Western blot quantitation |
| *ChimeraX* | UCSF | Calculation of RMSD, superimposing structures, etc. |
| *Weblogo* | UC Berkeley | Creation of logos |
| *Chemdraw* | Revvity Signals | Chemical structures |
| *Alphafold3* | Google DeepMind and EMBL-EBI | Modeling and docking |
| *Biorender* | Biorender (licenced to MJCL) | Diagrams |
| *ChopChop* | https://chopchop.cbu.uib.no/ | gRNA design |

### Cell culture and stimulation

All cells were cultured at 37 °C in a humidified atmosphere with 5 % CO_2_. HEK293T cells were cultured in Dulbecco’s Modified Eagle Medium (DMEM) in 10 % fetal calf serum (FCS, Biowest). Jurkat T cells, and Raji cells were cultured in Roswell Park Memorial Institute (RPMI)-1640 medium supplemented with 10 % FCS. For stimulation, cells were maintained at 2 million/ml in 6-well plates and equilibrated in an incubator for at least 2 h. Typically, 6 million cells in total were used per condition. A mixture of phorbol 12-myristate 13-acetate (PMA; 20 ng/mL; Sigma) and ionomycin (1 μM; Sigma) (referred to as PI concentration = 20, e.g. **Figure 4h**), delivered as a 1000x concentration in DMSO was used for stimulation (PI). For stimulation of Raji cells to measure cytokine (IL-8) production, 3 ng/ml PMA, and 0.15 μM ionomycin was used, otherwise standard concentrations were used. Time courses were carried out as reverse kinetic, with inhibitor pretreatment [HOIPIN-8 (15 µM), TAK-243 (1 µM), Pevondistat (1 µM) (MedChemExpress) or DMSO occurring on all samples prior to stimulation]. CM-TMR Cells were lysed in *lysis buffer*: 50 mM HEPES, 150 mM NaCl, 1 % Triton X-100, pH 7.6 with ROCHE complete inhibitor cocktail (2X), PR-619 (50 µM), Na_3_VO_4_ (2 mM), NaF (2.5 mM), and Na_4_P_2_O_5_ (500 µM). Inhibitors were added fresh each time. To observe MALT1-monoubiquitination, the same lysis buffer supplemented with 0.5 % sodium deoxycholate could also be used, although this did not yield consistently better results. MLT-748 MedChemExpress was used at 10 µM. TMB (3,3’,5,5’-tetramethylbenzidine) was from Thermoscientific and read out by absorption at 450 nm after dilution 50 % in 2 M HCl.

### Western blot and antibodies

Western blot was performed in Towbin buffer, 20 % ethanol, on ice for 84 V for 1.5 h, followed by 124 V for 30 min. To visualize MALT1 ubiquitination, a 6.25 % Anderson gel (made with fresh mix) was best used. Membrane was stained in Ponceau, washed, blocked in 10 % milk in PBS-Tween, exposed to primary antibody at 4 °C overnight in 5 % milk in PBS-Tween, then washed 3 times, and exposed to secondary antibody at 1:10’000 dilution in 5 % milk in PBS-Tween (1 h). This was washed 3 times over 40 min, then developed using SuperSignal reagents (Pierce) and visualized using a Vilbur Fusion imager. Antibodies: RelB (Cell Signaling, 4954); Roquin-1 (Abcam); Regnase-1 (R&D, MAB7875); BCL10 (SantaCruz, SC-5273); biotin-HRP conjugate (Cell Signaling, D5A7); HOIL-1 (SantaCruz, SC-365523); CYLD (Cell Signaling, D1A10); HOIP (Cell Signaling, E6M5B); IκBα (Cell Signaling, L35A5); pERK (Sigma, M8159); pIκBα Ser32/36 (Cell signaling, 5A5); tubulin (Sigma, T5168); CD3 (Invitrogen, clone OKT3); CD28 (Invitrogen, CD28.2); cross-linking antibody (Jackson Immunoresearch, 115-006-062); CHC (BD, 610500); β-actin (Santa Cruz, SC-47778). Assignment of cleavage and non-specific bands was made based on absence of corresponding bands in KO cell lines or mutant expressing cells or in cells treated with MLT-748. Ubiquitin antibodies were validated using positive controls, such as bortezomib, HOIPIN-8 (e.g. **S3A**) and deubiquitinating enzymes.

### Western blot and imaging quantifications

These were performed using ImageJ using the lane quantification tool. To calculate ratios of ubiquitinated protein : full length protein, or of cleaved : full length proteins, the same blot at the same exposure was used unless one of the components was overexposed, then different exposures were used.

### Transfection and transduction of cells

Transient transfection of HEK293T cells and lentiviral transduction of cells were enacted as described in prior publications (*19, 20*). For expression, HEK293T cells were diluted 1:10 DMEM with 10 % FCS and 1 % penicillin streptomycin (Gibco). After 18-25 h, cells were transfected, media were replaced after 6 h, and then 18-25 h later, cells were harvested. To silence protein expression, cells were infected with lentiviruses delivering Cas9 and a single guide RNA (sgRNA) and selected using puromycin (1 µg/ml), blasticidin (15 µg/ml) or G418 (1 mg/ml), using vectors derived from pLENTICRISPRV2. To rescue expression, cells were subsequently transduced to constitutively express the protein of interest together with GFP-BSR (pWPI-MCS-IRES-GFP-BLASTICIDIN(r)). Only polyclonal lines were used. Silenced lines were used independently, but in groups with replicates shown side by side in blots or as points in ELISAs.

### Identification of MALT1 binding partners using ultraID-MALT1 and ultraID-P2A-MALT1

MALT1-KO Jurkat lines were reconstituted separately with the above constructs that give constitutive expression and co-express GFP and blasticidin resistance, driven by an IRES (pWPI-ultraID-GENE-IRES-GFP-Blasticidin(r)). These were grown in biotin-free RPMI medium plus glutamine (MyBioSource, MBS653376). Biotin (200 µM) was added for 18 h prior to lysis. Approximately 100 million cells were used per replicate. Cells were lysed in cell lysis buffer to a concentration of ∼10 mg/ml, and lysates were diluted to 1 mg/ml in the same buffer with a final concentration of 1% LDS. > 4 mg total lysate was incubated with streptavidin magnetic beads (88817, Pierce) with end over end rotation for 2 h at RT. Beads were washed once with PBS (500 mM NaCl), then 3 times with PBS and then analyzed by mass spectrometry using label free analysis. Each set was performed in triplicate, and data were compared between ultraID-MALT1 and ultraID-P2A-MALT1 expressing cells. Proteins significantly enriched in cells expressing ultraID-MALT1 versus ultraID-P2A-MALT1 were considered positive hits.

### Protein expression

Plasmid (pET28a backbone) was transformed into BL21(DE3) *E. coli*, then grown in a starter culture to saturation at 37 °C in L-Broth. Culture was then grown to saturation again, and diluted into 2 L L-Broth and grown to an OD of 0.6. Culture was chilled on ice to 17 °C, and then IPTG (final concentration 0.2 mM) was added and grown overnight. *E. coli* were centrifuged and then lysed in (50 mM HEPES, 500 mM NaCl, 10 mM imidazole, 1 mM BME, 1 % NP-40, pH 7.6 with ROCHE complete EDTA free protease cocktail) using a French Press. Lysate was clarified, then bound to Nickel resin (Qiagen) under rotation at 4 °C. Resin was loaded into a minicolumn and then washed 4 times in lysis buffer minus NP-40, the with lysis buffer with increasing concentrations of imidazole. Protein elution was detected by gel analysis. Fractions were dialyzed into 50 mM HEPES, 150 mM NaCl, 2 mM BME, 50 % glycerol by stepwise reduction of NaCl concentration and then concentrated.

### IL-2/IL-8 ELISA

Cells were counted and then diluted to 1 million cells per mL; typical starting concentrations were 2 million cells per ml. For B-cells, cells were then centrifuged, old medium was discarded, and then fresh medium was replaced. 160 µL cells were placed in each well of a round bottomed 96 well plate. These were left for 30 min-1 h at 37 °C. Then PI (or anti-CD3, anti-CD28 with cross linker) were added as a 5x solution. Cells were incubated for 14-18 h and then centrifuged at 1000g. Supernatants were taken and then cytokine secretion was measured using an ELISA kit for IL-2 (Beckton Dickinson, OptEIA) or IL-8 (Immunotools, 31670089). Detection was performed using 3,3’,5,5’-tetramethylbenzidine (TMB) and readouts were taken on a Spectromax ID5 (Molecular Devices).

### Crispr/CAS9 and virus production

Guide RNAs were created using CHOPCHOP. These were used to create individual lines that gave similar data in each case. All constructs were cloned into pLENTICRISPRV2 carrying either puromycin, neomycin, or blasticidin resistance as needed, following the guide on ADDGENE. The following sequences were used to target the stated genes. Most sequences are exon targeting aimed at the center of the gene. Some sequences target intron/exon junctions and were used for rescue experiments. MALT1: GGCATCGCCCTCCTTTGTCA; ATTATACCTCTTCACAGCTG; HOIL: CAGCTTGCCCGAGCACGAGT; AGAGACGCCACTGTCATATC; HOIP: TTGACACCACGCCAGTACCG; TCCACCGTGCTGCGAAAGAC; Ube2L3: TGTCTTACCTTGGTCGGTTT; GGCACATGTCTTACCTTGGT; Cells were selected with 1 µg/ml puromycin (Sigma); 1 mg/ml neomycin (Gibco); and 25 µg/ml blasticidin (Sigma). Expression was driven by infection with variants of pWPI vector custom made in the laboratory, including pWPI-MCS-IRES-GFP-BLASTICIDIN(r) and pWPI-ultraID-MCS-IRES-GFP-BLASTICIDIN(r) (where MCS is a multicloning site, and BALSTICIDIN(r) is a gene encoding blasticidin resistance). These were all created by Hawaii cloning as described (*74*) and sequenced in the relevant regions. 293T cells grown in DMEM (Gibco) with 10 % FCS (Biowest) were split 1:10 from a confluent 10 cm plate to a fresh 10 cm plate. After 18 h these cells were transfected with the relevant vector (pWPI for expression or pLENTICRISPRV2) 10 µg, PSPAX2 (7.5 µg) and pCMV VSV-g (5 µg) using calcium phosphate. Medium was changed to OPTIMEM (Gibo) with 1 % FCS and 1x penicillin streptomycin (Gibco) (6 ml) and left for 2 days. Medium was then filtered through 0.45-micron filter, into 3.5 ml 1X Dulbecco’s PBS (Gibco), 2 ml PEG 8000 (200 g in 200 ml PBS), and 0.33 ml 5 M NaCl. This was precipitated overnight at 4 °C during end-to-end rotation, then centrifuged at 5000 g for 1 h. Supernatant was decanted and residue aspirated using a pipette, then pellet was resuspended in 100 µL PBS. For expression, supernatants from up to three 10 cm plates were pooled, precipitated (using appropriately scaled concentrations of the above) and resuspended in 100 µL PBS to improve infection. This mixture was added to 300 µL cells at 1 million per ml in a 24 well plate and left for 3 days. Medium was diluted to 1 ml, then 2 ml as cells grew. Medium was aspirated and replaced with 500 µL fresh medium. After cells had been washed 3 times, they were considered virus free and selection commenced.

### FRET-based MALT1 protease activity assay

FRET-based MALT1 protease activity assays were performed as described (*20*) using an EYFP-LVSR-ECFP reporter construct that was co-transfected in 293T cells together with indicated MALT1 constructs and CARMA1, as indicated. 24 h after transfection, cells were analysed by flow cytometry using a BD LSR-II instrument (BD Biosciences) containing 405, 488, 561 and 640 nm lasers. ECFP and FRET signals were measured by exciting the transfected cells with a 405 nm laser, using a standard 450/50 filter to collect the ECFP fluorescence and a 585/42 filter for FRET fluorescence, respectively. At least 5000 EYFP-ECFP positive cells were counted for each sample.

### Molecular docking

The entire sequence of MALT1 and the stated E2 (Ube2L3-wt and Y631A, Ube2a (27 % identity with Ube2L3), UBC13 (33 % identity with Ube2L3), and Ube2D2 (38 % identity with Ube2L3)) were input into AlphaFold server and analyzed by AlphaFold3. The calculation was run a total of 10 times using the “clone and reuse” function. In most cases a random seed was used, although in some cases a specified seed was used. The top ranked structure from each batch run was analyzed based on ranking score, and then the top two ranking scores were used. These were above 0.3. Hits were analyzed using ChimeraX. Only Ube2L3 showed docking poses near UBS1 and *syn* to C687.

### Modeling conformational flexibility

Conformational flexibility was modelled in several ways. (1) AlphaFold3: MALT1 structure was searched on AlphaFold server. The top ∼50 structures in the search function were downloaded and input into ChimeraX. The Ig3 domain was selected and was overlapped using the Matchmaker function (chain pairing specific chain in reference structure and best aligning chain in match structure; also restrict to selection tab checked). Rotamers of C687 were analyzed using Dunbrack conformations, and the rotamer with cysteine *syn* to K644 was chosen. Similar was done for K644, with the rotamers closest to C687 chosen. The shortest distances between the K644-amine and C687-sulfur were chosen. (2) similar analysis was performed on the gamut of human MALT1 structures from the PDB. (3) b-factors from (2) were analyzed from these structures using ChimeraX. (4) the conformational poses of the K644-containing loop were modelled in Moma (AFflecto) as described using 3uo8 as a basis.

## Data availability and accession codes

The minimum data sets are already available in the paper. Primary data are available on request. Mass spectrometry data have been deposited to the PRIDE database (PXD069479) and detailed information can be found in Table S1.

## Author contributions

MJCL: performed in cell experiments, bar optimization of MS data, and some work with ultraID, and helped with paper writing; ML: optimized ultraID experiments and aided with data collection; CC: provided technical assistance with all experiments in cell culture; RC: prepared MALT1(Y631A) for NMR; CD: provided technical assistance for cell culture and line maintenance; GH oversaw NMR data collection; MT: project oversight, interpretation of data, initial model generation, paper writing.

## Acknowledgements

We are grateful to Manfredo Quadroni for support by the Protein Analysis Facility at UNIL. This study was supported by the Swiss Cancer League (KFS-5411-08-2021, to M.T. and M.J.C.L.), the Swiss National Science Foundation (310030_214974/1, to M.T.), Emma Muschamp Foundation (to M.T.) and Novartis Foundation for Medical-Biological Research (23B094, to M.J.C.L.). Figures were made using Biorender where applicable (licensed to M.J.C.L.).

## References

1. M. Juilland, M. Thome, Holding All the CARDs: How MALT1 Controls CARMA/CARD-Dependent Signaling. Frontiers in Immunology 9, (2018).

2. T. J. O’Neill, A. Gewies, T. Seeholzer, D. Krappmann, TRAF6 controls T cell homeostasis by maintaining the equilibrium of MALT1 scaffolding and protease functions. Frontiers in Immunology Volume 14 - 2023, (2023).

3. Y. Shimizu, L. Taraborrelli, H. Walczak, Linear ubiquitination in immunity. Immunol Rev 266, 190–207 (2015).

4. C. Pelzer et al., The protease activity of the paracaspase MALT1 is controlled by monoubiquitination. Nat Immunol 14, 337–345 (2013).

5. B. Coornaert et al., T cell antigen receptor stimulation induces MALT1 paracaspase–mediated cleavage of the NF-κB inhibitor A20. Nature Immunology 9, 263–271 (2008).

6. J. Staal et al., T-cell receptor-induced JNK activation requires proteolytic inactivation of CYLD by MALT1. EMBO J 30, 1742–1752 (2011).

7. S. Hailfinger et al., Malt1-dependent RelB cleavage promotes canonical NF-κB activation in lymphocytes and lymphoma cell lines. Proceedings of the National Academy of Sciences 108, 14596–14601 (2011).

8. T. Uehata et al., Malt1-Induced Cleavage of Regnase-1 in CD4+ Helper T Cells Regulates Immune Activation. Cell 153, 1036–1049 (2013).

9. K. M. Jeltsch et al., Cleavage of roquin and regnase-1 by the paracaspase MALT1 releases their cooperatively repressed targets to promote T(H)17 differentiation. Nat Immunol 15, 1079–1089 (2014).

10. M. Juilland, M. Thome, Role of the CARMA1/BCL10/MALT1 complex in lymphoid malignancies. Current Opinion in Hematology 23, 402–409 (2016).

11. M. Jaworski et al., Malt1 protease inactivation efficiently dampens immune responses but causes spontaneous autoimmunity. The EMBO Journal 33, 2765–2781 (2014).

12. A. Gewies et al., Uncoupling Malt1 Threshold Function from Paracaspase Activity Results in Destructive Autoimmune Inflammation. Cell Reports 9, 1292–1305 (2014).

13. J. W. Yu et al., MALT1 Protease Activity Is Required for Innate and Adaptive Immune Responses. PLoS One 10, e0127083 (2015).

14. F. Bornancin et al., Deficiency of MALT1 Paracaspase Activity Results in Unbalanced Regulatory and Effector T and B Cell Responses Leading to Multiorgan Inflammation. The Journal of Immunology 194, 3723–3734 (2015).

15. F. Schlauderer et al., Structural Analysis of Phenothiazine Derivatives as Allosteric Inhibitors of the MALT1 Paracaspase. Angewandte Chemie International Edition 52, 10384–10387 (2013).

16. A. G. Uren et al., Identification of paracaspases and metacaspases: two ancient families of caspase-like proteins, one of which plays a key role in MALT lymphoma. Molecular cell 6, 961–967 (2000).

17. J. Hachmann et al., Mechanism and specificity of the human paracaspase MALT1. Biochem J 443, 287–295 (2012).

18. S. Hailfinger, C. Pelzer, M. Thome, Detection and measurement of paracaspase MALT1 activity. Methods Mol Biol 1133, 177–188 (2014).

19. F. Rebeaud et al., The proteolytic activity of the paracaspase MALT1 is key in T cell activation. Nature Immunology 9, 272–281 (2008).

20. R. Schairer et al., Allosteric activation of MALT1 by its ubiquitin-binding Ig3 domain. Proceedings of the National Academy of Sciences 117, 3093–3102 (2020).

21. C. Pelzer et al., UBE1L2, a Novel E1 Enzyme Specific for Ubiquitin. Journal of Biological Chemistry 282, 23010–23014 (2007).

22. C. Michelle, P. Vourc’h, L. Mignon, C. R. Andres, What was the set of ubiquitin and ubiquitin-like conjugating enzymes in the eukaryote common ancestor? J Mol Evol 68, 616–628 (2009).

23. S. Singh, J. Ng, J. Sivaraman, Exploring the “Other” subfamily of HECT E3-ligases for therapeutic intervention. Pharmacology & Therapeutics 224, 107809 (2021).

24. J. Weber, S. Polo, E. Maspero, HECT E3 Ligases: A Tale With Multiple Facets. Frontiers in Physiology Volume 10 - 2019, (2019).

25. I. K. Kerzeli et al., MALT1 inhibition suppresses antigen-specific T cell responses. Cellular Immunology 397-398, 104814 (2024).

26. K. Martin et al., Pharmacological Inhibition of MALT1 Protease Leads to a Progressive IPEX-Like Pathology. Front Immunol 11, 745 (2020).

27. S. Biswas et al., Pharmacological Inhibition of MALT1 Ameliorates Autoimmune Pathogenesis and Can Be Uncoupled From Effects on Regulatory T-Cells. Frontiers in Immunology 13, (2022).

28. A. Demeyer et al., Long-Term MALT1 Inhibition in Adult Mice Without Severe Systemic Autoimmunity. iScience 23, 101557 (2020).

29. M. Di Pilato et al., Targeting the CBM complex causes T(reg) cells to prime tumours for immune checkpoint therapy. Nature 570, 112–116 (2019).

30. M. Rosenbaum et al., Bcl10-controlled Malt1 paracaspase activity is key for the immune suppressive function of regulatory T cells. Nat Commun 10, 2352 (2019).

31. S. M. Marino, V. N. Gladyshev, Cysteine function governs its conservation and degeneration and restricts its utilization on protein surfaces. J Mol Biol 404, 902–916 (2010).

32. L. Yuan, Z. Lv, M. J. Adams, S. K. Olsen, Crystal structures of an E1–E2–ubiquitin thioester mimetic reveal molecular mechanisms of transthioesterification. Nature Communications 12, 2370 (2021).

33. H. Qian et al., Structure and Function of HECT E3 Ubiquitin Ligases and their Role in Oxidative Stress. J Transl Int Med 8, 71–79 (2020).

34. E. Weerapana et al., Quantitative reactivity profiling predicts functional cysteines in proteomes. Nature 468, 790–795 (2010).

35. M. J. C. Long, Y. Aye, Let’s get biophysical – How to get your favorite protein’s digits. Bioorganic & Medicinal Chemistry 112, 117873 (2024).

36. M. V. Shapovalov, R. L. Dunbrack, Jr., A smoothed backbone-dependent rotamer library for proteins derived from adaptive kernel density estimates and regressions. Structure 19, 844–858 (2011).

37. J. Quancard et al., An allosteric MALT1 inhibitor is a molecular corrector rescuing function in an immunodeficient patient. Nature Chemical Biology 15, 304–313 (2019).

38. S. Jackson, Y. Xiong, CRL4s: the CUL4-RING E3 ubiquitin ligases. Trends in Biochemical Sciences 34, 562-570 (2009).

39. M. J. C. Long, Y. Aye, Climbing into their Skin to Understand Contextual Protein-Protein Associations and Localizations: Functional Investigations in Transgenic Live Model Organisms. Chembiochem, e202400005 (2024).

40. D. M. Wenzel, A. Lissounov, P. S. Brzovic, R. E. Klevit, UBCH7 reactivity profile reveals parkin and HHARI to be RING/HECT hybrids. Nature 474, 105–108 (2011).

41. S. Kumar, W. H. Kao, P. M. Howley, Physical interaction between specific E2 and Hect E3 enzymes determines functional cooperativity. J Biol Chem 272, 13548–13554 (1997).

42. X. Zhang et al., Mechanism and Disease Association With a Ubiquitin Conjugating E2 Enzyme: UBE2L3. Front Immunol 13, 793610 (2022).

43. S. E. Kaiser et al., Protein standard absolute quantification (PSAQ) method for the measurement of cellular ubiquitin pools. Nat Methods 8, 691–696 (2011).

44. W. Magits, A. A. Sablina, The regulation of the protein interaction network by monoubiquitination. Current Opinion in Structural Biology 73, 102333 (2022).

45. T. Nakagawa, K. Nakayama, Protein monoubiquitylation: targets and diverse functions. Genes Cells 20, 543–562 (2015).

46. F. Zhao, C. W. Hicks, C. Wolberger, Mechanism of histone H2B monoubiquitination by Bre1. Nature Structural & Molecular Biology 30, 1623–1627 (2023).

47. S. Chen, C. Pan, J. Huang, T. Liu, ATR limits Rad18-mediated PCNA monoubiquitination to preserve replication fork and telomerase-independent telomere stability. The EMBO Journal 43, 1301–1324-1324 (2024).

48. D. Hoeller et al., E3-independent monoubiquitination of ubiquitin-binding proteins. Mol Cell 26, 891–898 (2007).

49. L. M. Fennell et al., Site-specific ubiquitination of the E3 ligase HOIP regulates apoptosis and immune signaling. The EMBO Journal 39, e103303 (2020).

50. P. D. Mabbitt et al., Structural basis for RING-Cys-Relay E3 ligase activity and its role in axon integrity. Nat Chem Biol 16, 1227–1236 (2020).

51. K. Ullah, E. Zubia, M. Narayan, J. Yang, G. Xu, Diverse roles of the E2/E3 hybrid enzyme UBE2O in the regulation of protein ubiquitination, cellular functions, and disease onset. Febs j 286, 2018–2034 (2019).

52. T. Ravid, M. Hochstrasser, Autoregulation of an E2 enzyme by ubiquitin-chain assembly on its catalytic residue. Nat Cell Biol 9, 422–427 (2007).

53. P. Gao et al., E3 ligase Nedd4l promotes antiviral innate immunity by catalyzing K29-linked cysteine ubiquitination of TRAF3. Nature Communications 12, 1194 (2021).

54. H. M. Burke, L. McSweeney, E. M. Scanlan, Exploring chemoselective S-to-N acyl transfer reactions in synthesis and chemical biology. Nature Communications 8, 15655 (2017).

55. N. H. Shah, T. W. Muir, Inteins: Nature’s Gift to Protein Chemists. Chem Sci 5, 446–461 (2014).

56. L. T. Henneberg, B. A. Schulman, Decoding the messaging of the ubiquitin system using chemical and protein probes. Cell Chemical Biology 28, 889–902 (2021).

57. C. G. Roberts, T. G. Franklin, J. N. Pruneda, Ubiquitin-targeted bacterial effectors: rule breakers of the ubiquitin system. Embo j 42, e114318 (2023).

58. G. S. Biggs et al., Robust proteome profiling of cysteine-reactive fragments using label-free chemoproteomics. Nature Communications 16, 73 (2025).

59. K. Yang et al., Accelerating multiplexed profiling of protein-ligand interactions: High-throughput plate-based reactive cysteine profiling with minimal input. Cell Chemical Biology 31, 565–576.e564 (2024).

60. M. E. H. White, J. Gil, E. W. Tate, Proteome-wide structural analysis identifies warhead-and coverage-specific biases in cysteine-focused chemoproteomics. Cell Chemical Biology 30, 828–838.e824 (2023).

61. X. Liu, M. J. C. Long, Y. Aye, Proteomics and Beyond: Cell Decision-Making Shaped by Reactive Electrophiles. Trends in biolochemical sciences, (2018).

62. P. B. P. S. Reis, D. Vila-Viçosa, W. Rocchia, M. Machuqueiro, PypKa: A Flexible Python Module for Poisson–Boltzmann-Based pKa Calculations. Journal of Chemical Information and Modeling 60, 4442–4448 (2020).

63. S. Parvez, M. J. Long, J. R. Poganik, Y. Aye, Redox signaling by reactive electrophiles and oxidants. Chemical reviews 118, 8798–8888 (2018).

64. K. N. M. Daeffler, H. A. Lester, D. A. Dougherty, Functionally Important Aromatic–Aromatic and Sulfur−π Interactions in the D2 Dopamine Receptor. Journal of the American Chemical Society 134, 14890–14896 (2012).

65. A. Gora, J. Brezovsky, J. Damborsky, Gates of Enzymes. Chemical Reviews 113, 5871–5923 (2013).

66. D. Li, F. Pucci, M. Rooman, PInteract: Detecting Aromatic-Involving Motifs in Proteins and Protein-Nucleic Acid Complexes. Biomolecules 15, 1204 (2025).

67. A. F. Alpi, V. Chaugule, H. Walden, Mechanism and disease association of E2-conjugating enzymes: lessons from UBE2T and UBE2L3. Biochemical Journal 473, 3401–3419 (2016).

68. T. J. O’Neill, M. J. Tofaute, D. Krappmann, Function and targeting of MALT1 paracaspase in cancer. Cancer Treat Rev 117, 102568 (2023).

69. M. D. Stewart, T. Ritterhoff, R. E. Klevit, P. S. Brzovic, E2 enzymes: more than just middle men. Cell Res 26, 423–440 (2016).

70. M. Pulvino et al., Inhibition of proliferation and survival of diffuse large B-cell lymphoma cells by a small-molecule inhibitor of the ubiquitin-conjugating enzyme Ubc13-Uev1A. Blood 120, 1668–1677 (2012).

71. M. J. Long, X. Liu, Y. Aye, Genie in a bottle: controlled release helps tame natural polypharmacology? Current opinion in chemical biology 51, 48–56 (2019).

72. H. Zhou, M. Q. Du, V. M. Dixit, Constitutive NF-kappaB activation by the t(11;18)(q21;q21) product in MALT lymphoma is linked to deregulated ubiquitin ligase activity. Cancer Cell 7, 425–431 (2005).

73. P. de Bie, A. Ciechanover, Ubiquitination of E3 ligases: self-regulation of the ubiquitin system via proteolytic and non-proteolytic mechanisms. Cell Death & Differentiation 18, 1393–1402 (2011).

74. K.-T. Huang et al., Z-REX: shepherding reactive electrophiles to specific proteins expressed tissue specifically or ubiquitously, and recording the resultant functional electrophile-induced redox responses in larval fish. Nature Protocols, 1–46 (2023).

