## Supplemental information for "MALT1 is an E3 ligase catalyzing its own activation by monoubiquitination"

**List of materials included:**

- 5 supporting figures (Figures S1-S5)
- 3 appendix figures
- legend for supporting table (Table S1)

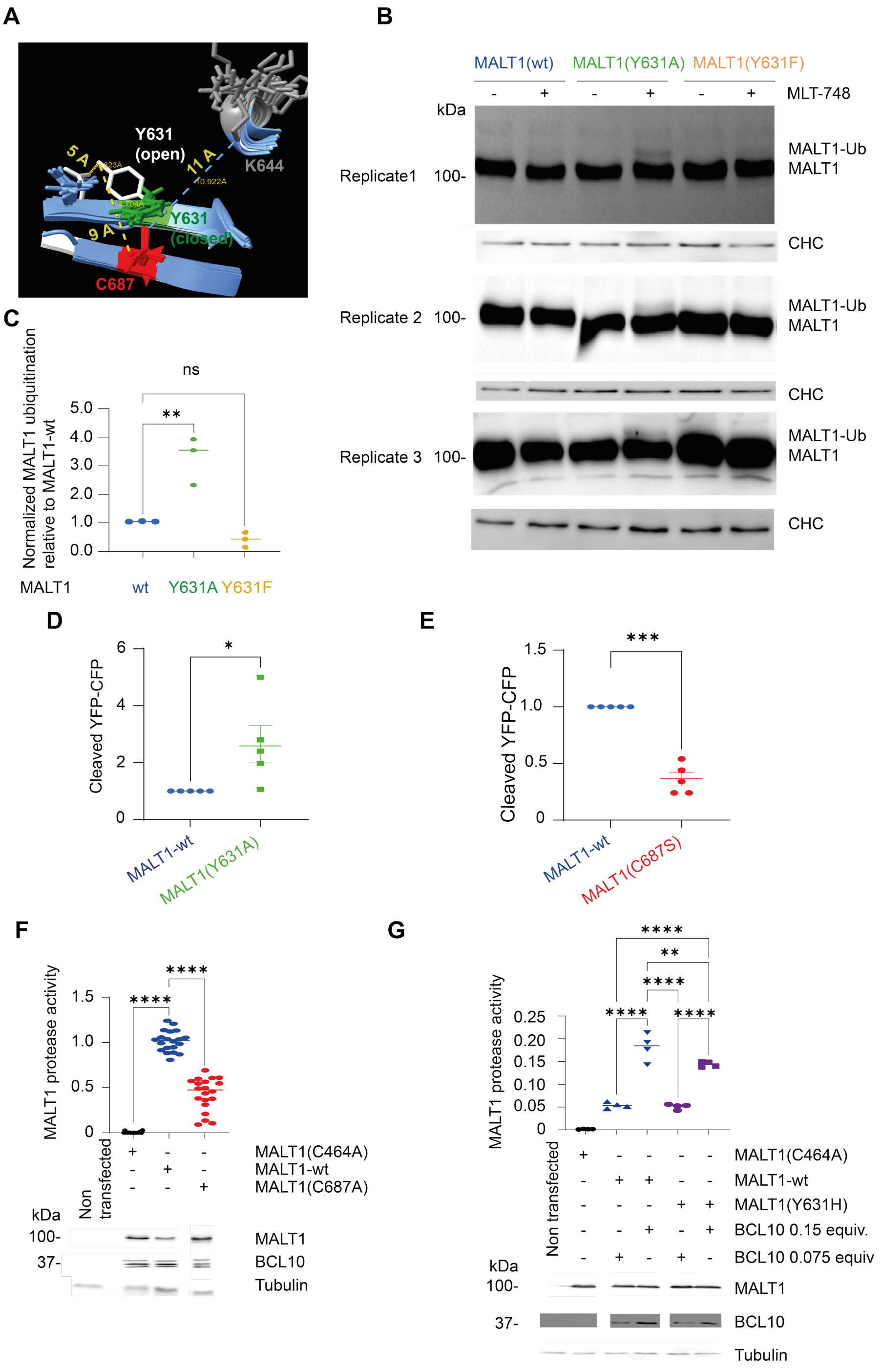


**Figure S1.** ***Y631 is a gatekeeper residue relying on both steric hinderance and its phenolic hydroxyl group to regulate MALT1 protease activity.***

(**A**) Structures from **Appendix Figure S1B** showing only residues in the β-sheet surrounding Y631 and C687 as well as the helical region around K644. One structure, 6H4A (*37*) (white), shows the phenol group of the tyrosyl side chain rotated to allow access of K644 to C687, consistent with a gating function. This appears to happen due to deformation of the β-strand housing Y631, allowing D629 to twist away from Y631. The conformation of the phenol group in white represents the "gate open" conformation. Note the phenolic OH is located over the carboxylate residue. (**B**) HEK293T cells were transfected as shown with either wt-MALT1, MALT1(Y631A), or MALT1(Y631F) together with BCL10 and treated with MLT-748, or DMSO for 24 h. After this time samples were analyzed by western blot as indicated. Triplicate data are shown. (**C**) Quantitation of data in **B**. Data were analyzed by one-way ANOVA followed by Dunnett's test. Clathrin heavy chain (CHC) was used as a loading control (n=3). (**D,E**) HEK293T cells were transfected with the stated MALT1 constructs with BCL10, together with a YFP-LVSR-CFP MALT1 protease activity reporter construct. MALT1 activity was assayed using western blot; quantitation of data in **Figure 2G**. (**F,G**) HEK293T cells were transfected with the stated MALT1 constructs with BCL10, together with a YFP-LVSR-CFP MALT1 protease activity reporter construct. MALT1 activity was assayed using flow cytometry. Western blot shown below (**F**: n=20; **G**: n=4). Data were analyzed by one-way ANOVA, followed by Dunnett's or Tukey's test in accordance with the number of comparisons required; *0.05>p≥0.01; **0.01>p≥0.001; ***0.001>p≥0.0001; ****p<0.0001.

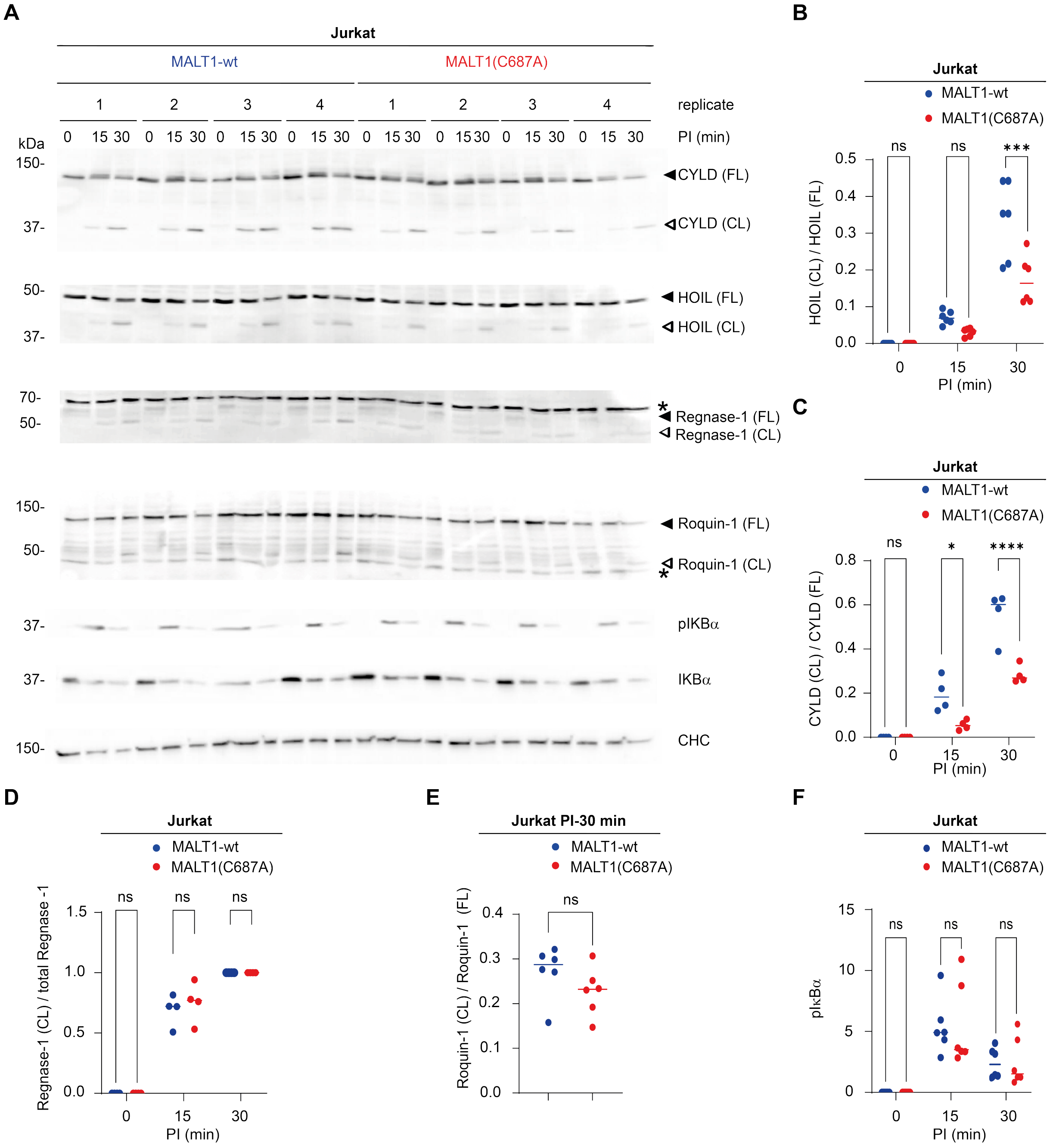


**Figure S2.** ***Cells expressing MALT1(C687A) show suppressed MALT1 protease activity, but normal scaffold functions.***

**A**) MALT1-silenced Jurkat cells expressing MALT1-wt or MALT1(C687A) were PI-stimulated (20 ng/ml PMA, and 1.00 µM ionomycin) for the stated periods and then probed with the stated antibodies by western blot; four independent replicates are shown. (**B-F**) Quantitation of data in **A** for HOIL (n=6), CYLD (n=4), Regnase-1 (n=4), Roquin-1 (n=6), and pIκBα (n=6) respectively. Data analyzed by two-way ANOVA followed by Dunnett's test, except for Roquin-1 cleavage that was not visible in the earliest time point so was analyzed by two-tailed, unpaired *t*-test. Clathrin heavy chain (CHC) was used as a loading control. CL – cleaved; FL – full length. Closed arrows indicate full length protein; open arrows indicate cleaved proteins. □ denotes non-specific band; *0.05>p≥0.01; **0.01>p≥0.001; ***0.001>p≥0.0001; ****p<0.0001.

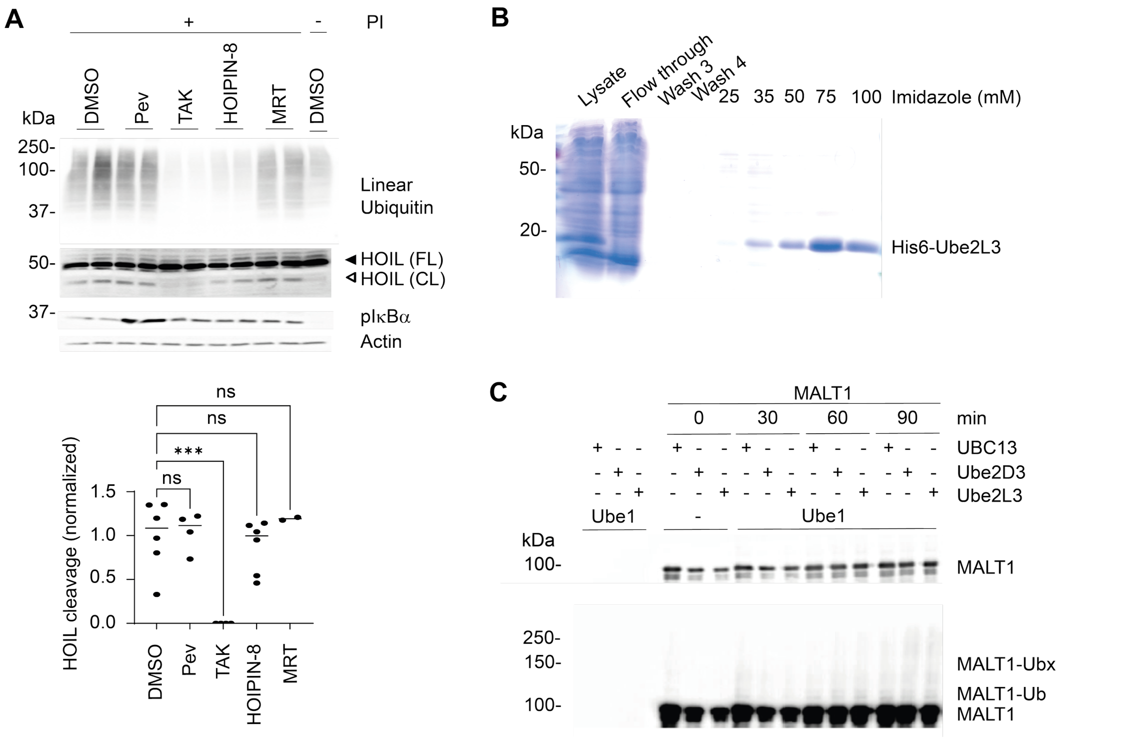


**Figure S3.** ***MALT1 can be ubiquitinated by E2 conjugating enzymes.***

(**A**) The stated Jurkat lines were treated with the stated inhibitor for 3 h, then PI-stimulated (20 ng/ml PMA, and 1.00 µM ionomycin, 20 min); (below) quantification. (**B**) Purification of His$_6$-Ube2L3 using Ni-NTA resin (n=6,4,4,6,2). (**C**) Recombinant purified MALT1 was mixed with the indicated E2 enzymes in the presence of ATP and Ube1 (E1) and incubated at 37 °C for the indicated times. The resulting mixtures were loaded on an SDS-PAGE-gel and analyzed by western blot for MALT1. Pev-pevondistat; TAK-TAK-243; MRT-MRT67307. Actin was used as loading control. CL – cleaved; FL – full length. Closed arrows indicate full length protein; open arrows indicate cleaved proteins. Higher molecular weight species corresponding to polyubiquitinated MALT1 (MALT1-Ubx) and a band likely corresponding to monoubiquitinated MALT1 (MALT1-Ub) are indicated. Data analyzed by one-way ANOVA followed by Dunnetts test; ***0.001>p≥0.0001.

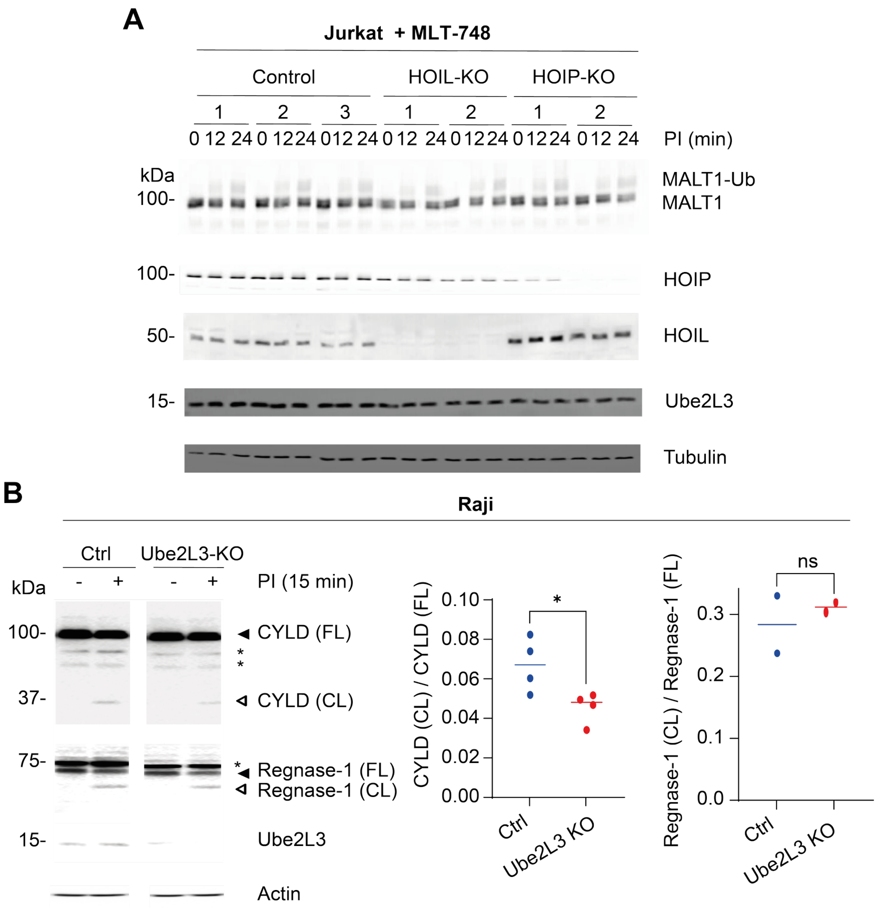


**Figure S4.** ***Inactivation of LUBAC does not affect MALT1 ubiquitination.***

(**A**) The stated Jurkat KO lines were PI-stimulated as described and then analyzed by western blot as described. (**B**) The stated Raji cells were PI-stimulated (15 min) (20 ng/ml PMA, and 1.00 µM ionomycin) then CYLD cleavage was assessed by western blot (n=4,2). Actin was used as a loading control. CL – cleaved; FL – full length. Closed arrows indicate full length protein; open arrows indicate cleaved proteins. Data were analyzed by a two-tailed unpaired t-test. ▯ denotes non-specific band; *0.05>p≥0.01.

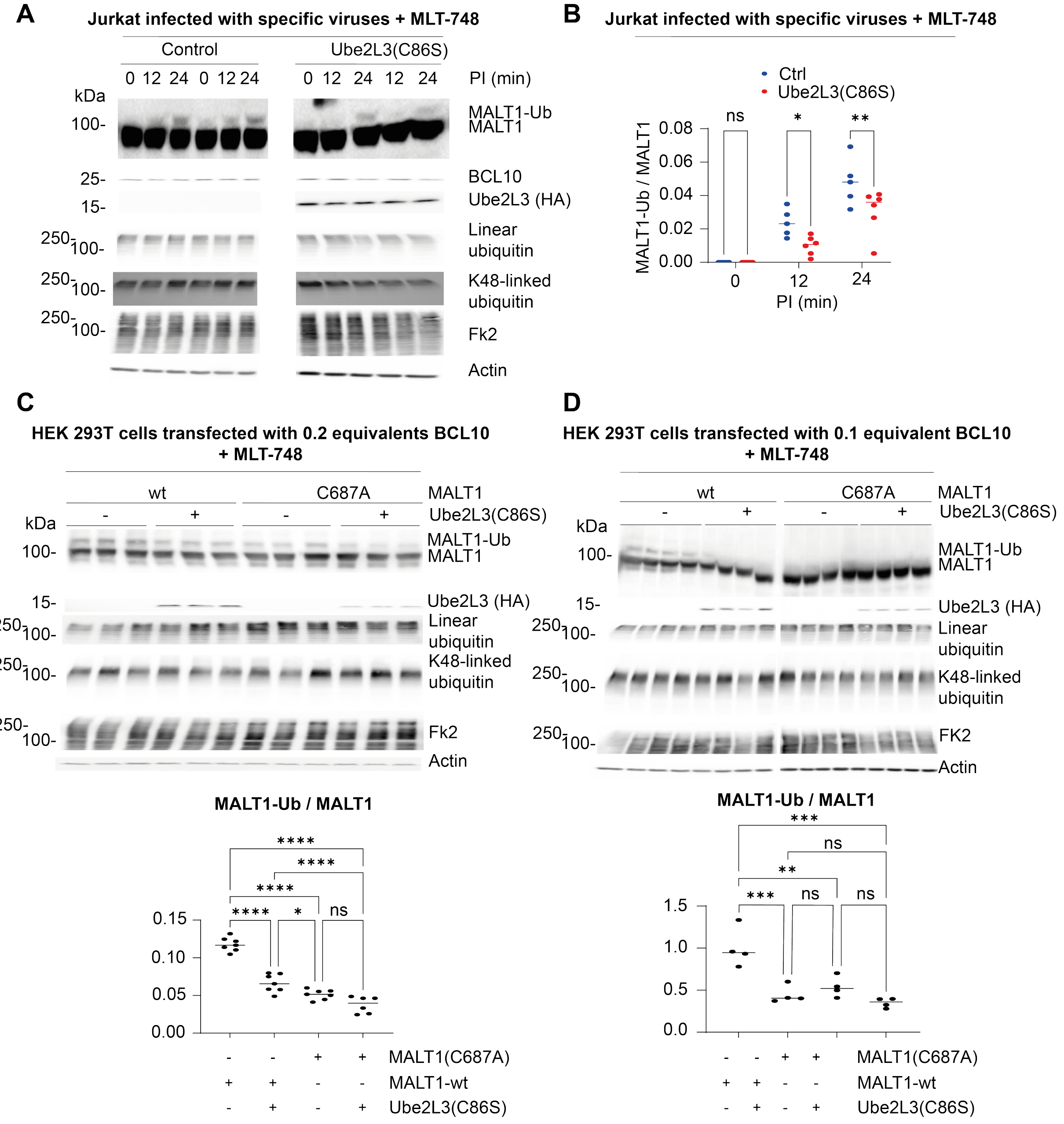


**Figure S5.** ***Interfering with MALT1-ubiquitination apparatus renders MALT1 resistant to ubiquitination changes induced by Ube2L3(C86S).***

(**A**) Jurkat cells were infected as described, treated with MLT-748 (10 µM) and then PI-stimulated for the stated time points. Note the 0 time point for the last data set in Ube2L3 expressing cells was omitted due to lane limitations. (**B**) quantitation of data in **A** (n=5,6). (**C**) HEK293T cells were transfected as described (0.2 equivalents of BCL10), treated with MLT-748 (10 µM) and then MALT1 ubiquitination was assessed by western blot; below quantitation (n=7). (**D**) Same as **C**, but 0.1 equivalents of BCL10 were used; below quantitation (n=4). FK2 detects mono- and polyubiquitinated proteins. Data were analyzed by one or two-way ANOVA followed by Tukey's or Dunnett's test as appropriate for the number of comparisons; *0.05>p≥0.01; **0.01>p≥0.001; ***0.001>p≥0.0001; ****p<0.0001.

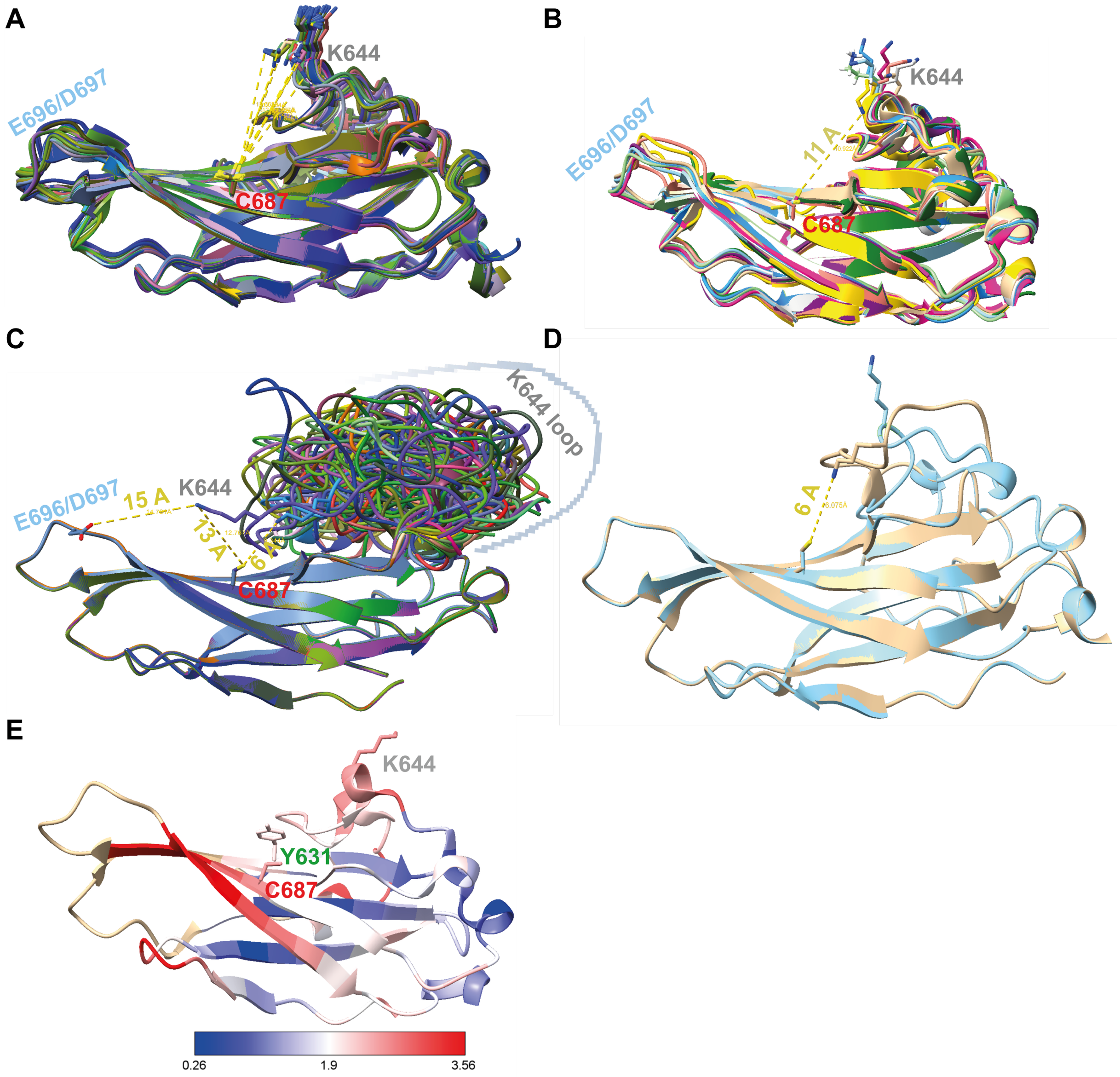


**Appendix Figure S1.** ***C687 is an ideal candidate residue for accepting ubiquitin from UBS1 and transferring it to K644.***

(**A**) The Ig3 domains of the top 40 AlphaFold structures of MALT1 were superimposed in ChimeraX and then for the closest C687 and K644 residues rotamers were displayed using the Dunbrack method; for these rotamers, the distances between the sulfur and terminal amine atoms were calculated. (**B**) Same analysis as in **A** for structures in PDB. (**C**) PDB structure of MALT1 3uoa was analyzed for K644 loop flexibility using the AFflecto package in MoMA (generating 100 conformations). The Ig3 domain was selected and matchmaker program in ChimeraX was used to superimpose these conformations. Then side chain conformations were added using the Dunbrack method. Relevant distances are shown only. (**D**) The standard conformation of the Ig3 domain (periwinkle) and the conformation with lysine closest to C687 (fawn) are shown. Note a C-S single bond is close to 2 Å in length. **(E)** Rendering of root mean square deviation of backbone NH residues in the MALT1 Ig3 domain between in 3V55 and 7PAV.

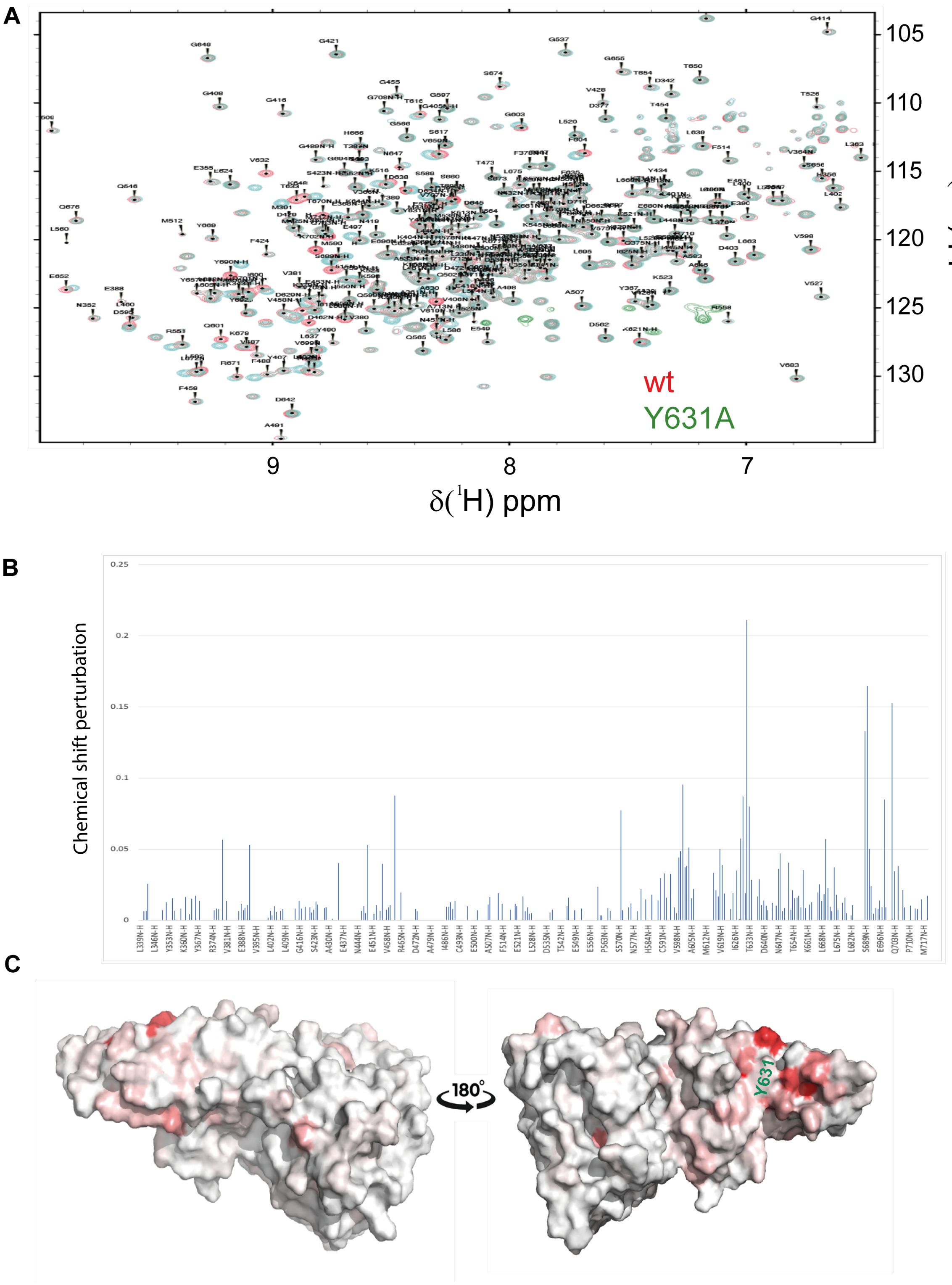


**Appendix Figure S2.** ***NMR of MALT1(Y631A) shows it is structurally similar to MALT1(wt).***

(**A**) TROSY HSQC-NMR of uniformly $^{15}N$ labelled MALT1 for both MALT1-wt (red) and MALT1(Y631A) (green) with peak assignments. (**B**) Chemical shift perturbations. (**C**) Surface representation of MALT1 showing largest perturbations upon Y631A mutation.

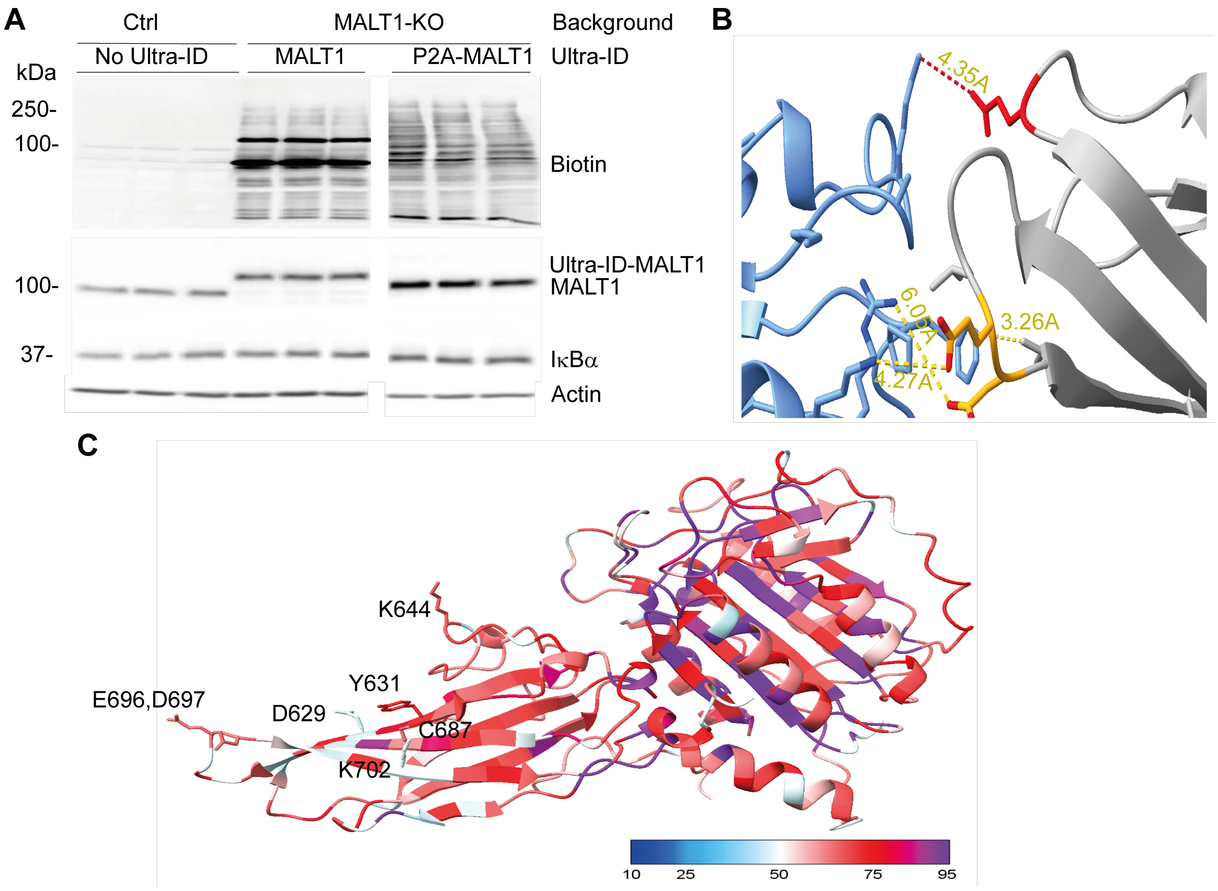


**Appendix Figure S3.** ***ultraID-MALT1 identifies Ube2L3 as a MALT1 associating enzyme; docking shows Ube2L3 can bind UBS1 in a similar way to how it binds HOIP.***

(**A**) MALT1-silenced Jurkat cells were infected with ultraID-MALT1 and ultraID-P2A-MALT1. These lines as well as a Ctrl line were treated with Biotin (200 μM) and then lysates were analyzed by western blotting as described. (**B**) Docking of Ube2L3(blue) with MALT1 (grey) showing the interaction interface; MALT1-UBS1 is shown in orange; E624 is shown in red. Actin was used as a loading control. (**C**) Sequences of 300 MALT1s from mammals, reptiles, birds, and nematodes were aligned and identity was plotted over the Alphafold structure of the caspase and Ig3 domain.

**Legend for supporting Table S1**

**Table S1. Comparison of MALT1 interactomes identified post streptavidin enrichment in MALT1-KO cells, MALT1-KO cells rescued with ultraID-MALT1, and MALT1-KO cells rescued with ultraID-P2A-MALT1.**

MALT1-deficient Jurkat lines were reconstituted separately with ultraID-MALT1 or ultraID-P2A-MALT1 constructs. Cells were grown in biotin-free medium and biotin was added for 18 h prior to lysis. Interactomes were enriched using streptavidin magnetic beads and identified by mass spectrometry. Each set was performed in triplicate, and data were compared between ultraID-MALT1 and ultraID-P2A-MALT1 expressing cells. Proteins significantly enriched in cells expressing ultraID-MALT1 versus ultraID-P2A-MALT1 were considered positive hits.
