## Supplemental Table 1 for "MALT1 is an E3 ligase catalyzing its own activation by monoubiquitination"

**Table S1. Comparison of MALT1 interactomes identified post streptavidin enrichment in MALT1-KO cells, MALT1-KO cells rescued with ultraID-MALT1, and MALT1-KO cells rescued with ultraID-P2A-MALT1.**

Italics indicates hit ultraID-MALT1 versus ultraID-P2A-MALT1; red indicates hit in each specific comparison; purple indicates hit in ultraID-MALT1 versus ultraID-P2A-MALT1 but not hit in ultraID-Malt1 versus control, or positive hit in ultraID-P2A-MALT1 versus control

Software version: MaxQuant 2.5.1.0
RefProt_Homo_sapiens_20240214
Universal_Contaminant_Protein
Samples were digested following the standard protocol for Turbo-ID labeling.

MS analysis was performed on Fusion instrument equipped with a FAIMS source.

| 2 | Gene.names | Peptides | Razor.unique.peptides | significant. MALT1 (wt)-Control | significant.MALT1 (wt)-UltraID-P2A-MALT1-wt |
|---|---|---|---|---|---|
| *Q7Z739;A0A024R7* | *YTHDF3* | *7* | *3* | *TRUE* | *TRUE* |
| O95999;A0A087W\ | BCL10 | 8 | 8 | TRUE | TRUE |
| *Q9NP79;A0A087W* | *VTA1* | *4* | *4* | *TRUE* | *TRUE* |
| *Q04206;Q2TAM5;A* | *RELA* | *6* | *6* | *TRUE* | *TRUE* |

| | | | | | |
|---|---|---|---|---|---|
| A0A096LP25;Q2M2 | AAK1 | 9 | 9 | TRUE | TRUE |
| Q53H96;B5MD87;A | PYCRL | 2 | 2 | TRUE | TRUE |
| Q9NR56;C9JP00;A | MBNL1 | 2 | 2 | TRUE | TRUE |
| Q8IZP0;B6VEX4;A0 | ABI1 | 5 | 5 | FALSE | TRUE |
| Q13617;A0A0A0M7 | CUL2 | 19 | 19 | TRUE | TRUE |
| Q58A45;A0A0C4DI | PAN3 | 8 | 8 | TRUE | TRUE |
| Q9Y2X7;J3QRU8;A | GIT1 | 5 | 5 | TRUE | TRUE |
| O00571;A0A2R8Y4 | DDX3X;DDX3Y | 18 | 18 | TRUE | TRUE |
| Q8N6H7;E9PQP3;A | ARFGAP2 | 4 | 4 | TRUE | TRUE |
| Q8N1K5;A0A0D9SI | THEMIS | 7 | 7 | TRUE | TRUE |
| Q9Y4K1;A0A0J9YW | AIM1 | 11 | 11 | TRUE | TRUE |
| Q6P3W7;A0A0U1R | SCYL2 | 15 | 15 | TRUE | TRUE |
| P42226;A0A494C0 | STAT6 | 3 | 3 | TRUE | TRUE |
| Q99700;H0YH87;A | ATXN2 | 5 | 5 | FALSE | TRUE |
| P67936;A0A2R8YG | TPM4 | 5 | 4 | TRUE | TRUE |
| Q9UHD2;A0A7I2V6 | TBK1 | 5 | 5 | TRUE | TRUE |
| Q92538;A0A669KB | GBF1 | 30 | 30 | TRUE | TRUE |
| Q99615;A0A6I8PU | DNAJC7 | 4 | 4 | TRUE | TRUE |
| Q96DH6;B4DHE8;A | MSI2;MSI1 | 2 | 2 | TRUE | TRUE |
| P49588;H3BPK7;A | AARS | 17 | 17 | TRUE | TRUE |
| P50747;A0A6Q8PF | HLCS | 13 | 13 | FALSE | TRUE |
| Q9Y223;A0A7I2SU | GNE | 9 | 9 | TRUE | TRUE |
| E9PAV3;F8VZJ2;F8 | NACA | 2 | 2 | TRUE | TRUE |

| | | | | | |
|---|---|---|---|---|---|
| Q13561;F8VW18;A | DCTN2 | 2 | 2 | TRUE | TRUE |
| P52732;A0A7I2V3A | KIF11 | 8 | 8 | TRUE | TRUE |
| P20290;A0A7I2YQI | BTF3 | 2 | 2 | TRUE | TRUE |
| Q15032;A0A804HI. | R3HDM1 | 7 | 7 | TRUE | TRUE |
| O00170;A0A804Hk | AIP | 8 | 8 | TRUE | TRUE |
| O95260;A0A8I5KVI | ATE1 | 6 | 6 | TRUE | TRUE |
| O75534;A0A8I5QJ/ | CSDE1 | 7 | 7 | TRUE | TRUE |
| Q15149;A0A8I5KSI | PLEC | 9 | 9 | FALSE | TRUE |
| Q92540;E9PD50;A | SMG7 | 16 | 16 | TRUE | TRUE |
| P13804;H0YKF0;H: | ETFA | 2 | 2 | FALSE | TRUE |
| Q9UKY7;A0A8I5KX | CDV3 | 5 | 5 | TRUE | TRUE |
| Q6ULP2;A0A8Q3W | AFTPH | 11 | 11 | TRUE | TRUE |
| Q7Z3B4;A0A8Q3W | NUP54 | 7 | 7 | TRUE | TRUE |
| Q9BVL2;A0A8Q3W | NUPL1 | 7 | 7 | TRUE | TRUE |
| A0A8V8TML6;H0YA | ERBB2IP | 18 | 18 | TRUE | TRUE |
| Q96RU3;A0A994J3 | FNBP1 | 6 | 6 | TRUE | TRUE |
| P48556;R4GMR5;A | PSMD8 | 5 | 5 | TRUE | TRUE |
| Q92900;A0A994J4 | UPF1 | 6 | 6 | TRUE | TRUE |
| Q9HAT8;A0A994J4 | PELI2 | 3 | 3 | TRUE | TRUE |
| Q9Y6Y8;A0A994J5 | SEC23IP | 8 | 8 | TRUE | TRUE |
| P42566;A0A994J5J | EPS15 | 14 | 14 | TRUE | TRUE |
| A5PLN9 | TRAPPC13 | 3 | 3 | TRUE | TRUE |
| A6NKD9;H0YJE0;G | CCDC85C | 6 | 6 | TRUE | TRUE |

| | | | | | |
|---|---|---|---|---|---|
| O43768;A6NMQ3;( | ENSA;ARPP19 | 2 | 2 | TRUE | TRUE |
| Q08257;C9JH92;A( | CRYZ | 5 | 5 | TRUE | TRUE |
| O15372;B3KS98;E! | EIF3H | 6 | 6 | TRUE | TRUE |
| Q99439;B4DDF4;B | CNN2 | 9 | 9 | TRUE | TRUE |
| Q13610;B4DJV5 | PWP1 | 3 | 3 | FALSE | TRUE |
| Q9NR46;B7ZC38;B | SH3GLB2 | 10 | 10 | TRUE | TRUE |
| O60573;B9A044;B | EIF4E2 | 3 | 3 | TRUE | TRUE |
| P30876;C9J2Y9;C! | POLR2B | 13 | 13 | FALSE | TRUE |
| C9J6P4;Q7Z2W4;A | ZC3HAV1 | 25 | 25 | TRUE | TRUE |
| P05166;E9PDR0;C | PCCB | 16 | 16 | FALSE | TRUE |
| P35998;C9JX88 | PSMC2 | 10 | 10 | TRUE | TRUE |
| P78344;D3DQV9;H | EIF4G2 | 15 | 15 | TRUE | TRUE |
| P46926;D6RAY7;D | GNPDA1 | 4 | 4 | TRUE | TRUE |
| P19838;D6RH30;A | NFKB1 | 18 | 18 | TRUE | TRUE |
| Q93045;E5RGX5 | STMN2 | 2 | 2 | TRUE | TRUE |
| P50914;E7EPB3 | RPL14 | 3 | 3 | FALSE | TRUE |
| Q12768;E7EQI7 | KIAA0196 | 10 | 10 | TRUE | TRUE |
| Q08209;E7ETC2;P: | PPP3CA;PPP3CB | 3 | 3 | TRUE | TRUE |
| Q9BWF3;E9PB51;I | RBM4 | 5 | 5 | TRUE | TRUE |
| Q5H9R7;H7BXH2;I | PPP6R3 | 15 | 15 | TRUE | TRUE |
| Q96K17;E9PL10 | BTF3L4 | 3 | 3 | TRUE | TRUE |
| E9PN18;E9PQ56;Q | PUF60 | 3 | 3 | TRUE | TRUE |
| Q9Y296;E9PQE8;E | TRAPPC4 | 2 | 2 | TRUE | TRUE |

| | | | | | |
|---|---|---|---|---|---|
| Q9UK41;E9PQR7;E | VPS28 | 3 | 3 | TRUE | TRUE |
| Q9H4A3;F5GWT4;A | WNK1 | 11 | 11 | TRUE | TRUE |
| Q9NRL3;F5GYK2;M | STRN4 | 4 | 3 | TRUE | TRUE |
| Q9NVZ3;F6SKB8;D | NECAP2 | 4 | 4 | TRUE | TRUE |
| P40616;F8VYN9;F8 | ARL1 | 2 | 2 | TRUE | TRUE |
| Q9BY44;F8WAE5 | EIF2A | 5 | 5 | TRUE | TRUE |
| Q8WW33;G3V1V4 | GTSF1 | 3 | 3 | TRUE | TRUE |
| Q9Y4F3;G8JLP4;F8 | KIAA0430 | 6 | 6 | TRUE | TRUE |
| Q15942;H0Y2Y8 | ZYX | 11 | 11 | TRUE | TRUE |
| P35606;H0Y938;H0 | COPB2 | 10 | 10 | TRUE | TRUE |
| H0YC42;H7C5V3 | MRPS28 | 4 | 4 | TRUE | TRUE |
| O00303;H0YDT6;A | EIF3F | 13 | 13 | TRUE | TRUE |
| P53041;H0YDU8 | PPP5C | 9 | 9 | TRUE | TRUE |
| H0YEF7;H0YD48 | PICALM | 4 | 2 | TRUE | TRUE |
| Q04726;H0YNT2;H | TLE3 | 3 | 3 | TRUE | TRUE |
| P25789;H0YLC2;H | PSMA4 | 2 | 2 | FALSE | TRUE |
| O75179;H0YM23 | ANKRD17 | 24 | 14 | TRUE | TRUE |
| O00161;H3BNE1;H | SNAP23 | 3 | 3 | TRUE | TRUE |
| P51159;H3BS49;H | RAB27A | 4 | 4 | TRUE | TRUE |
| Q86UK7;H3BPG6 | ZNF598 | 12 | 12 | TRUE | TRUE |
| Q96CD2;H3BSE3;H | PPCDC | 6 | 6 | TRUE | TRUE |
| O14562;H3BRL3 | UBFD1 | 2 | 2 | TRUE | TRUE |
| Q12968;H3BRS1;I3 | NFATC3 | 13 | 13 | TRUE | TRUE |

| | | | | | |
|---|---|---|---|---|---|
| P29016;H7C0I2 | CD1B | 2 | 2 | TRUE | TRUE |
| P52565;J3QQX2;J3 | ARHGDIA | 3 | 3 | FALSE | TRUE |
| O14950;P19105;J3 | MYL12B;MYL12A | 3 | 3 | FALSE | TRUE |
| P42694;J3QS41;A0 | HELZ | 22 | 22 | TRUE | TRUE |
| Q08117;K7ELE3;K7 | AES | 3 | 3 | TRUE | TRUE |
| O14737;K7EQA1 | PDCD5 | 2 | 2 | TRUE | TRUE |
| O00463 | TRAF5 | 14 | 14 | TRUE | TRUE |
| O00483 | NDUFA4 | 2 | 2 | FALSE | TRUE |
| O00487 | PSMD14 | 3 | 3 | TRUE | TRUE |
| O00628;A0A7I2V2V | PEX7 | 6 | 6 | FALSE | TRUE |
| O00743 | PPP6C | 2 | 2 | TRUE | TRUE |
| O14828 | SCAMP3 | 3 | 3 | TRUE | TRUE |
| O15027;F1T0I1;A0. | SEC16A | 31 | 31 | TRUE | TRUE |
| O15212;A2AB88 | PFDN6 | 3 | 3 | TRUE | TRUE |
| O15355 | PPM1G | 9 | 9 | TRUE | TRUE |
| O43237;J3KRI4;B4I | DYNC1LI2 | 4 | 3 | TRUE | TRUE |
| O60496 | DOK2 | 9 | 9 | TRUE | TRUE |
| O60645;D6RB59 | EXOC3 | 9 | 9 | TRUE | TRUE |
| O60664;K7ERZ3;K7 | PLIN3 | 5 | 5 | TRUE | TRUE |
| O75113 | N4BP1 | 3 | 3 | TRUE | TRUE |
| O75143;J9JIF6;E9P | ATG13 | 6 | 6 | FALSE | TRUE |
| O75420 | GIGYF1 | 7 | 7 | TRUE | TRUE |
| O75884 | RBBP9 | 2 | 2 | TRUE | TRUE |

| | | | | | |
|---|---|---|---|---|---|
| O76003 | GLRX3 | 4 | 4 | TRUE | TRUE |
| O76071 | CIAO1 | 5 | 5 | TRUE | TRUE |
| O94804 | STK10 | 8 | 7 | TRUE | TRUE |
| O95487 | SEC24B | 6 | 6 | TRUE | TRUE |
| O95721 | SNAP29 | 5 | 5 | TRUE | TRUE |
| O95757;E9PDE8;D | HSPA4L | 10 | 9 | TRUE | TRUE |
| O95793;Q5JW30 | STAU1 | 7 | 7 | TRUE | TRUE |
| O95801 | TTC4 | 11 | 11 | TRUE | TRUE |
| P00558 | PGK1 | 11 | 11 | FALSE | TRUE |
| P04083;Q5T3N1 | ANXA1 | 15 | 15 | TRUE | TRUE |
| P08758;D6RBE9;D | ANXA5 | 8 | 8 | TRUE | TRUE |
| P11021;A0A7P0TA | HSPA5 | 16 | 15 | FALSE | TRUE |
| P11142;E9PKE3;E9 | HSPA8 | 26 | 26 | TRUE | TRUE |
| P11498;A0A494C0 | PC | 48 | 48 | FALSE | TRUE |
| P13639 | EEF2 | 23 | 23 | FALSE | TRUE |
| P14317;E7EVW7 | HCLS1 | 14 | 14 | TRUE | TRUE |
| P15311 | EZR | 14 | 8 | TRUE | TRUE |
| P15531;E7ERL0;J3 | NME1;NME2;NME | 5 | 5 | FALSE | TRUE |
| P17987;E7EQR6;E | TCP1 | 28 | 28 | TRUE | TRUE |
| P21281;H0YC04 | ATP6V1B2 | 3 | 3 | TRUE | TRUE |
| P21333;Q60FE5;A( | FLNA | 63 | 63 | TRUE | TRUE |
| P22234;E9PBS1;D( | PAICS | 10 | 10 | TRUE | TRUE |
| P23634 | ATP2B4 | 3 | 3 | FALSE | TRUE |

| | | | | | |
|---|---|---|---|---|---|
| *P25685;M0QXK0;A* | *DNAJB1* | *9* | *9* | *TRUE* | *TRUE* |
| *P26196* | *DDX6* | *8* | *8* | *TRUE* | *TRUE* |
| *P29558;E7ETU5* | *RBMS1* | *5* | *5* | *TRUE* | *TRUE* |
| P29692;E9PRY8;E9 | EEF1D | 10 | 10 | FALSE | TRUE |
| P30041 | PRDX6 | 11 | 11 | FALSE | TRUE |
| *P30086* | *PEBP1* | *3* | *3* | *TRUE* | *TRUE* |
| *P31948* | *STIP1* | *23* | *23* | *TRUE* | *TRUE* |
| *P37802;X6RJP6* | *TAGLN2* | *9* | *9* | *TRUE* | *TRUE* |
| P37837;F2Z393 | TALDO1 | 10 | 10 | TRUE | TRUE |
| *P38606;A0A994J73* | *ATP6V1A* | *9* | *9* | *TRUE* | *TRUE* |
| *P46736;X6RJS7* | *BRCC3* | *5* | *5* | *TRUE* | *TRUE* |
| *P46937;H0YCI3* | *YAP1* | *4* | *4* | *TRUE* | *TRUE* |
| P47929 | LGALS7 | 6 | 6 | FALSE | TRUE |
| *P48643;B7ZAR1;E9* | *CCT5* | *22* | *22* | *TRUE* | *TRUE* |
| *P50990;H7C4C8* | *CCT8* | *24* | *24* | *TRUE* | *TRUE* |
| *P55010* | *EIF5* | *3* | *3* | *TRUE* | *TRUE* |
| *P57088;D6RAA6* | *TMEM33* | *4* | *4* | *FALSE* | *TRUE* |
| *P62424;Q5T8U3* | *RPL7A* | *4* | *4* | *FALSE* | *TRUE* |
| *P68036;A0A8Q3W* | *UBE2L3* | *3* | *3* | *TRUE* | *TRUE* |
| *P78318* | *IGBP1* | *4* | *4* | *TRUE* | *TRUE* |
| P78527 | PRKDC | 19 | 19 | FALSE | TRUE |
| P86791;P86790 | CCZ1;CCZ1B | 21 | 21 | TRUE | TRUE |
| *P98170;B1AKU2* | *XIAP* | *4* | *4* | *TRUE* | *TRUE* |

| | | | | | |
|---|---|---|---|---|---|
| Q00013 | MPP1 | 11 | 11 | TRUE | TRUE |
| Q01085;E7ETJ9;A6 | TIAL1 | 7 | 7 | TRUE | TRUE |
| Q01196;C9JWM1;/ | RUNX1 | 3 | 3 | TRUE | TRUE |
| Q01850;H3BTR1;H | CDR2 | 2 | 2 | TRUE | TRUE |
| Q02790 | FKBP4 | 9 | 9 | FALSE | TRUE |
| Q06210;A0A6I8PRI | GFPT1 | 10 | 10 | TRUE | TRUE |
| Q06323 | PSME1 | 6 | 6 | TRUE | TRUE |
| Q08752 | PPID | 13 | 13 | TRUE | TRUE |
| Q13033 | STRN3 | 10 | 10 | TRUE | TRUE |
| Q13043;A0A8V8TL | STK4 | 4 | 4 | FALSE | TRUE |
| Q13085 | ACACA | 100 | 100 | FALSE | TRUE |
| Q13151 | HNRNPA0 | 3 | 3 | FALSE | TRUE |
| Q13371 | PDCL | 5 | 5 | TRUE | TRUE |
| Q13492 | PICALM | 8 | 8 | TRUE | TRUE |
| Q13546 | RIPK1 | 7 | 7 | TRUE | TRUE |
| Q13596;H0YK42 | SNX1 | 9 | 9 | TRUE | TRUE |
| Q14247 | CTTN | 5 | 5 | TRUE | TRUE |
| Q14671;Q5T1Z4;Q. | PUM1 | 16 | 16 | TRUE | TRUE |
| Q14677 | CLINT1 | 9 | 9 | TRUE | TRUE |
| Q15008;C9J7B7;C! | PSMD6 | 7 | 7 | TRUE | TRUE |
| Q15018 | FAM175B | 6 | 6 | TRUE | TRUE |
| Q15052 | ARHGEF6 | 8 | 8 | TRUE | TRUE |
| Q15274 | QPRT | 5 | 5 | TRUE | TRUE |

| | | | | | |
|---|---|---|---|---|---|
| Q15365 | PCBP1 | 12 | 12 | TRUE | TRUE |
| Q15654 | TRIP6 | 5 | 5 | TRUE | TRUE |
| Q16543;K7EQA9;K | CDC37 | 8 | 8 | TRUE | TRUE |
| Q16643;D6R9Q9 | DBN1 | 3 | 3 | TRUE | TRUE |
| Q4G0X4 | KCTD21 | 3 | 3 | TRUE | TRUE |
| Q53ET0;Q5T4K5;H | CRTC2 | 4 | 4 | TRUE | TRUE |
| Q5D1E8;A0A1W2P | ZC3H12A | 4 | 4 | TRUE | TRUE |
| Q5MNZ9;K7ELY0;G | WIPI1 | 2 | 2 | TRUE | TRUE |
| Q6P2E9 | EDC4 | 25 | 25 | TRUE | TRUE |
| Q6P2H3 | CEP85 | 16 | 16 | TRUE | TRUE |
| Q6PKG0;A0A8I5KW | LARP1 | 17 | 17 | TRUE | TRUE |
| Q709C8 | VPS13C | 116 | 116 | TRUE | TRUE |
| Q70EK8 | USP53 | 4 | 4 | FALSE | TRUE |
| Q71RC2;X6RLN4 | LARP4 | 5 | 5 | TRUE | TRUE |
| Q7Z417 | NUFIP2 | 11 | 11 | TRUE | TRUE |
| Q7Z434 | MAVS | 6 | 6 | TRUE | TRUE |
| Q7Z4H7;Q5VY60 | HAUS6 | 24 | 24 | TRUE | TRUE |
| Q7Z5R6 | APBB1IP | 7 | 7 | TRUE | TRUE |
| Q86TB9 | PATL1 | 17 | 17 | TRUE | TRUE |
| Q86WR0;G3V121;E | CCDC25 | 2 | 2 | TRUE | TRUE |
| Q96MW1;Q86WV7 | CCDC43 | 3 | 3 | FALSE | TRUE |
| Q86YV0 | RASAL3 | 20 | 20 | TRUE | TRUE |
| Q86YV5 | SGK223 | 9 | 9 | TRUE | TRUE |

| | | | | | |
|---|---|---|---|---|---|
| *Q8IVD9* | *NUDCD3* | *8* | *8* | *TRUE* | *TRUE* |
| *Q8IWD4;B0QYH3* | *CCDC117* | *4* | *4* | *TRUE* | *TRUE* |
| *Q8IWZ3;E9PDP5* | *ANKHD1* | *32* | *32* | *TRUE* | *TRUE* |
| *Q8IX90* | *SKA3* | *2* | *2* | *TRUE* | *TRUE* |
| *Q8IXJ6;A0A0A0MR* | *SIRT2* | *9* | *9* | *TRUE* | *TRUE* |
| *Q8IZD4;F5GZK9* | *DCP1B* | *12* | *12* | *TRUE* | *TRUE* |
| *Q8IZH2;H7C5E4* | *XRN1* | *17* | *17* | *TRUE* | *TRUE* |
| *Q8N5G2;A0A3B3IT* | *TMEM57* | *6* | *6* | *TRUE* | *TRUE* |
| *Q8N5M4;H0YDR3;* | *TTC9C* | *3* | *3* | *TRUE* | *TRUE* |
| *Q8N6T3;E5RHC5;E* | *ARFGAP1* | *6* | *6* | *TRUE* | *TRUE* |
| *Q8NDV7;H7C269* | *TNRC6A* | *18* | *18* | *TRUE* | *TRUE* |
| *Q8NHV4* | *NEDD1* | *6* | *6* | *TRUE* | *TRUE* |
| *Q8TC07* | *TBC1D15* | *8* | *8* | *TRUE* | *TRUE* |
| *Q8TCG1* | *KIAA1524* | *14* | *14* | *TRUE* | *TRUE* |
| *Q8WWM7* | *ATXN2L* | *27* | *27* | *TRUE* | *TRUE* |
| *Q92572;F5H459* | *AP3S1* | *3* | *3* | *TRUE* | *TRUE* |
| *Q92783;O75886;A* | *STAM;STAM2* | *2* | *2* | *TRUE* | *TRUE* |
| *Q92905;E5RHH5* | *COPS5* | *6* | *6* | *TRUE* | *TRUE* |
| *Q96BY7* | *ATG2B* | *33* | *33* | *TRUE* | *TRUE* |
| *Q96CN9* | *GCC1* | *18* | *18* | *TRUE* | *TRUE* |
| Q96EA4 | SPDL1 | 7 | 7 | FALSE | TRUE |
| *Q96H20;H0Y8S5;D* | *SNF8* | *2* | *2* | *TRUE* | *TRUE* |
| *Q96JH7* | *VCPIP1* | *22* | *22* | *TRUE* | *TRUE* |

| | | | | | |
|---|---|---|---|---|---|
| Q96K76 | USP47 | 22 | 22 | TRUE | TRUE |
| Q96KP4 | CNDP2 | 5 | 5 | TRUE | TRUE |
| Q99808 | SLC29A1 | 2 | 2 | TRUE | TRUE |
| Q9BSJ8 | ESYT1 | 6 | 6 | TRUE | TRUE |
| Q9BUT9 | FAM195A | 2 | 2 | TRUE | TRUE |
| Q9H2G2 | SLK | 16 | 16 | TRUE | TRUE |
| Q9H2J4 | PDCL3 | 6 | 6 | TRUE | TRUE |
| Q9H3S7 | PTPN23 | 30 | 30 | TRUE | TRUE |
| Q9H444 | CHMP4B | 2 | 2 | TRUE | TRUE |
| Q9H467 | CUEDC2 | 4 | 4 | TRUE | TRUE |
| Q9H6S1 | AZI2 | 3 | 3 | FALSE | TRUE |
| Q9H6T3 | RPAP3 | 9 | 9 | TRUE | TRUE |
| Q9H6Z4;B7Z7F3;K | RANBP3 | 4 | 4 | FALSE | TRUE |
| Q9HB71 | CACYBP | 13 | 13 | TRUE | TRUE |
| Q9HBK9 | AS3MT | 4 | 4 | TRUE | TRUE |
| Q9HCJ0;A0A1B0G | TNRC6C | 17 | 16 | TRUE | TRUE |
| Q9HD26;A0A0J9Y | GOPC | 3 | 3 | TRUE | TRUE |
| Q9NQR4;H7C579 | NIT2 | 6 | 6 | TRUE | TRUE |
| Q9NT62;F8WDI0 | ATG3 | 3 | 3 | TRUE | TRUE |
| Q9NTZ6 | RBM12 | 4 | 4 | TRUE | TRUE |
| Q9NUJ3;E9PJ55 | TCP11L1 | 6 | 6 | TRUE | TRUE |
| Q9NVH1;A0A8I5KX | DNAJC11 | 2 | 2 | TRUE | TRUE |
| Q9NVX0;H3BQU2; | HAUS2 | 4 | 4 | TRUE | TRUE |

| | | | | | |
|---|---|---|---|---|---|
| Q9NYP7;A0A0A0M | ELOVL5 | 3 | 3 | FALSE | TRUE |
| *Q9NZQ3;C9JSC3* | *NCKIPSD* | *9* | *9* | *TRUE* | *TRUE* |
| *Q9UBC2;M0R165;I* | *EPS15L1* | *20* | *20* | *TRUE* | *TRUE* |
| Q9UDY8;A0A3B3IS | MALT1 | 30 | 30 | TRUE | TRUE |
| Q9UGP4;C9JRJ5 | LIMD1 | 20 | 20 | TRUE | TRUE |
| *Q9UKX7* | *NUP50* | *7* | *7* | *TRUE* | *TRUE* |
| *Q9ULE6;A0A8V8TL* | *PALD1* | *13* | *13* | *TRUE* | *TRUE* |
| *Q9ULV4;B4E3S0* | *CORO1C* | *5* | *5* | *TRUE* | *TRUE* |
| *Q9UMZ2* | *SYNRG* | *19* | *19* | *TRUE* | *TRUE* |
| *Q9UNZ2;F2Z2K0* | *NSFL1C* | *6* | *6* | *TRUE* | *TRUE* |
| *Q9UPN7* | *PPP6R1* | *8* | *8* | *TRUE* | *TRUE* |
| *Q9UPQ9;H0Y720* | *TNRC6B* | *35* | *35* | *TRUE* | *TRUE* |
| Q9Y266 | NUDC | 15 | 15 | TRUE | TRUE |
| Q9Y277;E5RJN6;E5 | VDAC3 | 5 | 5 | FALSE | TRUE |
| *Q9Y2G2* | *CARD8* | *3* | *3* | *TRUE* | *TRUE* |
| *Q9Y2Z0* | *SUGT1* | *19* | *19* | *TRUE* | *TRUE* |
| *Q9Y450;B7Z524;H* | *HBS1L* | *12* | *12* | *TRUE* | *TRUE* |
| *Q9Y490* | *TLN1* | *55* | *55* | *TRUE* | *TRUE* |
| *Q9Y4K3* | *TRAF6* | *4* | *4* | *TRUE* | *TRUE* |
| *Q9Y570* | *PPME1* | *2* | *2* | *TRUE* | *TRUE* |
| *Q9Y5A9* | *YTHDF2* | *15* | *15* | *TRUE* | *TRUE* |
| *Q9Y5K6* | *CD2AP* | *14* | *14* | *TRUE* | *TRUE* |
| *Q9Y6G9;E9PHI6* | *DYNC1LI1* | *5* | *5* | *TRUE* | *TRUE* |

| | | | | | |
|---|---|---|---|---|---|
| Q00341;A0A024R4 | HDLBP | 7 | 7 | FALSE | FALSE |
| Q9H4M9;C9JC03;A | EHD1 | 3 | 3 | TRUE | FALSE |
| P25787;A0A024RA | PSMA2 | 4 | 4 | FALSE | FALSE |
| Q16181;A0A024RA | SEPT7 | 7 | 7 | FALSE | FALSE |
| Q3ZCM7;A0A075B | TUBB8 | 6 | 2 | FALSE | FALSE |
| P02766;A0A087W1 | TTR | 2 | 2 | FALSE | FALSE |
| P11940;A0A7I2YQ8 | PABPC1;PABPC3 | 9 | 9 | FALSE | FALSE |
| Q9NP73;A0A096LN | ALG13 | 4 | 4 | TRUE | FALSE |
| Q8TCU4;A0A804HI | ALMS1 | 39 | 39 | FALSE | FALSE |
| Q14511;A0A087WI | NEDD9 | 7 | 7 | FALSE | FALSE |
| O60841;A0A087WI | EIF5B | 9 | 9 | TRUE | FALSE |
| Q9Y6K9;A0A8I5KQ | IKBKG | 4 | 4 | TRUE | FALSE |
| Q01082;A0A087WI | SPTBN1 | 28 | 28 | FALSE | FALSE |
| Q00610;A0A8V8TR | CLTC | 31 | 31 | TRUE | FALSE |
| Q4R9M9;A0A087W | KIF1Bbeta;KIF1B | 8 | 8 | TRUE | FALSE |
| P61962;A0A087WV | DCAF7 | 8 | 8 | TRUE | FALSE |
| A0A087WWU8 | TPM3 | 5 | 5 | FALSE | FALSE |
| Q13148;A0A087WX | TARDBP | 3 | 3 | TRUE | FALSE |
| P18621;A0A087WX | RPL17;RPL17-C1 | 2 | 2 | FALSE | FALSE |
| Q9UPT6;E9PFH7;A | MAPK8IP3 | 4 | 4 | TRUE | FALSE |
| O43399;A0A087WY | TPD52L2 | 4 | 4 | TRUE | FALSE |
| Q15185;A0A087WY | PTGES3 | 3 | 3 | FALSE | FALSE |
| O43314;D6RBU4;A | PPIP5K2;PPIP5K1 | 2 | 2 | FALSE | FALSE |

| | | | | | |
|---|---|---|---|---|---|
| P09668;A0A7I2V67 | CTSH | 2 | 2 | TRUE | FALSE |
| P81877;D6RAC5;A | SSBP2;SSBP3 | 3 | 3 | TRUE | FALSE |
| P53365;A0A087X1 | ARFIP2 | 4 | 4 | TRUE | FALSE |
| P30566;A0A1B0GV | ADSL | 5 | 5 | FALSE | FALSE |
| A0A0A0MQY9;O60 | KLK8 | 5 | 5 | FALSE | FALSE |
| P45880;A0A0A0MF | VDAC2 | 7 | 7 | FALSE | FALSE |
| P15498;A0A0A0MF | VAV1 | 5 | 5 | TRUE | FALSE |
| Q9BUJ2;A0A0A0MF | HNRNPUL1 | 4 | 4 | TRUE | FALSE |
| P41252;A0A804HL | IARS | 13 | 13 | FALSE | FALSE |
| Q63HN8;A0A0A0M | RNF213 | 5 | 5 | FALSE | FALSE |
| P57737;A0A0A6YY | CORO7;CORO7-F | 4 | 4 | TRUE | FALSE |
| P13693;Q5W0H4;A | TPT1 | 3 | 3 | FALSE | FALSE |
| P08865;C9J9K3;A0 | RPSA | 8 | 8 | FALSE | FALSE |
| Q96DA0;A0A0C4D | ZG16B | 4 | 4 | FALSE | FALSE |
| P39656;A0A0C4D( | DDOST | 7 | 7 | FALSE | FALSE |
| Q13813;A0A0D9SF | SPTAN1 | 32 | 32 | FALSE | FALSE |
| Q6NYC8;A0A0G2JH | PPP1R18 | 3 | 3 | FALSE | FALSE |
| A0A0G2JMB2 | IGHA2 | 5 | 2 | FALSE | FALSE |
| P01850;A0A5H1ZR | TRBC1;TRBC2 | 2 | 2 | TRUE | FALSE |
| P08133;A0A0S2Z3 | ANXA6 | 18 | 18 | TRUE | FALSE |
| P04049;H7C155;A | RAF1;ARAF | 4 | 4 | TRUE | FALSE |
| Q9Y484;C9J5L0;C | WDR45 | 2 | 2 | TRUE | FALSE |
| Q7Z401;A0A7I2RA | DENND4A | 3 | 3 | FALSE | FALSE |

| | | | | | |
|---|---|---|---|---|---|
| P26599;A6NLN1;A( | PTBP1 | 8 | 8 | FALSE | FALSE |
| O75955;A0A140T9 | FLOT1 | 6 | 6 | FALSE | FALSE |
| A0A1B0GV12 | GOLGA8N | 2 | 2 | FALSE | FALSE |
| A0A1B0GVL4;P783 | FAM193A | 9 | 9 | TRUE | FALSE |
| A0A1C7CYX9;Q16! | DPYSL2 | 15 | 15 | TRUE | FALSE |
| Q9H3U1;A0A1W2P | UNC45A | 7 | 7 | TRUE | FALSE |
| Q06124;A0A8I5QJ! | PTPN11 | 9 | 9 | FALSE | FALSE |
| Q14353;A0A1W2PI | GAMT | 3 | 3 | TRUE | FALSE |
| Q92841;A0A1X7SB | DDX17 | 11 | 10 | FALSE | FALSE |
| O43175;A0A286YF | PHGDH | 10 | 10 | FALSE | FALSE |
| P31350;A0A7P0SB | RRM2 | 4 | 4 | TRUE | FALSE |
| Q13555;Q5SWX3;/ | CAMK2G | 3 | 3 | TRUE | FALSE |
| Q92574;A0A2R8YE | TSC1 | 2 | 2 | FALSE | FALSE |
| Q15813;A0A2R8Y5 | TBCE | 4 | 4 | TRUE | FALSE |
| P62081;A0A2R8Y6 | RPS7 | 2 | 2 | FALSE | FALSE |
| P06733;A0A2R8Y6 | ENO1 | 10 | 10 | FALSE | FALSE |
| P46777;A0A2R8Y6. | RPL5 | 7 | 7 | FALSE | FALSE |
| Q8N8A2;H7C209;/ | ANKRD44 | 6 | 6 | TRUE | FALSE |
| P62263;A0A2R8Y8 | RPS14 | 3 | 3 | TRUE | FALSE |
| Q9HCK8;A0A2R8Y! | CHD8 | 3 | 3 | FALSE | FALSE |
| P51659;A0A804HK | HSD17B4 | 4 | 4 | TRUE | FALSE |
| P61313;A0A2R8YE | RPL15 | 4 | 4 | FALSE | FALSE |
| Q15437;A0A2R8YF | SEC23B | 2 | 2 | FALSE | FALSE |

| | | | | | |
|---|---|---|---|---|---|
| B1ALK7;A0A8V8TQ | ARHGEF7 | 7 | 5 | FALSE | FALSE |
| P11171;A0A2U3TZ | EPB41 | 8 | 8 | FALSE | FALSE |
| P13987;E9PNW4;A | CD59 | 3 | 3 | FALSE | FALSE |
| P17812;A0A3B3IRI | CTPS1 | 6 | 6 | FALSE | FALSE |
| Q9HCM7;A0A7P0Z | FBRSL1 | 3 | 3 | TRUE | FALSE |
| O14639;F8W8M4;A | ABLIM1 | 4 | 4 | FALSE | FALSE |
| P33991;A0A3B3IU | MCM4 | 11 | 11 | TRUE | FALSE |
| P52948;A0A8V8TR | NUP98 | 5 | 5 | FALSE | FALSE |
| P14866;M0QXS5;A | HNRNPL | 6 | 6 | FALSE | FALSE |
| P50851;A0A494BZ | LRBA | 10 | 10 | FALSE | FALSE |
| Q08188;A0A494C0 | TGM3 | 14 | 14 | FALSE | FALSE |
| P04150;A0A494C0 | NR3C1 | 13 | 13 | TRUE | FALSE |
| P12955;A0A8V8TLI | PEPD | 2 | 2 | TRUE | FALSE |
| P35658;A0A494C1 | NUP214 | 15 | 15 | FALSE | FALSE |
| Q15020;A0A499FI | SART3 | 3 | 3 | FALSE | FALSE |
| A0FGR8;A0A499FI | ESYT2 | 9 | 9 | TRUE | FALSE |
| Q9H3K6;A0A499FJ | BOLA2 | 2 | 2 | FALSE | FALSE |
| Q13303;A0A590UJ | KCNAB2 | 4 | 4 | FALSE | FALSE |
| Q14BN4;A0A994J5 | SLMAP | 4 | 4 | FALSE | FALSE |
| Q13423;A0A590UK | NNT | 4 | 4 | FALSE | FALSE |
| Q8TEU7;A0A5F9ZH | RAPGEF6 | 7 | 7 | FALSE | FALSE |
| P07195;A0A5F9ZH | LDHB | 8 | 7 | TRUE | FALSE |
| Q68CZ6;A0A5F9ZH | HAUS3 | 3 | 3 | TRUE | FALSE |

| | | | | | |
|---|---|---|---|---|---|
| P49643;A0A5F9ZH | PRIM2 | 2 | 2 | TRUE | FALSE |
| P40692;A0A669KA | MLH1 | 5 | 5 | TRUE | FALSE |
| P42224;A0A8V8TN | STAT1 | 6 | 6 | FALSE | FALSE |
| Q93009;A0A669KB | USP7 | 12 | 12 | FALSE | FALSE |
| P12081;B4E1C5;A | HARS | 3 | 3 | FALSE | FALSE |
| P20810;A0A6Q8PF | CAST | 3 | 3 | TRUE | FALSE |
| P43034;A0A6Q8PF | PAFAH1B1 | 4 | 4 | FALSE | FALSE |
| P51149;A0A6Q8PC | RAB7A | 6 | 6 | FALSE | FALSE |
| P56524;A0A7I2SVS | HDAC4 | 5 | 5 | FALSE | FALSE |
| P40763;A0A7I2V39 | STAT3 | 10 | 10 | TRUE | FALSE |
| P38646;A0A7I2V2 | HSPA9 | 9 | 9 | FALSE | FALSE |
| P53634;A0A7I2YQ | CTSC | 6 | 6 | FALSE | FALSE |
| P51991;A0A7I2V2 | HNRNPA3 | 10 | 10 | FALSE | FALSE |
| P12268;A0A7I2V2 | IMPDH2 | 11 | 11 | FALSE | FALSE |
| Q6IBS0;D6RG15;A | TWF2 | 3 | 3 | FALSE | FALSE |
| P10809;A0A7I2V5 | HSPD1 | 14 | 14 | FALSE | FALSE |
| P22626;A0A7I2V3 | HNRNPA2B1;HN | 11 | 11 | TRUE | FALSE |
| P19338;A0A7I2V5 | NCL | 9 | 9 | TRUE | FALSE |
| P09651;A0A7I2V3 | HNRNPA1;HNRN | 10 | 10 | FALSE | FALSE |
| Q01813;A0A7I2V3 | PFKP | 7 | 7 | TRUE | FALSE |
| P62136;A0A7I2V4 | PPP1CA;PPP1CC | 6 | 6 | TRUE | FALSE |
| Q15233;A0A7I2V4 | NONO | 4 | 4 | TRUE | FALSE |
| O60506;A0A7I2V4 | SYNCRIP | 4 | 4 | FALSE | FALSE |

| | | | | | |
|---|---|---|---|---|---|
| O75569;A0A7I2YQ | PRKRA | 5 | 5 | TRUE | FALSE |
| P40429;A0A7I2V5L | RPL13A;RPL13a; | 2 | 2 | TRUE | FALSE |
| Q8TD19;A0A7I2V5 | NEK9 | 5 | 5 | TRUE | FALSE |
| P06748;A0A7I2V5 | NPM1 | 7 | 7 | FALSE | FALSE |
| P07858;A0A7I2YQ | CTSB | 7 | 7 | FALSE | FALSE |
| Q04637;A0A7I2Y1 | EIF4G1 | 15 | 15 | TRUE | FALSE |
| Q13347;A0A7I2YQ | EIF3I | 8 | 8 | FALSE | FALSE |
| P60866;A0A7P0S5 | RPS20 | 3 | 3 | FALSE | FALSE |
| P27797;A0A7P0T8 | CALR | 7 | 7 | FALSE | FALSE |
| Q16531;A0A7P0TB | DDB1 | 9 | 9 | FALSE | FALSE |
| Q9Y5X1;A0A7P0T8 | SNX9 | 8 | 8 | TRUE | FALSE |
| P07237;I3L3P5;A0 | P4HB | 6 | 6 | FALSE | FALSE |
| P08195;A0A7P0Z4 | SLC3A2 | 9 | 9 | TRUE | FALSE |
| O95573;A0A7P0TA | ACSL3 | 11 | 11 | TRUE | FALSE |
| Q14203;E7EX90;A | DCTN1;DKFZp68 | 6 | 6 | FALSE | FALSE |
| O95630;F5H5B9;A | STAMBP | 5 | 5 | TRUE | FALSE |
| Q9BTW9;A0A804H | TBCD | 2 | 2 | FALSE | FALSE |
| P07954;A0A804HI | FH | 2 | 2 | TRUE | FALSE |
| P42704;A0A804HI | LRPPRC | 13 | 13 | FALSE | FALSE |
| P12814;A0A804HK | ACTN1 | 14 | 14 | FALSE | FALSE |
| P50402;A0A804HI | EMD | 3 | 3 | TRUE | FALSE |
| Q9GZZ9;A0A804HJ | UBA5 | 4 | 4 | TRUE | FALSE |
| A0A804HKD7;Q5R | RABGAP1L | 14 | 14 | TRUE | FALSE |

| | | | | | |
|---|---|---|---|---|---|
| P27816;E7EVA0;A( | MAP4 | 24 | 24 | TRUE | FALSE |
| Q08378;A0A8I5KW | GOLGA3 | 7 | 7 | FALSE | FALSE |
| P00387;A0A8I5QK) | CYB5R3 | 6 | 6 | FALSE | FALSE |
| P49848;C9JTY6;A0 | TAF6 | 8 | 8 | FALSE | FALSE |
| P30101;A0A8I5KW | PDIA3 | 9 | 9 | FALSE | FALSE |
| O60307;A0A8V8TM | MAST3 | 5 | 5 | FALSE | FALSE |
| Q15370;A0A8I5KU: | TCEB2 | 3 | 3 | TRUE | FALSE |
| P35579;A0A8I5KW | MYH9 | 47 | 47 | FALSE | FALSE |
| Q5T1M5;A0A8I5KX | FKBP15 | 9 | 9 | FALSE | FALSE |
| Q06787;A0A8I5KZ: | FMR1 | 9 | 7 | TRUE | FALSE |
| P35580;A0A8I5KZ3 | MYH10 | 9 | 2 | FALSE | FALSE |
| Q14789;A0A8Q3W | GOLGB1 | 6 | 6 | FALSE | FALSE |
| P00813;A0A8Q3SI/ | ADA | 9 | 9 | FALSE | FALSE |
| P11413;A0A8Q3SJI | G6PD | 7 | 7 | TRUE | FALSE |
| P00491;A0A8V8TL: | PNP | 4 | 4 | FALSE | FALSE |
| P11586;A0A8V8TL! | MTHFD1 | 8 | 8 | TRUE | FALSE |
| Q9Y244;A0A8V8TM | POMP | 2 | 2 | FALSE | FALSE |
| P02786;A0A8V8TLI | TFRC | 11 | 11 | TRUE | FALSE |
| Q86UX7;A0A8V8TN | FERMT3 | 14 | 14 | TRUE | FALSE |
| P30085;Q5T0D2;A( | CMPK1 | 2 | 2 | FALSE | FALSE |
| P51114;A0A8V8TN | FXR1 | 14 | 14 | TRUE | FALSE |
| Q86TI0;A0A8V8TN! | TBC1D1 | 4 | 4 | FALSE | FALSE |
| O75083;A0A8V8TN | WDR1 | 12 | 12 | FALSE | FALSE |

| | | | | | |
|---|---|---|---|---|---|
| Q96B97;Q5JPT2;A( | SH3KBP1 | 10 | 10 | TRUE | FALSE |
| P14618;A0A8V8TN | PKM | 20 | 20 | FALSE | FALSE |
| P61204;P84077;A( | ARF3;ARF1;ARF5 | 4 | 4 | FALSE | FALSE |
| P20963;A0A8V8TP | CD247;TCRZ | 5 | 5 | FALSE | FALSE |
| Q99567;J3KMX1;A( | NUP88 | 8 | 8 | FALSE | FALSE |
| P30153;A0A994J3I | PPP2R1A | 8 | 8 | FALSE | FALSE |
| Q96AE4;E9PEB5;A( | FUBP1 | 5 | 4 | TRUE | FALSE |
| P62873;F6X3N5;A( | GNB1;GNB2 | 6 | 6 | FALSE | FALSE |
| Q14103;H0YA96;D | HNRNPD | 3 | 3 | FALSE | FALSE |
| P31629;A0A994J4 | HIVEP2 | 4 | 4 | FALSE | FALSE |
| Q6P2Q9;I3L3Z8;A0 | PRPF8 | 7 | 7 | FALSE | FALSE |
| P61978;A0A994J6 | HNRNPK | 11 | 11 | TRUE | FALSE |
| Q8IW35;A0A994J4 | CEP97 | 7 | 7 | FALSE | FALSE |
| E9PK52;E9PHY5;A( | EPB41L2 | 4 | 4 | TRUE | FALSE |
| P04844;A0A994J5J | RPN2 | 4 | 4 | FALSE | FALSE |
| P46459;A0A994J5I | NSF | 3 | 3 | FALSE | FALSE |
| Q92614;A0A994J7 | MYO18A | 13 | 13 | FALSE | FALSE |
| Q8TAT6;A0A994J7I | NPLOC4 | 7 | 7 | FALSE | FALSE |
| A0A994J7P9;P511 | FXR2 | 12 | 10 | TRUE | FALSE |
| Q9Y217;A0A9L9PX | MTMR6 | 4 | 4 | FALSE | FALSE |
| P13861;A0A9L9PY | PRKAR2A | 6 | 6 | TRUE | FALSE |
| A0AA34QVE6;Q14 | UBAP2L | 19 | 19 | FALSE | FALSE |
| P51858;A0AA34QV | HDGF | 3 | 3 | FALSE | FALSE |

| | | | | | |
|---|---|---|---|---|---|
| A5YKK6 | CNOT1 | 12 | 12 | FALSE | FALSE |
| P51610;A6NEM2 | HCFC1 | 6 | 6 | FALSE | FALSE |
| Q9UKK9;A6NFX8;A | NUDT5 | 14 | 14 | TRUE | FALSE |
| Q9H910;A6NGP5;H | HN1L | 4 | 4 | TRUE | FALSE |
| P46020;A6NIT2;A6 | PHKA1 | 4 | 4 | FALSE | FALSE |
| P54578;A6NJA2 | USP14 | 6 | 6 | FALSE | FALSE |
| Q9H4A4;A6NKB8;A | RNPEP | 4 | 4 | TRUE | FALSE |
| Q99829;F2Z2V0;A6 | CPNE1 | 2 | 2 | FALSE | FALSE |
| P55036;Q5VWC4;A | PSMD4 | 3 | 3 | TRUE | FALSE |
| P18124;A8MUD9 | RPL7 | 5 | 5 | FALSE | FALSE |
| P55039;A8MZF9;J3 | DRG2 | 2 | 2 | TRUE | FALSE |
| P48444;B0YIW6 | ARCN1 | 5 | 5 | TRUE | FALSE |
| P12956;B1AHC9 | XRCC6 | 7 | 7 | TRUE | FALSE |
| P47756;B1AK88;B1 | CAPZB | 9 | 9 | FALSE | FALSE |
| Q14141;B1AMS2 | SEPT6 | 4 | 4 | FALSE | FALSE |
| Q9Y2S7;B4DEM9 | POLDIP2 | 3 | 3 | TRUE | FALSE |
| P49368;B4DUR8 | CCT3 | 18 | 18 | FALSE | FALSE |
| P0CG48;P0CG47;C | UBC;UBB;RPS27A | 6 | 6 | FALSE | FALSE |
| P50542;B4E0T2;J3 | PEX5 | 5 | 5 | FALSE | FALSE |
| Q15019;B5MCX3;H | SEPT2 | 3 | 3 | TRUE | FALSE |
| Q99613;B5ME19 | EIF3C;EIF3CL | 10 | 10 | FALSE | FALSE |
| Q9UI42;B7Z5J4 | CPA4 | 10 | 10 | FALSE | FALSE |
| P60660;F8W1R7;J3 | MYL6 | 7 | 7 | FALSE | FALSE |

| | | | | | |
|---|---|---|---|---|---|
| Q6JBY9;B7ZKW8;F( | RCSD1 | 6 | 6 | TRUE | FALSE |
| Q9Y618;C9JE98;C9 | NCOR2 | 4 | 4 | FALSE | FALSE |
| Q99733;E9PNW0;E | NAP1L4 | 5 | 5 | TRUE | FALSE |
| P05090;C9JF17 | APOD | 5 | 5 | FALSE | FALSE |
| Q8N3P4;C9JJN9 | VPS8 | 7 | 7 | FALSE | FALSE |
| P52434;C9JLU1 | POLR2H | 4 | 4 | FALSE | FALSE |
| Q99729;D6R9P3;D | HNRNPAB | 4 | 4 | FALSE | FALSE |
| P32969;D6RAN4;A | RPL9 | 5 | 5 | FALSE | FALSE |
| P06730;D6RBW1;D | EIF4E | 2 | 2 | TRUE | FALSE |
| Q9NVA2;D6RDU5;I | SEPT11 | 3 | 2 | TRUE | FALSE |
| Q96CN7;D6RGE2 | ISOC1 | 2 | 2 | TRUE | FALSE |
| P63208;E5RJR5;E5 | SKP1 | 2 | 2 | FALSE | FALSE |
| Q7L5N1;E7EM64 | COPS6 | 3 | 3 | TRUE | FALSE |
| P0DP25;P0DP24;P | CALM2;CALM1 | 3 | 3 | FALSE | FALSE |
| Q14254;J3QLD9;E7 | FLOT2 | 10 | 10 | FALSE | FALSE |
| Q9H0E2;E9PP67;E | TOLLIP | 3 | 3 | FALSE | FALSE |
| Q9Y520;E7EPN9 | PRRC2C | 19 | 19 | FALSE | FALSE |
| P02788;E7EQB2;E7 | LTF | 16 | 16 | FALSE | FALSE |
| Q9NRY5;E7ESJ7 | FAM114A2 | 9 | 9 | FALSE | FALSE |
| Q07866;G3V2E7;F8 | KLC1 | 7 | 7 | TRUE | FALSE |
| P23588;E7EX17;F8 | EIF4B | 7 | 7 | TRUE | FALSE |
| P78559;E9PGC8 | MAP1A | 22 | 22 | TRUE | FALSE |
| Q15003;E9PHA2;C | NCAPH | 7 | 7 | TRUE | FALSE |

| | | | | | |
|---|---|---|---|---|---|
| P23528;G3V1A4;E9 | CFL1 | 11 | 11 | TRUE | FALSE |
| Q99873;E9PKG1;E9 | PRMT1 | 7 | 7 | TRUE | FALSE |
| P06239;E9PKQ8;E9 | LCK | 7 | 7 | FALSE | FALSE |
| P62917;E9PKZ0;G3 | RPL8 | 3 | 3 | TRUE | FALSE |
| Q8N9N8;E9PQD0;I | EIF1AD | 2 | 2 | TRUE | FALSE |
| P78406;E9PQ57;E9 | RAE1 | 5 | 5 | FALSE | FALSE |
| P78371;F5GWF6;F | CCT2 | 12 | 12 | TRUE | FALSE |
| O00233;F5H169;F5 | PSMD9 | 2 | 2 | TRUE | FALSE |
| Q15436;F5H365 | SEC23A | 2 | 2 | FALSE | FALSE |
| P27708;F8VPD4 | CAD | 16 | 16 | FALSE | FALSE |
| P11802;F8VXD2;F8 | CDK4 | 3 | 2 | FALSE | FALSE |
| F8VVM2;Q00325;F | SLC25A3 | 3 | 3 | FALSE | FALSE |
| Q14161;F8W822;F | GIT2 | 4 | 3 | FALSE | FALSE |
| Q9UBQ0;F8VXU5 | VPS29 | 5 | 5 | TRUE | FALSE |
| Q9NR82;F8WEA4 | KCNQ5 | 5 | 5 | FALSE | FALSE |
| Q14444;G3V153;E9 | CAPRIN1 | 4 | 4 | TRUE | FALSE |
| Q9NY33;G3V1D3;G | DPP3 | 4 | 4 | FALSE | FALSE |
| Q07020;G3V203;H | RPL18 | 3 | 3 | FALSE | FALSE |
| P49757;G3V3Z8 | NUMB | 4 | 4 | TRUE | FALSE |
| Q9H9C1;G3V4K3 | VIPAS39 | 6 | 6 | TRUE | FALSE |
| Q9H6D7;G3V541;G | HAUS4 | 4 | 4 | TRUE | FALSE |
| P60900;G3V5Z7;G3 | PSMA6 | 4 | 4 | TRUE | FALSE |
| Q15653;I3L4X3;G5 | NFKBIB | 6 | 6 | TRUE | FALSE |

| | | | | | |
|---|---|---|---|---|---|
| P39023;G5E9G0;B | RPL3 | 9 | 9 | FALSE | FALSE |
| P31943;G8JLB6;A0 | HNRNPH1 | 9 | 9 | FALSE | FALSE |
| Q9BQA1;H0Y711 | WDR77 | 3 | 3 | TRUE | FALSE |
| Q12849;H0Y8R1;H | GRSF1 | 5 | 5 | FALSE | FALSE |
| Q7LG56;H0YAV1 | RRM2B | 3 | 2 | TRUE | FALSE |
| Q96Q05;H0YBR0 | TRAPPC9 | 3 | 3 | FALSE | FALSE |
| Q3LXA3;H0YCY6;E | DAK;TKFC | 4 | 4 | FALSE | FALSE |
| H0YNW5;P33316;H | DUT | 3 | 3 | TRUE | FALSE |
| P62491;Q15907;H | RAB11A;RAB11B | 5 | 5 | FALSE | FALSE |
| P04075;H3BQN4;J | ALDOA | 11 | 11 | FALSE | FALSE |
| Q9BWH6;H3BRE8; | RPAP1 | 4 | 4 | FALSE | FALSE |
| Q8N1F7;H3BVG0 | NUP93 | 5 | 5 | FALSE | FALSE |
| O96019;H7C5S0 | ACTL6A | 3 | 3 | TRUE | FALSE |
| Q6Y7W6;I1E4Y6;A( | GIGYF2 | 12 | 12 | FALSE | FALSE |
| Q99447;I3L1R7;I3L | PCYT2 | 4 | 4 | TRUE | FALSE |
| O14641;I3L2W9;I3 | DVL2 | 4 | 4 | TRUE | FALSE |
| P63241;I3L397;I3L | EIF5A;EIF5A2;EIF | 4 | 4 | FALSE | FALSE |
| P52655;J3KNC0 | GTF2A1 | 2 | 2 | TRUE | FALSE |
| Q8WYJ6;J3KNL2 | SEPT1 | 11 | 11 | TRUE | FALSE |
| P62826;J3KQE5;F5 | RAN | 12 | 12 | TRUE | FALSE |
| P17844;J3KTA4;A0 | DDX5 | 10 | 6 | FALSE | FALSE |
| Q9UHR5;J3QQJ0;J | SAP30BP | 2 | 2 | TRUE | FALSE |
| P62195;J3QSA9;J3 | PSMC5 | 5 | 5 | FALSE | FALSE |

| | | | | | |
|---|---|---|---|---|---|
| O43684;J3QT28;J3 | BUB3 | 5 | 5 | TRUE | FALSE |
| O96008;K7EJ57 | TOMM40 | 3 | 3 | FALSE | FALSE |
| Q96EP5;K7EQ02;K | DAZAP1 | 2 | 2 | FALSE | FALSE |
| O75821;K7ER90;K7 | EIF3G | 3 | 3 | TRUE | FALSE |
| Q96EE3;K7ELV2;K7 | SEH1L | 4 | 4 | FALSE | FALSE |
| Q99426;K7EP07;K7 | TBCB | 5 | 5 | TRUE | FALSE |
| M0QY01 | EPS15L1 | 2 | 2 | FALSE | FALSE |
| Q92945;M0R0C6;M | KHSRP | 9 | 9 | FALSE | FALSE |
| Q02543;M0R0P7;M | RPL18A | 3 | 3 | FALSE | FALSE |
| O95336;M0R261 | PGLS | 4 | 4 | TRUE | FALSE |
| Q9NZ01;M0R3C3;M | TECR | 2 | 2 | FALSE | FALSE |
| O00151 | PDLIM1 | 14 | 14 | TRUE | FALSE |
| O00154;K7EKP8 | ACOT7 | 7 | 7 | FALSE | FALSE |
| O00221;Q7LC14 | NFKBIE | 11 | 11 | TRUE | FALSE |
| O00231;J3QRY4 | PSMD11 | 8 | 8 | TRUE | FALSE |
| O00273 | DFFA | 3 | 3 | FALSE | FALSE |
| O00299 | CLIC1 | 8 | 8 | FALSE | FALSE |
| O00505 | KPNA3 | 4 | 4 | FALSE | FALSE |
| O00763 | ACACB | 64 | 54 | FALSE | FALSE |
| O14745;J3QRP6 | SLC9A3R1 | 8 | 8 | TRUE | FALSE |
| O14818;Q8TAA3;A | PSMA7;PSMA8 | 4 | 4 | TRUE | FALSE |
| O14929 | HAT1 | 4 | 4 | FALSE | FALSE |
| O14976;D6RAW3 | GAK | 2 | 2 | FALSE | FALSE |

| | | | | | |
|---|---|---|---|---|---|
| O15042;E7ET15 | U2SURP | 4 | 4 | TRUE | FALSE |
| O15067 | PFAS | 13 | 13 | FALSE | FALSE |
| O15143;C9JEY1;C9 | ARPC1B | 7 | 7 | FALSE | FALSE |
| O15173;U3KQM0 | PGRMC2 | 4 | 4 | TRUE | FALSE |
| O15371 | EIF3D | 7 | 7 | TRUE | FALSE |
| O15397 | IPO8 | 5 | 5 | TRUE | FALSE |
| O15511;B1ALC0 | ARPC5 | 2 | 2 | TRUE | FALSE |
| O43143 | DHX15 | 6 | 6 | FALSE | FALSE |
| O43164 | PJA2 | 4 | 4 | TRUE | FALSE |
| O43290 | SART1 | 3 | 3 | FALSE | FALSE |
| O43303 | CCP110 | 13 | 13 | FALSE | FALSE |
| O43396;K7ER96 | TXNL1 | 9 | 9 | FALSE | FALSE |
| O43809;H3BV41;H | NUDT21 | 4 | 4 | FALSE | FALSE |
| O43815 | STRN | 5 | 5 | FALSE | FALSE |
| O60271 | SPAG9 | 7 | 7 | FALSE | FALSE |
| O60711;B7Z5P7 | LPXN | 13 | 13 | TRUE | FALSE |
| O60749 | SNX2 | 6 | 5 | FALSE | FALSE |
| O60880 | SH2D1A | 3 | 3 | FALSE | FALSE |
| O60888;C9IZG4 | CUTA | 2 | 2 | FALSE | FALSE |
| O75131;A0A087W | CPNE3 | 4 | 4 | FALSE | FALSE |
| O75170 | PPP6R2 | 10 | 10 | FALSE | FALSE |
| O75223;M0QZK8;E | GGCT | 7 | 7 | FALSE | FALSE |
| O75351 | VPS4B | 4 | 4 | TRUE | FALSE |

| | | | | | |
|---|---|---|---|---|---|
| O75369;A0A804Hk | FLNB | 23 | 19 | FALSE | FALSE |
| O75533 | SF3B1 | 15 | 15 | FALSE | FALSE |
| O75635 | SERPINB7 | 5 | 5 | FALSE | FALSE |
| O75663 | TIPRL | 11 | 11 | TRUE | FALSE |
| O75792;A0A2R8Y6 | RNASEH2A | 2 | 2 | FALSE | FALSE |
| O75995 | SASH3 | 3 | 3 | TRUE | FALSE |
| O95274 | LYPD3 | 2 | 2 | FALSE | FALSE |
| O95347 | SMC2 | 3 | 3 | FALSE | FALSE |
| O95372;Q5QPQ0 | LYPLA2 | 3 | 3 | TRUE | FALSE |
| O95373 | IPO7 | 7 | 7 | TRUE | FALSE |
| O95405 | ZFYVE9 | 3 | 3 | FALSE | FALSE |
| O95433;H0YJG7;G | AHSA1 | 8 | 8 | TRUE | FALSE |
| O95782 | AP2A1 | 11 | 11 | FALSE | FALSE |
| O95819 | MAP4K4 | 9 | 9 | FALSE | FALSE |
| P00338 | LDHA | 15 | 15 | FALSE | FALSE |
| P00441;H7BYH4 | SOD1 | 6 | 6 | FALSE | FALSE |
| P01037;P01036 | CST1;CST4 | 3 | 3 | FALSE | FALSE |
| P01040;C9J0E4 | CSTA | 13 | 13 | FALSE | FALSE |
| P01833 | PIGR | 11 | 11 | FALSE | FALSE |
| P01857;A0A0A0MS | IGHG1 | 7 | 7 | FALSE | FALSE |
| P01876 | IGHA1 | 6 | 6 | FALSE | FALSE |
| P02787 | TF | 8 | 8 | FALSE | FALSE |
| P04040 | CAT | 17 | 17 | FALSE | FALSE |

| | | | | | |
|---|---|---|---|---|---|
| P04080;A0A1W2PS | CSTB | 4 | 4 | FALSE | FALSE |
| P04406;E7EUT5 | GAPDH | 15 | 15 | FALSE | FALSE |
| P04818 | TYMS | 5 | 5 | TRUE | FALSE |
| P04843;B7Z4L4 | RPN1 | 10 | 10 | FALSE | FALSE |
| P05023 | ATP1A1 | 6 | 6 | FALSE | FALSE |
| P05089;A0A5F9ZH | ARG1 | 9 | 9 | FALSE | FALSE |
| P05109 | S100A8 | 5 | 5 | FALSE | FALSE |
| P05165;A0A1B0GU | PCCA | 25 | 25 | FALSE | FALSE |
| P05198;H0YJS4;G3 | EIF2S1 | 7 | 7 | FALSE | FALSE |
| P05387;H0YDD8 | RPLP2 | 2 | 2 | TRUE | FALSE |
| P06576;H0YH81;F8 | ATP5B | 11 | 11 | FALSE | FALSE |
| P06702 | S100A9 | 7 | 7 | FALSE | FALSE |
| P07355;H0YMW4;H | ANXA2;ANXA2P2 | 17 | 17 | FALSE | FALSE |
| P07384 | CAPN1 | 11 | 11 | FALSE | FALSE |
| P07686;Q5URX0 | HEXB | 3 | 3 | FALSE | FALSE |
| P07737;K7EJ44;I3L | PFN1 | 8 | 8 | FALSE | FALSE |
| P07814;V9GYZ6 | EPRS | 16 | 16 | TRUE | FALSE |
| P07900 | HSP90AA1 | 29 | 29 | TRUE | FALSE |
| P08238 | HSP90AB1 | 28 | 16 | FALSE | FALSE |
| P08670;B0YJC4;B0 | VIM | 18 | 18 | FALSE | FALSE |
| P09211;A8MX94 | GSTP1 | 8 | 8 | FALSE | FALSE |
| P09960 | LTA4H | 6 | 6 | FALSE | FALSE |
| P0DUB6;P0DTE8;P | AMY2B;AMY2A | 14 | 14 | FALSE | FALSE |

| | | | | | |
|---|---|---|---|---|---|
| P10412;P16402 | HIST1H1E;HIST1H | 4 | 4 | FALSE | FALSE |
| P10599 | TXN | 4 | 4 | FALSE | FALSE |
| P11274 | BCR | 3 | 3 | TRUE | FALSE |
| P11279 | LAMP1 | 2 | 2 | FALSE | FALSE |
| P11717;A0A7I2V38 | IGF2R | 71 | 71 | TRUE | FALSE |
| P11908;P60891;B1 | PRPS2;PRPS1;PR | 4 | 4 | TRUE | FALSE |
| P12004 | PCNA | 3 | 3 | FALSE | FALSE |
| P12236;P12235;V9 | SLC25A6;SLC25A | 7 | 7 | FALSE | FALSE |
| P12273 | PIP | 6 | 6 | FALSE | FALSE |
| P13010 | XRCC5 | 10 | 10 | TRUE | FALSE |
| P13489 | RNH1 | 9 | 9 | FALSE | FALSE |
| P13612;E7EP60 | ITGA4 | 3 | 3 | FALSE | FALSE |
| P13796 | LCP1 | 28 | 28 | TRUE | FALSE |
| P14174 | MIF | 2 | 2 | FALSE | FALSE |
| P14324;A0A087X1 | FDPS | 4 | 4 | FALSE | FALSE |
| P14868 | DARS | 17 | 17 | FALSE | FALSE |
| P14923 | JUP | 20 | 20 | FALSE | FALSE |
| P15170;H3BR35 | GSPT1 | 7 | 7 | TRUE | FALSE |
| P15880;H0YEN5;E | RPS2 | 7 | 7 | FALSE | FALSE |
| P15924 | DSP | 74 | 74 | FALSE | FALSE |
| P15927;Q5TEJ0 | RPA2 | 3 | 3 | FALSE | FALSE |
| P16401 | HIST1H1B | 2 | 2 | FALSE | FALSE |
| P16615;H7C5W9 | ATP2A2 | 7 | 7 | FALSE | FALSE |

| | | | | | |
|---|---|---|---|---|---|
| P17900 | GM2A | 4 | 4 | FALSE | FALSE |
| P18206 | VCL | 9 | 9 | FALSE | FALSE |
| P18510 | IL1RN | 2 | 2 | FALSE | FALSE |
| P18669 | PGAM1 | 7 | 7 | FALSE | FALSE |
| P19474 | TRIM21 | 3 | 3 | TRUE | FALSE |
| P19623;K7ESL0 | SRM | 6 | 6 | FALSE | FALSE |
| P21291;E9PND2;E | CSRP1 | 3 | 3 | TRUE | FALSE |
| P21399 | ACO1 | 20 | 20 | TRUE | FALSE |
| P21580 | TNFAIP3 | 2 | 2 | TRUE | FALSE |
| P21796;C9JI87 | VDAC1 | 9 | 9 | FALSE | FALSE |
| P22059 | OSBP | 9 | 9 | FALSE | FALSE |
| P22061;F6S8N6;A | PCMT1 | 5 | 5 | TRUE | FALSE |
| P22102 | GART | 10 | 10 | TRUE | FALSE |
| P22314;Q5JRR6 | UBA1 | 12 | 12 | FALSE | FALSE |
| P22735 | TGM1 | 6 | 6 | FALSE | FALSE |
| P22830;A0A499FJI | FECH | 3 | 3 | FALSE | FALSE |
| P23396;E9PPU1;H | RPS3 | 8 | 8 | FALSE | FALSE |
| P23743;G3V4E1 | DGKA | 4 | 4 | FALSE | FALSE |
| P24534 | EEF1B2 | 5 | 4 | TRUE | FALSE |
| P25098;E9PRV7;P | ADRBK1;ADRBK2 | 4 | 4 | TRUE | FALSE |
| P25205;J3KQ69;A | MCM3 | 9 | 9 | TRUE | FALSE |
| P25311;C9JEV0 | AZGP1 | 11 | 11 | FALSE | FALSE |
| P25398 | RPS12 | 5 | 5 | FALSE | FALSE |

| | | | | | |
|---|---|---|---|---|---|
| P25705;K7ERX7 | ATP5A1 | 10 | 10 | FALSE | FALSE |
| P25788;G3V4X5 | PSMA3 | 4 | 4 | FALSE | FALSE |
| P26038;A0A8V8TKI | MSN | 22 | 22 | FALSE | FALSE |
| P26368 | U2AF2 | 8 | 8 | TRUE | FALSE |
| P26373;J3QSB4 | RPL13 | 4 | 4 | FALSE | FALSE |
| P26641 | EEF1G | 8 | 8 | FALSE | FALSE |
| P27348;E9PG15 | YWHAQ | 9 | 7 | FALSE | FALSE |
| P27482 | CALML3 | 5 | 5 | FALSE | FALSE |
| P27694 | RPA1 | 8 | 8 | FALSE | FALSE |
| P28066 | PSMA5 | 5 | 5 | FALSE | FALSE |
| P28072;I3L3X7;A0/ | PSMB6 | 2 | 2 | FALSE | FALSE |
| P28482 | MAPK1 | 7 | 7 | FALSE | FALSE |
| P29401 | TKT | 5 | 5 | FALSE | FALSE |
| P29508;P48594;H( | SERPINB3;SERPII | 16 | 16 | FALSE | FALSE |
| P30044 | PRDX5 | 2 | 2 | FALSE | FALSE |
| P30048 | PRDX3 | 6 | 6 | FALSE | FALSE |
| P30050 | RPL12 | 3 | 3 | TRUE | FALSE |
| P31025 | LCN1 | 6 | 6 | FALSE | FALSE |
| P31146;H3BRY3 | CORO1A | 10 | 10 | FALSE | FALSE |
| P31151 | S100A7 | 9 | 9 | FALSE | FALSE |
| P31153 | MAT2A | 3 | 3 | TRUE | FALSE |
| P31689 | DNAJA1 | 7 | 7 | FALSE | FALSE |
| P31930 | UQCRC1 | 5 | 5 | FALSE | FALSE |

| | | | | | |
|---|---|---|---|---|---|
| P31939 | ATIC | 11 | 11 | FALSE | FALSE |
| P31944 | CASP14 | 18 | 18 | FALSE | FALSE |
| P32119 | PRDX2 | 10 | 9 | FALSE | FALSE |
| P32321;D6RBJ9 | DCTD | 4 | 4 | TRUE | FALSE |
| P33993;C9J8M6;A( | MCM7 | 8 | 8 | TRUE | FALSE |
| P34932 | HSPA4 | 16 | 16 | TRUE | FALSE |
| P35268;K7EJT5;K7I | RPL22 | 3 | 3 | FALSE | FALSE |
| P36578;H3BU31;H | RPL4 | 9 | 9 | FALSE | FALSE |
| P37198;M0QXN5 | NUP62 | 7 | 7 | FALSE | FALSE |
| P37235;E9PC71 | HPCAL1 | 5 | 5 | TRUE | FALSE |
| P40121;E7ENU9;A( | CAPG | 7 | 7 | FALSE | FALSE |
| P40227 | CCT6A | 12 | 12 | TRUE | FALSE |
| P40926 | MDH2 | 3 | 3 | FALSE | FALSE |
| P41091;Q2VIR3 | EIF2S3;EIF2S3L | 8 | 8 | TRUE | FALSE |
| P41240 | CSK | 5 | 5 | TRUE | FALSE |
| P42166 | TMPO | 11 | 11 | FALSE | FALSE |
| P42285 | SKIV2L2 | 3 | 3 | FALSE | FALSE |
| P42574 | CASP3 | 9 | 9 | TRUE | FALSE |
| P43403 | ZAP70 | 9 | 9 | TRUE | FALSE |
| P43487;C9JXG8;C! | RANBP1 | 5 | 5 | TRUE | FALSE |
| P46060 | RANGAP1 | 10 | 10 | TRUE | FALSE |
| P46109 | CRKL | 9 | 9 | TRUE | FALSE |
| P47755;F8W9N7;C | CAPZA2 | 7 | 5 | FALSE | FALSE |

| | | | | | |
|---|---|---|---|---|---|
| P48634 | PRRC2A | 10 | 10 | FALSE | FALSE |
| P48735;H0YL11 | IDH2 | 7 | 7 | FALSE | FALSE |
| P49321;Q5T624;H( | NASP | 5 | 5 | FALSE | FALSE |
| P49327;A0A0U1RQ | FASN | 42 | 42 | TRUE | FALSE |
| P49411 | TUFM | 7 | 7 | FALSE | FALSE |
| P49458;Q6P2S0;Q | SRP9;DKFZp564M | 2 | 2 | FALSE | FALSE |
| P49591;Q5T5C7 | SARS | 19 | 19 | TRUE | FALSE |
| P49750;F8VU51 | YLPM1 | 5 | 5 | FALSE | FALSE |
| P49773 | HINT1 | 5 | 5 | FALSE | FALSE |
| P49792;A0A8V8TLI | RANBP2 | 42 | 42 | TRUE | FALSE |
| P49862;M0QYU8 | KLK7 | 4 | 4 | FALSE | FALSE |
| P50395 | GDI2 | 14 | 14 | FALSE | FALSE |
| P50502;H7C3I1;Q8 | ST13;ST13P5;ST1 | 7 | 7 | TRUE | FALSE |
| P50570;K7EMQ3 | DNM2 | 3 | 3 | FALSE | FALSE |
| P51452;K7ES89;K7 | DUSP3 | 4 | 4 | TRUE | FALSE |
| P51571;A6NLM8 | SSR4 | 3 | 3 | FALSE | FALSE |
| P52272 | HNRNPM | 10 | 10 | TRUE | FALSE |
| P52306 | RAP1GDS1 | 13 | 13 | FALSE | FALSE |
| P52566;F5H6Q0;F5 | ARHGDIB | 9 | 9 | FALSE | FALSE |
| P52594;B8ZZY2;C9 | AGFG1 | 7 | 7 | TRUE | FALSE |
| P52597 | HNRNPF | 4 | 2 | FALSE | FALSE |
| P52907 | CAPZA1 | 8 | 8 | FALSE | FALSE |
| P53004;C9J1E1 | BLVRA | 3 | 3 | TRUE | FALSE |

| | | | | | | |
|---|---|---|---|---|---|---|
| P53367 | ARFIP1 | 8 | | 8 | TRUE | FALSE |
| P53396 | ACLY | 10 | | 10 | FALSE | FALSE |
| P53618;E9PP73 | COPB1 | 16 | | 16 | TRUE | FALSE |
| P53990;F5GXM3;H | IST1 | 7 | | 7 | TRUE | FALSE |
| P53999 | SUB1 | 2 | | 2 | TRUE | FALSE |
| P54577;A0A6Q8PF | YARS | 11 | | 11 | TRUE | FALSE |
| P54652 | HSPA2 | 9 | | 3 | FALSE | FALSE |
| P55000 | SLURP1 | 2 | | 2 | FALSE | FALSE |
| P55735;A8MXL6 | SEC13 | 6 | | 6 | TRUE | FALSE |
| P55884;C9JZG1 | EIF3B | 6 | | 6 | FALSE | FALSE |
| P56537;B7ZBH1 | EIF6 | 5 | | 5 | FALSE | FALSE |
| P58546 | MTPN | 2 | | 2 | FALSE | FALSE |
| P60174;U3KPZ0 | TPI1 | 9 | | 9 | FALSE | FALSE |
| P60842;J3QS69;J3 | EIF4A1 | 10 | | 10 | FALSE | FALSE |
| P60953;A0A8Q3WI | CDC42 | 7 | | 7 | FALSE | FALSE |
| P60981;F6RFD5 | DSTN | 4 | | 4 | FALSE | FALSE |
| P61006 | RAB8A | 3 | | 2 | FALSE | FALSE |
| P61009 | SPCS3 | 3 | | 3 | FALSE | FALSE |
| P61011;G3V480;G | SRP54 | 17 | | 17 | TRUE | FALSE |
| P61019;E9PKL7;Q8 | RAB2A;RAB2B | 5 | | 5 | FALSE | FALSE |
| P61026 | RAB10 | 3 | | 2 | FALSE | FALSE |
| P61081 | UBE2M | 6 | | 6 | TRUE | FALSE |
| P61088;Q5JXB2;F8 | UBE2N;UBE2NL | 3 | | 3 | FALSE | FALSE |

| | | | | | |
|---|---|---|---|---|---|
| P61106;X6RFL8;A0 | RAB14 | 5 | 5 | FALSE | FALSE |
| P61158;B4DXW1 | ACTR3 | 15 | 15 | TRUE | FALSE |
| P61221;D6R9I9 | ABCE1 | 13 | 13 | TRUE | FALSE |
| P61326;Q96A72;F5 | MAGOH;MAGOHB | 5 | 5 | TRUE | FALSE |
| P61586;A0A7I2YQ\ | RHOA;RHOC | 4 | 4 | FALSE | FALSE |
| P61604;B8ZZ54;B8 | HSPE1 | 4 | 3 | FALSE | FALSE |
| P61970;H3BRV9;A( | NUTF2 | 6 | 6 | FALSE | FALSE |
| P62241;Q5JR95 | RPS8 | 5 | 5 | TRUE | FALSE |
| P62258;K7EM20 | YWHAE | 10 | 10 | TRUE | FALSE |
| P62314 | SNRPD1 | 3 | 3 | TRUE | FALSE |
| P62318;H3BT13 | SNRPD3 | 2 | 2 | FALSE | FALSE |
| P62328 | TMSB4X | 4 | 4 | FALSE | FALSE |
| P67775;P62714;A0 | PPP2CA;PPP2CB | 4 | 4 | FALSE | FALSE |
| P62805 | HIST1H4A | 3 | 3 | FALSE | FALSE |
| P62937;F8WE65;C | PPIA | 7 | 7 | FALSE | FALSE |
| P63010;A0A087WU | AP2B1;AP1B1 | 5 | 5 | TRUE | FALSE |
| P63244;J3KPE3;D6 | GNB2L1 | 12 | 12 | FALSE | FALSE |
| P68032;P68133;A6 | ACTC1;ACTA1 | 14 | 14 | FALSE | FALSE |
| P68363;Q71U36;P( | TUBA1B;TUBA1A; | 16 | 16 | TRUE | FALSE |
| P68371;P04350 | TUBB4B;TUBB4A | 16 | 3 | FALSE | FALSE |
| P68871;F8W6P5;A | HBB;HBD | 4 | 3 | FALSE | FALSE |
| P81605 | DCD | 6 | 6 | FALSE | FALSE |
| P84095 | RHOG | 5 | 4 | TRUE | FALSE |

| | | | | | |
|---|---|---|---|---|---|
| P98175 | RBM10 | 4 | 4 | TRUE | FALSE |
| Q00534 | CDK6 | 5 | 4 | TRUE | FALSE |
| Q01469;I6L8B7 | FABP5 | 8 | 8 | FALSE | FALSE |
| Q01804 | OTUD4 | 4 | 4 | FALSE | FALSE |
| Q02413 | DSG1 | 27 | 27 | FALSE | FALSE |
| Q02878 | RPL6 | 9 | 9 | TRUE | FALSE |
| Q04759 | PRKCQ | 5 | 5 | TRUE | FALSE |
| Q04837;C9K0U8;A | SSBP1 | 2 | 2 | FALSE | FALSE |
| Q06203;D6RCC8 | PPAT | 8 | 8 | TRUE | FALSE |
| Q06830;A0A0A0M | PRDX1 | 12 | 12 | FALSE | FALSE |
| Q07021;I3L3Q7;I3I | C1QBP | 2 | 2 | FALSE | FALSE |
| Q07960;H0YE29 | ARHGAP1 | 10 | 10 | TRUE | FALSE |
| Q08211 | DHX9 | 7 | 7 | FALSE | FALSE |
| Q08554 | DSC1 | 21 | 21 | FALSE | FALSE |
| Q09028;H0YF10 | RBBP4 | 20 | 20 | TRUE | FALSE |
| Q12789 | GTF3C1 | 7 | 7 | FALSE | FALSE |
| Q12851;C9JCU6 | MAP4K2 | 8 | 8 | TRUE | FALSE |
| Q12874 | SF3A3 | 5 | 5 | FALSE | FALSE |
| Q13017 | ARHGAP5 | 6 | 6 | TRUE | FALSE |
| Q13137 | CALCOCO2 | 3 | 3 | FALSE | FALSE |
| Q13177 | PAK2 | 12 | 12 | TRUE | FALSE |
| Q13200 | PSMD2 | 9 | 9 | FALSE | FALSE |
| Q13263;M0R0K9 | TRIM28 | 9 | 9 | TRUE | FALSE |

| | | | | | |
|---|---|---|---|---|---|
| Q13363;H0Y8W7 | CTBP1 | 6 | 6 | FALSE | FALSE |
| Q13469 | NFATC2 | 16 | 16 | TRUE | FALSE |
| Q13526;K7EN45 | PIN1 | 3 | 3 | FALSE | FALSE |
| Q13541 | EIF4EBP1 | 3 | 3 | TRUE | FALSE |
| Q13542 | EIF4EBP2 | 4 | 4 | TRUE | FALSE |
| Q13547;Q5TEE2 | HDAC1 | 7 | 7 | TRUE | FALSE |
| Q13619;A0A0A0MI | CUL4A | 14 | 14 | TRUE | FALSE |
| Q13835 | PKP1 | 9 | 9 | FALSE | FALSE |
| Q13838;A0A0G2JJ | DDX39B | 9 | 9 | FALSE | FALSE |
| Q14008 | CKAP5 | 17 | 17 | TRUE | FALSE |
| Q14019;H3BT58 | COTL1 | 7 | 7 | FALSE | FALSE |
| Q14166 | TTLL12 | 8 | 8 | TRUE | FALSE |
| Q14186;F5H452;Q | TFDP1 | 2 | 2 | TRUE | FALSE |
| Q14289 | PTK2B | 6 | 6 | TRUE | FALSE |
| Q14318;U3KQI1;A( | FKBP8 | 3 | 3 | FALSE | FALSE |
| Q14566 | MCM6 | 10 | 10 | FALSE | FALSE |
| Q14574 | DSC3 | 11 | 11 | FALSE | FALSE |
| Q14694 | USP10 | 5 | 5 | TRUE | FALSE |
| Q14697;F5H6X6;E | GANAB | 13 | 13 | FALSE | FALSE |
| Q14847;C9J9W2 | LASP1 | 8 | 8 | TRUE | FALSE |
| Q15025;E7EMV7;A | TNIP1 | 3 | 3 | FALSE | FALSE |
| Q15029 | EFTUD2 | 6 | 6 | TRUE | FALSE |
| Q15046 | KARS | 3 | 3 | FALSE | FALSE |

| | | | | | |
|---|---|---|---|---|---|
| Q15084 | PDIA6 | 8 | 8 | FALSE | FALSE |
| Q15102;M0R389;M | PAFAH1B3 | 5 | 5 | TRUE | FALSE |
| Q15181;Q5SQT6 | PPA1 | 5 | 5 | FALSE | FALSE |
| Q15276 | RABEP1 | 5 | 5 | FALSE | FALSE |
| Q15366;H3BRU6;F | PCBP2 | 9 | 7 | TRUE | FALSE |
| Q15417;E9PDU6 | CNN3 | 7 | 6 | FALSE | FALSE |
| Q15459 | SF3A1 | 6 | 6 | TRUE | FALSE |
| Q15517 | CDSN | 7 | 7 | FALSE | FALSE |
| Q15645 | TRIP13 | 3 | 3 | FALSE | FALSE |
| Q15717;M0QZR9 | ELAVL1 | 6 | 6 | TRUE | FALSE |
| Q15758;M0QX44;N | SLC1A5 | 3 | 3 | TRUE | FALSE |
| Q15828 | CST6 | 3 | 3 | FALSE | FALSE |
| Q15843;F8VSA6;E | NEDD8;NEDD8-M | 3 | 3 | TRUE | FALSE |
| Q16566 | CAMK4 | 4 | 4 | FALSE | FALSE |
| Q16576;E9PC52;Q | RBBP7 | 16 | 12 | FALSE | FALSE |
| Q16610 | ECM1 | 7 | 7 | FALSE | FALSE |
| Q16891;B9A067;H | IMMT | 5 | 5 | FALSE | FALSE |
| Q2NKX8;B5MDQ0 | ERCC6L | 12 | 12 | TRUE | FALSE |
| Q32MZ4 | LRRFIP1 | 10 | 10 | FALSE | FALSE |
| Q4G0F5 | VPS26B | 5 | 5 | TRUE | FALSE |
| Q53QZ3 | ARHGAP15 | 2 | 2 | TRUE | FALSE |
| Q5JSH3;H7BY83 | WDR44 | 4 | 4 | FALSE | FALSE |
| Q5JSP0 | FGD3 | 5 | 5 | TRUE | FALSE |

| | | | | | |
|---|---|---|---|---|---|
| Q5JXI8;Q13642;Q5 | FHL1 | 7 | 7 | TRUE | FALSE |
| Q9HB21;Q5RGS4 | PLEKHA1 | 2 | 2 | FALSE | FALSE |
| Q5SW79;H0Y2V6 | CEP170 | 9 | 9 | FALSE | FALSE |
| Q5T5Y3 | CAMSAP1 | 9 | 9 | FALSE | FALSE |
| Q5T6F2;A0A8C8M( | UBAP2 | 14 | 14 | TRUE | FALSE |
| Q5T749 | KPRP | 23 | 23 | FALSE | FALSE |
| Q9Y5Q8;Q5T7U0;Q | GTF3C5 | 2 | 2 | TRUE | FALSE |
| Q9UGV2;Q5TH30;F | NDRG3 | 5 | 5 | TRUE | FALSE |
| Q5W0B1 | RNF219 | 4 | 4 | FALSE | FALSE |
| Q641Q2;Q9Y4E1;F | FAM21A;FAM21C | 10 | 10 | FALSE | FALSE |
| Q68DQ2 | CRYBG3 | 9 | 9 | TRUE | FALSE |
| Q6FI81;H3BV90;H3 | CIAPIN1 | 8 | 8 | TRUE | FALSE |
| Q6IA86 | ELP2 | 6 | 6 | FALSE | FALSE |
| Q6P1N0 | CC2D1A | 7 | 7 | FALSE | FALSE |
| Q6P4A8 | PLBD1 | 6 | 6 | FALSE | FALSE |
| Q6PJI9;B7Z7V6 | WDR59 | 5 | 5 | TRUE | FALSE |
| Q6UWP8 | SBSN | 20 | 20 | FALSE | FALSE |
| Q6XZF7;A0A1C7CY | DNMBP | 10 | 10 | FALSE | FALSE |
| Q6ZSZ5;A0A590UK | ARHGEF18 | 4 | 4 | TRUE | FALSE |
| Q7KZF4 | SND1 | 11 | 11 | TRUE | FALSE |
| Q7L523;Q5VZM2;Q | RRAGA;RRAGB | 6 | 6 | TRUE | FALSE |
| Q7L9L4;Q9H8S9;D | MOB1B;MOB1A | 4 | 4 | FALSE | FALSE |
| Q7Z5L9 | IRF2BP2 | 3 | 3 | FALSE | FALSE |

| | | | | | |
|---|---|---|---|---|---|
| Q7Z5W3 | BCDIN3D | 2 | 2 | FALSE | FALSE |
| Q86UW6;H0YA93 | N4BP2 | 14 | 14 | FALSE | FALSE |
| Q86VP6;A0A0C4D( | CAND1 | 12 | 12 | FALSE | FALSE |
| Q86WV1;J3QRG3 | SKAP1 | 4 | 4 | TRUE | FALSE |
| Q8IUI8 | CRLF3 | 4 | 4 | TRUE | FALSE |
| Q8IW75 | SERPINA12 | 11 | 11 | FALSE | FALSE |
| Q8IWR0;I3L2K5 | ZC3H7A | 9 | 9 | FALSE | FALSE |
| Q8N0X7 | SPG20 | 2 | 2 | TRUE | FALSE |
| Q8N5C8;F6SS63 | TAB3 | 6 | 6 | FALSE | FALSE |
| Q8NBF2 | NHLRC2 | 6 | 6 | FALSE | FALSE |
| Q8ND24;E9PN76 | RNF214 | 4 | 4 | FALSE | FALSE |
| Q8NG31;H0YCZ2 | CASC5 | 13 | 13 | FALSE | FALSE |
| Q8TB72;C9JW01;A | PUM2 | 5 | 5 | TRUE | FALSE |
| Q8TD16 | BICD2 | 26 | 26 | TRUE | FALSE |
| Q8TDL5 | BPIFB1 | 3 | 3 | FALSE | FALSE |
| Q8TDX7 | NEK7 | 3 | 3 | TRUE | FALSE |
| Q8WUF5 | PPP1R13L | 10 | 10 | FALSE | FALSE |
| Q8WVJ2;E5RFP0 | NUDCD2 | 4 | 4 | TRUE | FALSE |
| Q8WVV4 | POF1B | 11 | 11 | FALSE | FALSE |
| Q8WW12 | PCNP | 3 | 3 | TRUE | FALSE |
| Q8WX93 | PALLD | 11 | 11 | FALSE | FALSE |
| Q8WZA0 | LZIC | 7 | 7 | TRUE | FALSE |
| Q92598;A0A0A0M! | HSPH1 | 6 | 4 | TRUE | FALSE |

| | | | | | |
|---|---|---|---|---|---|
| Q92608;H0YB76;E | DOCK2 | 9 | 9 | TRUE | FALSE |
| Q92609;C9J3F6 | TBC1D5 | 12 | 12 | TRUE | FALSE |
| Q92636 | NSMAF | 3 | 3 | FALSE | FALSE |
| Q92820;A0A7I2YQ | GGH | 7 | 7 | FALSE | FALSE |
| Q92844;E7EQA9;E | TANK | 6 | 6 | FALSE | FALSE |
| Q93052;A0A087W | LPP | 16 | 16 | TRUE | FALSE |
| Q93084 | ATP2A3 | 5 | 4 | FALSE | FALSE |
| Q96A49 | SYAP1 | 4 | 4 | FALSE | FALSE |
| Q96AP4;X6R6X3 | ZUFSP | 12 | 12 | TRUE | FALSE |
| Q96C19;H0Y4Y4;Q | EFHD2;EFHD1 | 4 | 4 | TRUE | FALSE |
| Q96C86;A0A3B3IT | DCPS | 5 | 5 | FALSE | FALSE |
| Q96D71;E9PMG1;I | REPS1 | 4 | 4 | TRUE | FALSE |
| Q96F86 | EDC3 | 5 | 5 | TRUE | FALSE |
| Q96GA3 | LTV1 | 3 | 3 | FALSE | FALSE |
| Q96H79 | ZC3HAV1L | 3 | 3 | FALSE | FALSE |
| Q96HS1;F5GXG4 | PGAM5 | 11 | 11 | FALSE | FALSE |
| Q96I24 | FUBP3 | 7 | 7 | TRUE | FALSE |
| Q96J01;D6REC9 | THOC3 | 3 | 3 | TRUE | FALSE |
| Q96JQ2 | CLMN | 6 | 6 | FALSE | FALSE |
| Q96P63 | SERPINB12 | 14 | 14 | FALSE | FALSE |
| Q96RQ3;E9PHF7;E | MCCC1 | 23 | 23 | FALSE | FALSE |
| Q96S59 | RANBP9 | 8 | 8 | TRUE | FALSE |
| Q96T21 | SECISBP2 | 5 | 5 | TRUE | FALSE |

| | | | | | |
|---|---|---|---|---|---|
| Q99497;K7EN27;K | PARK7 | 4 | 4 | FALSE | FALSE |
| Q99538;G3V4E4 | LGMN | 3 | 3 | TRUE | FALSE |
| Q99614;A0A804HI | TTC1 | 14 | 14 | TRUE | FALSE |
| Q99832 | CCT7 | 8 | 8 | FALSE | FALSE |
| Q9BQE3;F5H5D3 | TUBA1C | 16 | 2 | FALSE | FALSE |
| Q9BRG1;K7EKV4;K | VPS25 | 6 | 6 | TRUE | FALSE |
| Q9BRP1 | PDCD2L | 5 | 5 | TRUE | FALSE |
| Q9BU76 | MMTAG2 | 2 | 2 | FALSE | FALSE |
| Q9BV57;A0A0G2JQ | ADI1 | 4 | 4 | FALSE | FALSE |
| Q9BXB4 | OSBPL11 | 3 | 3 | FALSE | FALSE |
| Q9BXW9;A0A6Q8P | FANCD2 | 5 | 5 | FALSE | FALSE |
| Q9BYJ9 | YTHDF1 | 9 | 6 | FALSE | FALSE |
| Q9C0B0;K7EQ93 | UNK | 3 | 3 | FALSE | FALSE |
| Q9C0F1;D6RGX6 | CEP44 | 2 | 2 | TRUE | FALSE |
| Q9GZT9 | EGLN1 | 2 | 2 | FALSE | FALSE |
| Q9GZZ8;F8W0V3 | LACRT | 2 | 2 | FALSE | FALSE |
| Q9H0U4;E9PLD0;C | RAB1B;RAB1C | 4 | 4 | FALSE | FALSE |
| Q9H1A4;H0Y564 | ANAPC1 | 3 | 3 | FALSE | FALSE |
| Q9H1E1 | RNASE7 | 2 | 2 | FALSE | FALSE |
| Q9H2H8;B8ZZ77;H | PPIL3 | 5 | 5 | TRUE | FALSE |
| Q9H3P7 | ACBD3 | 3 | 3 | FALSE | FALSE |
| Q9H425;E5RFY9 | C1orf198 | 4 | 4 | TRUE | FALSE |
| Q9H6R7 | C2orf44 | 4 | 4 | FALSE | FALSE |

| | | | | | |
|---|---|---|---|---|---|
| Q9H9B4;D6RDG7;S | SFXN1 | 3 | 3 | FALSE | FALSE |
| Q9HB07;F8VQQ3 | C12orf10 | 5 | 5 | FALSE | FALSE |
| Q9HB90 | RRAGC | 3 | 3 | TRUE | FALSE |
| Q9HC38;I3L3Q4 | GLOD4 | 4 | 4 | TRUE | FALSE |
| Q9HCN4 | GPN1 | 3 | 3 | FALSE | FALSE |
| Q9HCY8 | S100A14 | 3 | 3 | FALSE | FALSE |
| Q9HD45;Q5TB53 | TM9SF3 | 4 | 4 | FALSE | FALSE |
| Q9HD89 | RETN | 3 | 3 | FALSE | FALSE |
| Q9NP72;A0A804HI | RAB18 | 4 | 4 | FALSE | FALSE |
| Q9NPI6 | DCP1A | 9 | 9 | TRUE | FALSE |
| Q9NPJ6;Q5T911 | MED4 | 2 | 2 | FALSE | FALSE |
| Q9NQ38;E5RFU9;E | SPINK5 | 3 | 3 | FALSE | FALSE |
| Q9NQC3 | RTN4 | 13 | 13 | TRUE | FALSE |
| Q9NR45 | NANS | 3 | 3 | TRUE | FALSE |
| Q9NUQ3 | TXLNG | 11 | 11 | TRUE | FALSE |
| Q9NVI7;Q5T9A4;Q | ATAD3A;ATAD3B; | 6 | 6 | FALSE | FALSE |
| Q9NWS0;M0QYA2; | PIH1D1 | 11 | 11 | TRUE | FALSE |
| Q9NXC5 | MIOS | 12 | 12 | FALSE | FALSE |
| Q9NYJ8;A0A1B0GV | TAB2 | 5 | 5 | FALSE | FALSE |
| Q9NYL9;H0YNJ8;H | TMOD3 | 11 | 11 | TRUE | FALSE |
| Q9NZ09;A0A0D9S( | UBAP1 | 4 | 4 | TRUE | FALSE |
| Q9NZH8 | IL36G | 6 | 6 | FALSE | FALSE |
| Q9NZM3 | ITSN2 | 5 | 5 | FALSE | FALSE |

| | | | | | |
|---|---|---|---|---|---|
| Q9NZT1 | CALML5 | 9 | 9 | FALSE | FALSE |
| Q9P035;H3BPZ1;H | HACD3 | 5 | 5 | FALSE | FALSE |
| Q9P0L0 | VAPA | 5 | 5 | FALSE | FALSE |
| Q9P258 | RCC2 | 8 | 8 | TRUE | FALSE |
| Q9P287 | BCCIP | 2 | 2 | TRUE | FALSE |
| Q9P2M7 | CGN | 10 | 10 | FALSE | FALSE |
| Q9UBB9 | TFIP11 | 6 | 6 | TRUE | FALSE |
| Q9UEE9 | CFDP1 | 5 | 5 | FALSE | FALSE |
| Q9UHD1 | CHORDC1 | 7 | 7 | FALSE | FALSE |
| Q9UHD8 | SEPT9 | 15 | 15 | TRUE | FALSE |
| Q9UIM3 | FKBPL | 4 | 4 | TRUE | FALSE |
| Q9UJ70;C9JEV6;H7 | NAGK | 6 | 6 | TRUE | FALSE |
| Q9UJU6;B4DDD6;F | DBNL | 5 | 5 | TRUE | FALSE |
| Q9UK76;J3KT51 | HN1 | 5 | 5 | TRUE | FALSE |
| Q9ULM3 | YEATS2 | 10 | 10 | TRUE | FALSE |
| Q9UMS4;F5GY56;H | PRPF19 | 4 | 4 | FALSE | FALSE |
| Q9UQ80;F8VR77 | PA2G4 | 6 | 6 | FALSE | FALSE |
| Q9Y224;G3V4C6 | C14orf166 | 4 | 4 | TRUE | FALSE |
| Q9Y228;B1AJU2;E2 | TRAF3IP3 | 8 | 8 | TRUE | FALSE |
| Q9Y263;H0YBW4;E | PLAA | 3 | 3 | TRUE | FALSE |
| Q9Y2D5;C9JVY5 | AKAP2 | 6 | 6 | TRUE | FALSE |
| Q9Y2I7;E9PDH4 | PIKFYVE | 10 | 10 | FALSE | FALSE |
| Q9Y2V2;H3BSW7;H | CARHSP1 | 5 | 5 | FALSE | FALSE |

| | | | | | |
|---|---|---|---|---|---|
| Q9Y337 | KLK5 | 3 | 3 | FALSE | FALSE |
| Q9Y3F4 | STRAP | 8 | 8 | FALSE | FALSE |
| Q9Y3I0 | RTCB | 5 | 5 | TRUE | FALSE |
| Q9Y478 | PRKAB1 | 7 | 7 | TRUE | FALSE |
| Q9Y512 | SAMM50 | 2 | 2 | TRUE | FALSE |
| Q9Y5S9 | RBM8A | 5 | 5 | TRUE | FALSE |
| Q9Y613 | FHOD1 | 5 | 5 | FALSE | FALSE |
| Q9Y6A5;A0A494BZ | TACC3 | 8 | 8 | TRUE | FALSE |
| S4R3N1;Q9Y3A3 | HSPE1-MOB4;MC | 4 | 4 | TRUE | FALSE |
